# Genomics of a complete butterfly continent

**DOI:** 10.1101/829887

**Authors:** Jing Zhang, Qian Cong, Jinhui Shen, Paul A. Opler, Nick V. Grishin

**Affiliations:** Howard Hughes Medical Institute, University of Texas Southwestern Medical Center, Dallas, TX, 75390, USA; Departments of Biophysics and Biochemistry, University of Texas Southwestern Medical Center, Dallas, TX, 75390, USA; Institute for Protein Design and Department of Biochemistry, University of Washington, Seattle, WA, 98195, USA; Department of Bioagricultural Sciences and Pest Management, Colorado State University, Fort Collins, CO, 80523, USA

## Abstract

Never before have we had the luxury of choosing a continent, picking a large phylogenetic group of animals, and obtaining genomic data for its every species. Here, we sequence all 845 species of butterflies recorded from North America north of Mexico. Our comprehensive approach reveals the pattern of diversification and adaptation occurring in this phylogenetic lineage as it has spread over the continent, which cannot be seen on a sample of selected species. We observe bursts of diversification that generated taxonomic ranks: subfamily, tribe, subtribe, genus, and species. The older burst around 70 Mya resulted in the butterfly subfamilies, with the major evolutionary inventions being unique phenotypic traits shaped by high positive selection and gene duplications. The recent burst around 5 Mya is caused by explosive radiation in diverse butterfly groups associated with diversification in transcription and mRNA regulation, morphogenesis, and mate selection. Rapid radiation correlates with more frequent introgression of speciation-promoting and beneficial genes among radiating species. Radiation and extinction patterns over the last 100 million years suggest the following general model of animal evolution. A population spreads over the land, adapts to various conditions through mutations, and diversifies into several species. Occasional hybridization between these species results in accumulation of beneficial alleles in one, which eventually survives, while others become extinct. Not only butterflies, but also the hominids may have followed this path.

---

Butterflies are among the most beloved animals, beautiful and harmless, they have attracted human attention since prehistoric times (*1*). Being one of the best-studied insects phenotypically (*2*), butterflies remain largely unexplored by genomics. Until recently, genomic studies of butterflies were confined to a couple of model organisms and pests, such as *Heliconius*, monarch, and cabbage white, with initial studies leading to groundbreaking insights into mimicry, migration, and toxin resistance (*3-5*). We have been expanding these efforts on butterfly genomics to cover a broader range of species (*6-11*). With the rapid decrease in the price of DNA sequencing and the constant development of analytical methods, the time is ripe to sequence the genomes of all butterfly species over a continent.

The diversity of butterflies, which form a clade within moths (*12*), is captured in 7 families worldwide (*13*). Six of these families are represented in North America north of Mexico, and the butterfly fauna of this region is well-documented (*2, 14, 15*). Here, we obtain and analyze the genomes of all 845 (Table S1) butterfly species in the United States and Canada (USC). A number of these species are of conservation concern, including 25 endangered and threatened taxa (*16*). The new genomic datasets comprehensively covering USC butterflies reveal the detailed history of their speciation and adaptation and suggest the genetic basis of their unique phenotypic traits. As a result, we find a bewildering pattern of phylogenetic diversification that we rationalize in a general model of animal evolution reaching beyond butterflies and insects. Moreover, recently developed analytical methods have demonstrated the power of extracting information from thick protein sequence alignments to accurately model spatial structures (*17*), screen interacting partners (*18*) and predict functions (*19*) of proteins. The diverse datasets of protein sequences we have obtained will allow structure and function prediction for many eukaryotic gene products, enabling future discoveries.

## Reference genomes of butterflies

We sequenced and annotated 23 reference genomes of butterflies from the United States. Combined with the 13 genomes published previously (*3-11, 20-23*) (8 by our group), a total of 36 reference genomes are used in this study. The N50 of the new genomic assemblies ranges from 50 to 3,700 kb, and they are over 95% complete in essential genes (*24*) (Table 1). While the size of these genomes is drastically different, between 217 and 1,040 Mb, they encode comparable numbers of proteins, about 15,000. In contrast, the fraction of repeats correlates strongly with the genome size (Pearson’s correlation coefficient > 0.7), suggesting that the genome size in butterflies varies due to repetitive and transposable elements. Genome size does not conform with butterfly phylogeny and can differ even for close relatives, likely due to the activity of transposons.

**Table 1.**
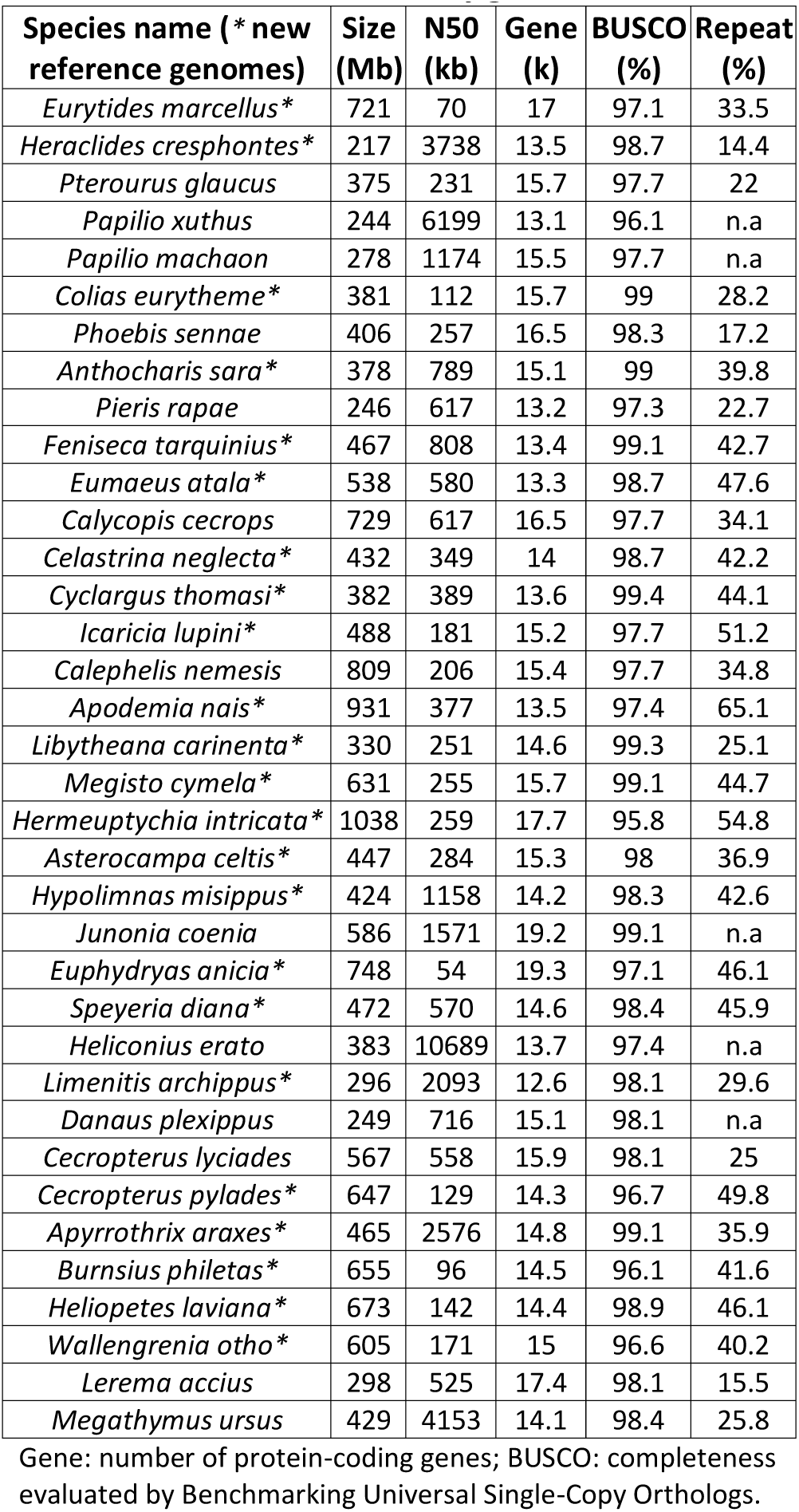
Statistics for butterfly genome assemblies.

The reference genomes were selected to cover diverse phylogenetic groups of the USC butterflies, allowing us to carry out genomic comparisons across the phylogeny. In butterfly genomes, we detected 530 Ultra-Conserved genomic Elements (UCE, Table S2). Similarly to mammals (*25, 26*), UCEs in butterflies mostly reside in the intergenic regions (Fig. 1A) and around the genes functioning in transcription regulation and developmental processes (Fig. 1B and Table S3). The UCEs constitute merely 0.01% of the genomes, and most other regions have diverged rapidly among butterflies. Only 7–14% of the genomic sequence can be confidently aligned between species from different families (*27*), and the remainder contains variable and repetitive regions. The observed tolerance to transposon activity in butterflies may be adaptive, allowing them to exploit the retrotransposition mechanism for gene duplication and expansion (*28*). We identified 8,581 orthologous gene groups present in at least 75% of the reference genomes, and each species experienced gene duplications in 2–9% of these groups. In addition to the high sequence divergence, frequent gene duplication may be another reason for phenotypic diversity and adaptation.

**Fig. 1.**
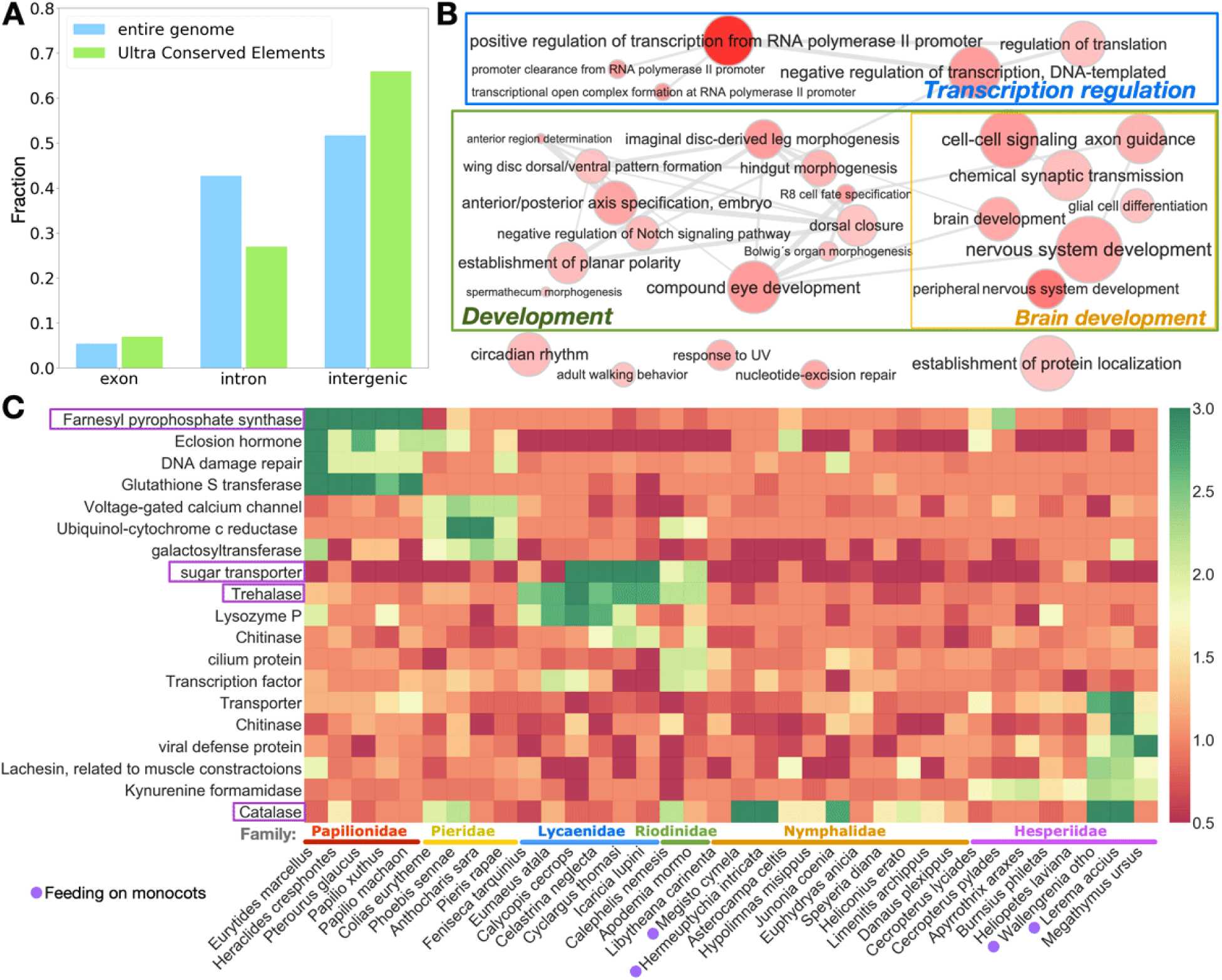
Ultra-Conserved genomic Elements (UCE) in butterfly genomes and lineage-specific gene expansion. (A) Distribution of UCEs across different types of regions in the *Heliconius erato* genome. (B) Enriched Gene Ontology (GO) terms associated with genes near (within 10 kb) UCEs in the genome visualized using REVIGO (*29*). Size of dots reflects the number of genes associated with this GO term in *Drosophila melanogaster*. Color of dots indicates the statistical significance, with darker color corresponding to lower P-values. Connections between dots indicate similarity in the meaning of the GO terms. (C) The most prominent lineage-specific gene expansions. The total length of proteins in an orthologous group is calculated for each species to obtain the median over all species, than the total length for each species is normalized by this median. The normalized total length is shown as heatmap. Each cell is colored from red through yellow to green for values from 0.5 to 3.0. Values below 0.5 are colored in the same color as 0.5 and values above 3.0 are colored as 3.0.

We systematically catalogued lineage-specific gene expansions (Table S4) in reference genomes, and a number of prominent examples are given in Fig. 1C. Many of the gene expansions occur in protein families involved in acquiring nutrients from food and resisting toxins and pathogens, but some (magenta boxes in Fig. 1C) may explain the lineage-specific phenotypes. We reported an expansion of farnesyl pyrophosphate synthase (FPPS) homologs in *Pterourus glaucus* (*11*), and now we find this expansion present in all sequenced Papilionidae species. FPPSs function in steroid and terpene synthesis (*30*), and the Papilionidae-specific organ, osmeterium, secrets terpenes to repel birds (*31*). Therefore, the unique expansion and diversification of FPPSs may produce the bouquet of terpenes aiding caterpillars’ arms races with predators. Other notable gene expansions involve sugar transporters and trehalases in Lycaenidae. The caterpillars of Lycaenidae secrete nectar-like liquids as a “reward” to trick ants into protecting and even feeding the caterpillars (*32*). Some Riodinidae species are also associated with ants and possess similar gene expansions. The additional sugar transporters may play a role in secreting sugars that are being fed to ants, while trehalase may convert trehalose from food plant to sweeter-tasting molecules, contributing to this adaptation.

Furthermore, some gene expansions may underlie convergence in phenotypes, such as the expansion of catalases in the two distant phylogenetic lineages—the satyrs (Satyrini) and grass-skippers (Hesperiinae)—whose caterpillars converged to feeding on monocots. The catalases decompose hydrogen peroxide and thus protect against oxidative damage (*33*). Feeding on nutrient-deficient grasses, sedges and palms (*34*) leads to a prolonged caterpillar stage and may increase the likelihood of oxidative damage that would be mitigated by additional catalases. Apparently switching to monocot feeding was an evolutionarily successful innovation that resulted in explosive speciation in both satyrs and grass-skippers (*35*). They became the most species-rich phylogenetic groups among American butterflies, and their parallel diversifications of catalases are intriguing.

## Phylogeny of USC butterflies

We obtained whole genome shotgun sequences of all 845 butterfly species recorded from the United States and Canada (*15*). Phylogenetic trees constructed from protein-coding genes in the nuclear genome (10,000–15,000 kb positions), Z chromosome (360**–**641 kb positions) and mitochondrial genome (11 kb positions) are largely consistent with each other (Fig. 2 and Fig. S1) and support established views about the deeper phylogeny of butterflies (family and subfamily). Namely, swallowtails (family Papilionidae) are sister to all other butterflies (*36*), and the topology of other butterfly families agrees with previous studies (*13*). Genetic divergence between gossamer-winged butterflies (family Lycaenidae) and metalmarks (family Riodinidae) is smaller than that between some subfamilies of brush-footed butterflies (Nymphalidae). Thus, it may be best to view metalmarks as one of the subfamilies of Lycaenidae as discussed previously (*13*).

**Fig. 2.**
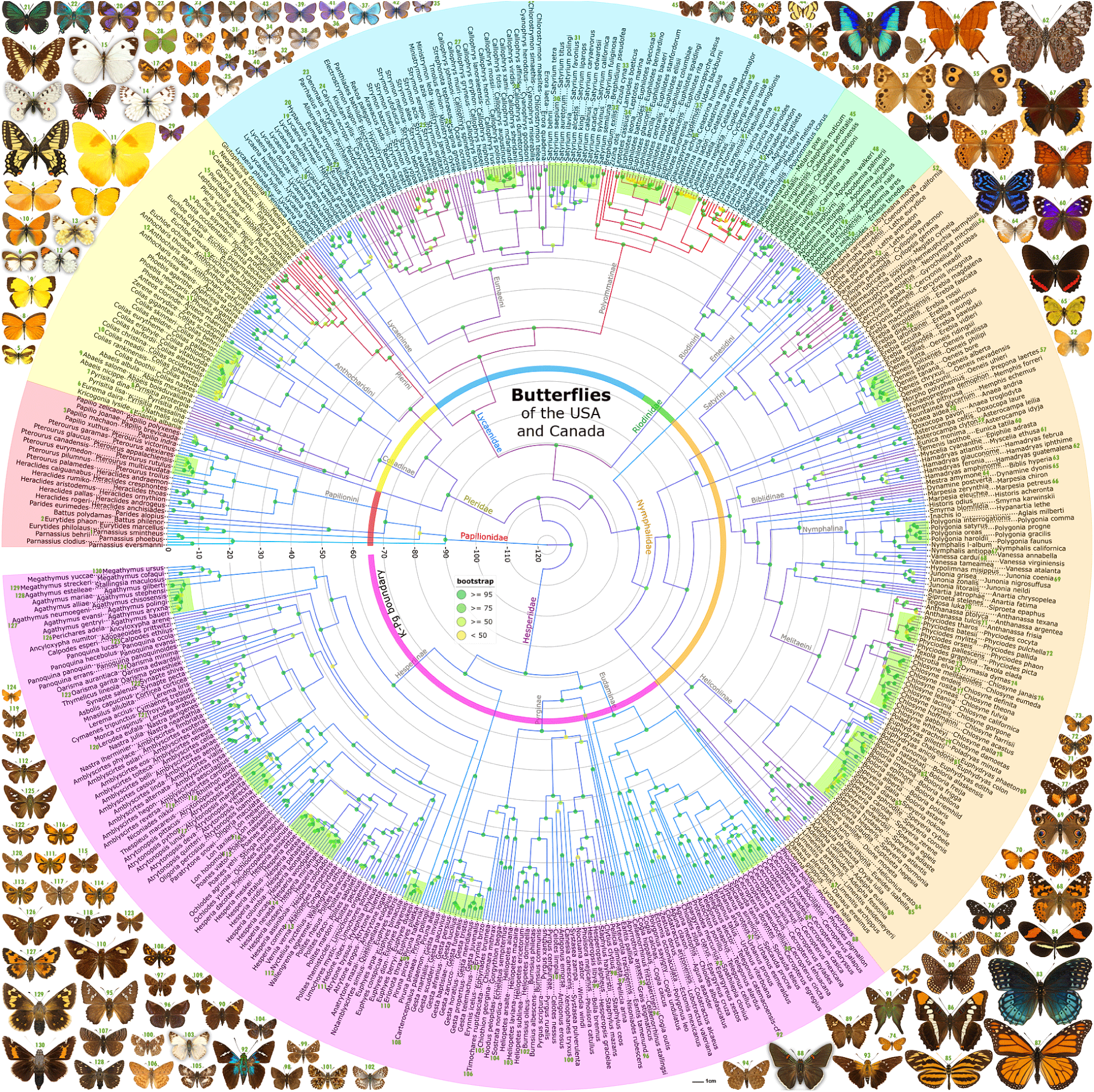
Time-calibrated phylogenetic tree of USC butterflies constructed from all nuclear genes. Tree branches are colored by relative substitution rate from cyan (slow evolving) through dark-blue and magenta to red (fast evolving). The time scale is in Mya. Species are arranged clockwise starting from the time-scale. A ring colored by butterfly family sectors is placed at the Cretaceous–Paleogene (K-Pg) boundary. Bootstrap support values are marked as color-coded dots on the tree nodes. Clusters of explosive radiation are highlighted in lime-green. The names of families and major phylogenetic lineages discussed in the text are given by their branches. Species names are highlighted by family and shown in two layers. Names in the inner layer are connected to the corresponding tree leaves with dots, and they label every other leaf in the tree. Each name in the outer layer is placed in between and connected by dots to two names of the inner layer, and it refers to the leaf in between the leaves indicated by these two names of the inner layer. Butterfly images represent all major phylogenetic lineages and numbers in green font associate specimens (a number is mostly above a specimen) with their names in the tree (numbers are either before or after the species names for the left and right halves of the tree, respectively).

However, our genome-level analysis revealed a number of problems with the current butterfly classification at a shallower phylogeny level (tribe and genus). We rectified these problems in dedicated publications (*37, 38*). Briefly, we proposed 6 new genera, 2 new subgenera, and reclassified 40 species (Table S1). Thus, the names of 6% USC butterflies were changed due to this expanded examination of genomes. In accordance with our previous findings (*39*), we stumble upon additional unexpected cases of rapid divergence and mimicry in wing patterns and shapes, where butterflies do not look similar to their close relatives, but resemble more distant species (Table S5). These insights illustrate the power of genomics in reshaping our knowledge of taxonomy and phylogeny of life.

Comparison of the three trees (autosome, Z chromosome, and mitogenome: Fig. S1) reveals confident incongruence between them. The incongruence is rampant among close relatives, and is present in almost every large genus. These inconsistencies likely reflect alternative evolutionary paths taken by different genomic segments of the same organism as a result of incomplete lineage sorting, introgression and hybridization (*3, 40, 41*). Generally, the nuclear trees correlate better with phenotypes than the mitogenome tree. In the genera that have experienced rapid radiation, such as *Colias, Euphilotes* and *Speyeria*, mitochondrial phylogeny is semi-random compared to nuclear phylogeny and phenotypes, and all 9 *Celastrina* species carry essentially identical mitochondrial DNA. Incongruence between the trees constructed from autosomes and the Z chromosome (Table S6) can originate in cases with extensive introgression (*40*). Sex chromosome-linked genes are shown to resist introgression in multiple species (*42-44*), and thus the Z chromosome tree may better reflect the history of speciation, not the averaged history of introgression. For instance, in accord with morphology, the Z chromosome suggests a sister relationship between morphologically similar but allopatric species *Junonia coenia* and *Junonia grisea* (*45, 46*). In contrast, the autosomal tree groups morphologically different but sympatric *Junonia grisea* and *Junonia nigrosuffusa*, who experience frequent hybridization and introgression.

## Diversification, extinction and bursts in radiation

We analyzed patterns of diversification in the time-calibrated phylogenetic tree of USC butterflies constructed from all protein-coding genes. The number of species from currently non-extinct lineages at each time point in the past is shown in Fig. 3A. This curve reflects both speciation and extinction, and is similar to exponential, but exhibits a decreased diversification rate in the last 2 million years (Myr). This apparent decrease is due to both the variation between individuals of a species (i.e., terminal branches lead to individuals, not species) and incomplete speciation events: some populations are on the way to become distinct species, but are not recognized as such today.

**Fig. 3.**
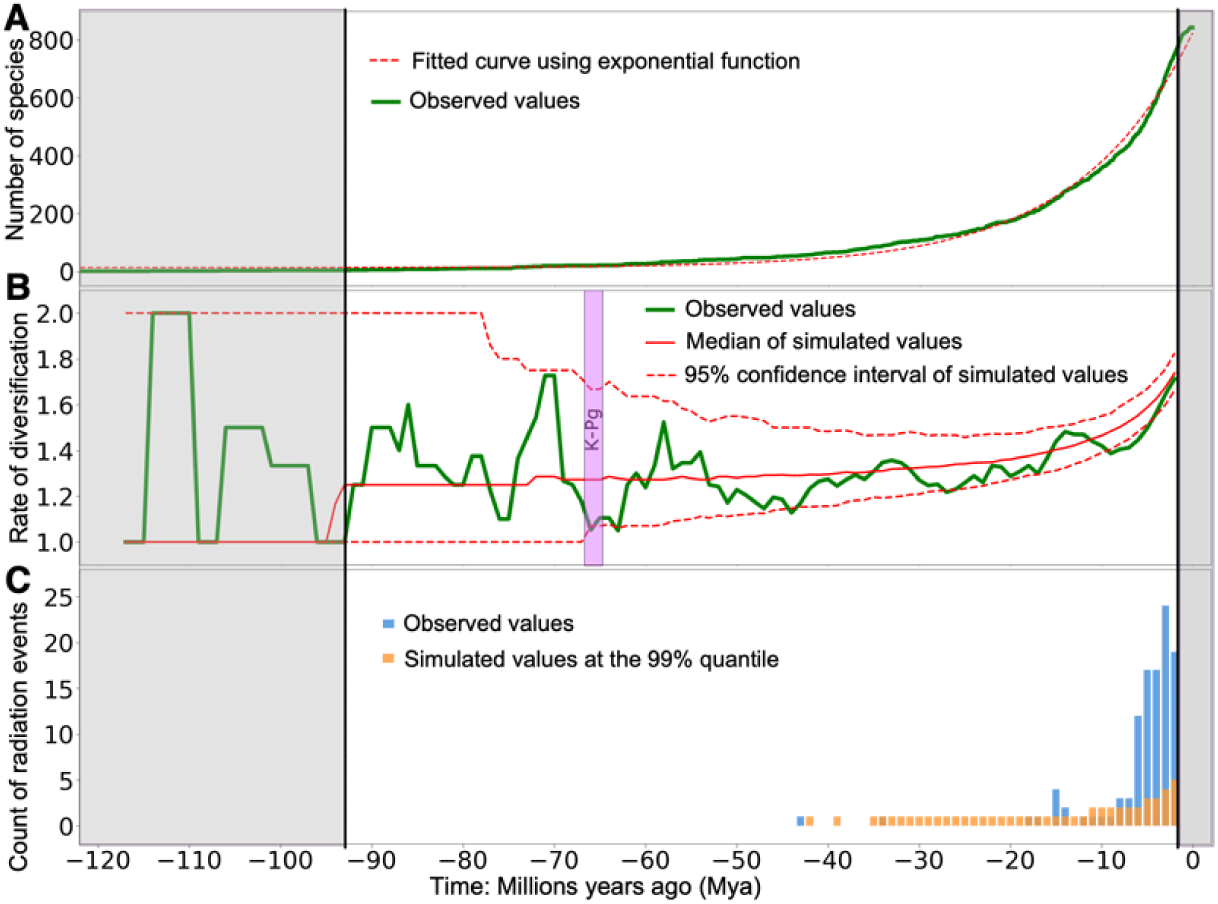
Uneven evolutionary rates and time progression of diversification in USC butterflies. (A) Growth in the number of non-extinct species over time. (B) Rate of diversification defined as the number of species at each time point divided by the number of species 5 Mya. The green curve shows observed data, and the red curves indicate simulations under constant speciation (0.15 per Myr) and extinction (0.08 per Myr) rates. The Cretaceous– Paleogene (K-Pg) boundary is marked as a purple bar. (C) Significantly more radiation events (i.e., a lineage splits into > 3 in < 2 Myr) are observed among USC butterflies (blue) in the last 8 Myr than in simulations (orange) that assume independence between speciation events.

Excluding the last 2 Myr, we fitted the diversification rate to a model with constant speciation and extinction rates, yielding estimated speciation rate of 0.15 per Myr and extinction rate of 0.08 per Myr (Fig. 3B). We exclude time points before 94 million years ago (Mya) because small number (≤ 5) of lineages is prone to random fluctuations. Both simulated and observed data show an increase in rate when approaching the present time. This increase is caused by species present at an earlier time point (but extinct by now) not being counted, leading to underestimation of the number of species that existed in the past (*47*). The observed species diversification rate (per 5 Myr) shows larger fluctuations than simulations, and these fluctuations are biologically meaningful. For instance, the minimum around 63–67 Mya reflects the Cretaceous–Paleogene extinction dated to 66 Mya (*48*), thus indirectly supporting the time-scale of the tree. Interestingly, the maxima around 90, 70, 60, 35, and 15 Mya (Fig. 3B) approximate the origins of clades corresponding to major levels in the taxonomic hierarchy: family, subfamily, tribe, subtribe, and genus, respectively. Starting from the diversification into subfamilies 70 Mya, which is the global maximum (94 Mya till now), there are 4 major peaks in the curve, as there are taxonomic levels. Discreteness of these levels may thus be a consequence of rapid speciation across phylogenetic lineages followed by extinctions that break the continuity of animal forms, leading to survival of only a few distant ones.

The latest increase in the diversification rate since 8 Mya (Fig. 3B) is the origin of species, and it brings a surprise. Inspection of the tree reveals many recent bursts of radiation, i.e., rapid diversification in some lineages leading to the origin of many species around the same time. To quantify this effect, we studied the time progression of the number of nodes in the tree that produce at least 4 branches within 2 Myr (Fig. 3C). Observed data (blue) show profoundly more radiation events in the recent past than simulations that assume an equal chance of speciation in every lineage (orange). These recent radiations recur across the tree of butterflies and were investigated in detail.

## Introgression of speciation-associated genes from distant relatives leads to radiation

We identified 18 clusters of species as undergoing explosive radiation (Table S7). These clusters belong to genera from four of the largest butterfly families. We looked for genes that diverge rapidly (P-value < 0.05) among species in each radiating cluster, but evolve relatively slowly in closely related non-radiating species. We found 273–846 such genes in each cluster, about 4% of all genes. A significant overlap in these genes exists among 18 radiating clusters with 430 common genes recurrently showing elevated divergence during radiation (P-value < 0.05, Table S8). The top 21 genes rapidly diverging in over a third of radiation clusters are shown in Fig. 4A. Proteins encoded by these 21 genes mostly belong to 4 major functional categories (green cells in Fig. 4A).

**Fig. 4.**
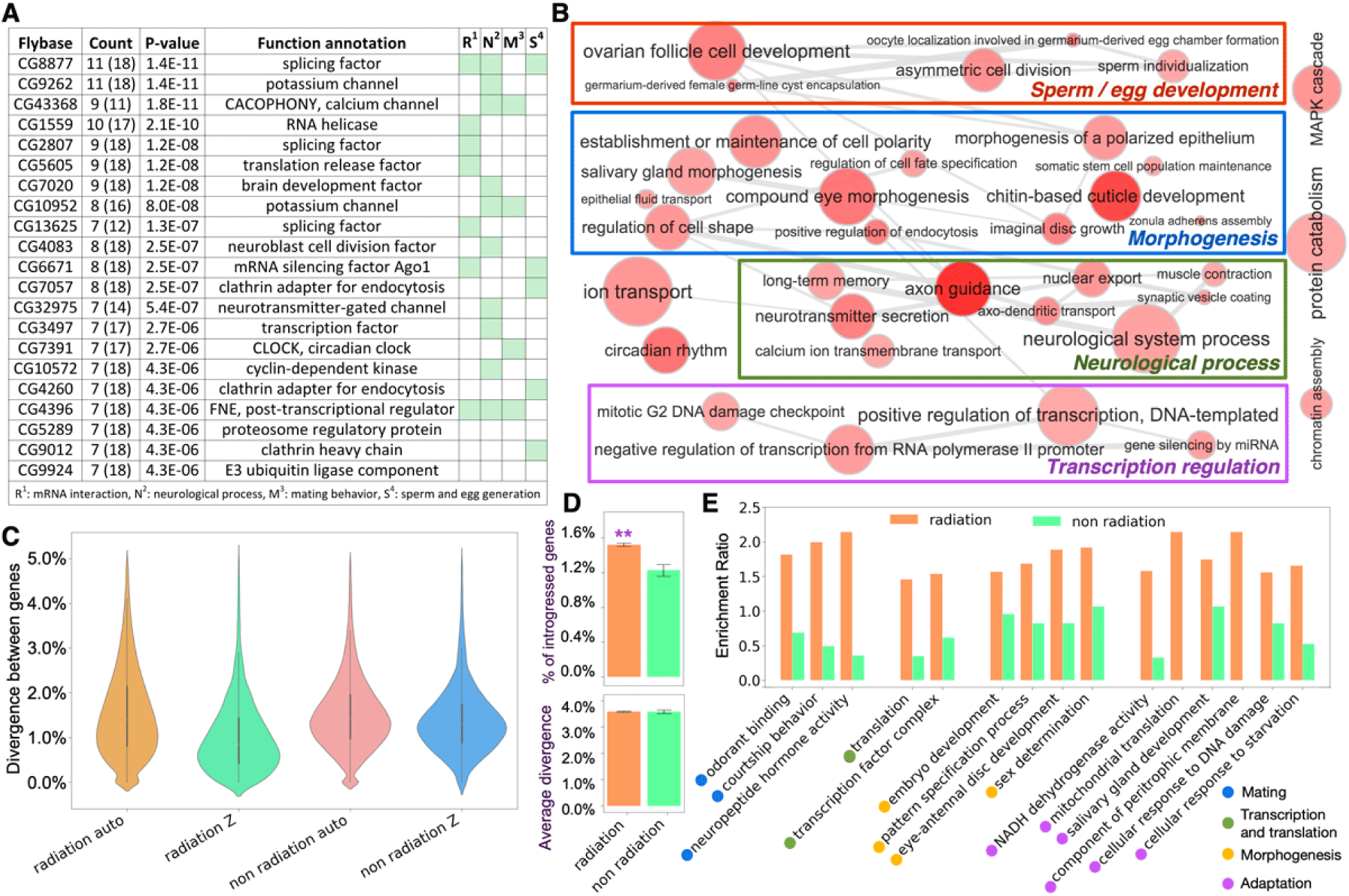
Radiation-associated proteins and the role of introgression in radiation. (A) Prominent radiation-associated proteins that are recurring in the largest number of radiation events. We define radiation-associated proteins as those that tend to show significantly (P-value < 0.01) elevated divergence in a radiating genus. (B) Enriched Gene Ontology (GO) terms associated with recurrent radiation-associated proteins. Size and color of dots reflect the number of genes associated with this term and the statistical significance (darker color = lower P-value) for this GO term’s enrichment, respectively. (C) Violin plots showing the distribution of sequence divergence of autosomal and Z-linked genes between sister species in radiating and non-radiating genera. (D) Distant, non-sister species from radiating genera exchange genes with each other significantly more frequently than those from non-radiating genera. (E) GO terms that are significantly enriched among genes that tend to introgress among distant species in radiating genera. Enrichment ratio is the probability for a GO term to be associated with a frequently introgressed gene divided by the chance for it to be associated with any gene. Orange and green bars show the enrichment for a GO term (annotation) in radiating and non-radiating genera, and the lack of green bars indicates a ratio of 0. Dots by the annotations mark the category of each GO term, and these categories are labeled in the right bottom corner.

First, 7 out of the 21 most frequent radiation-associated proteins are associated with splicing and silencing of mRNA. This observation echoes the studies of radiation in cichlids (*49*), suggesting that the increased complexity of mRNA regulation and maturation may be a general mechanism to rapidly generate divergence in animals bypassing the need of extensive variations in gene sequences. Second, 4 of the 21 proteins are directly related to mating (Fig. 4A). For instance, CACOPHONY is a calcium channel that senses species-specific mating song in *Drosophila* (*50*), and CLOCK is a transcription factor that regulates mating time (*51, 52*). Elevated divergence in such genes may directly alter mating behavior of butterflies, contribute to prezygotic isolation, and accelerate speciation. Finally, proteins involved in sperm and egg generation and neurological processes stand out not only among the 21 most frequent players (Fig. 4A), but also in all 430 recurrent radiation-associated proteins. Additionally, enrichment analysis of gene ontology (GO) terms associated with these 430 proteins reveals transcription regulation and morphogenesis as major functional categories for radiation-associated genes (Fig. 4B, Table S9). Differences in transcriptional factors are likely associated with divergence in DNA regulatory elements they bind to, and the latter has been shown to play an important role in *Drosophila* speciation (*53, 54*).

To understand why radiation-associated proteins exhibit elevated divergence among species in a radiating lineage, we first studied whether they are positively selected. Unexpectedly, radiation genes are characterized by a lower nonsynonymous substitution rate compared to other genes in all radiating genera but *Phyciodes* (Table S10). Thus, they are not undergoing stronger positive selection on their individual mutations than other genes. Instead, we find that radiation genes tend to introgress between distantly, but not closely, related species (P-value < 4.9e-12, see below). We hypothesize that introgression of speciation-promoting genes from more distant relatives is a mechanism that speeds up speciation of close relatives, causing explosive radiation.

To test this hypothesis, we compared the 18 radiating genera with others. We analyzed 63 pairs of sister species from radiating clusters and 68 pairs from non-radiating lineages (Table S11), and compared the distribution of sequence divergence of individual genes in autosomes and Z chromosome (Fig. 4C). Species pairs from radiating clusters show lower divergence in Z chromosome than in autosomes (green vs. orange in Fig. 4C), while non-radiating clusters do not display such a trend. Sex chromosome’s resistance to introgression has been documented (*42, 55, 56*), and thus the higher divergence of autosomal genes in radiating genera is a likely consequence of introgression from distant species. The elevated introgression of autosomal genes in radiating genera also leads to a larger deviation in the divergence of individual genes than that in non-radiating genera (orange vs. pink in Fig. 4C). More directly, we detected introgressed regions from relatively distant species by ABBA-BABA tests in 2,557 radiating triplets of species and 115 non-radiating ones (Table S12). We find that radiation clusters possess significantly more introgressed genes than non-radiating lineages at the same divergence level (Fig. 4D).

Next, we investigated the functions of introgressed genes. In the radiation clusters, we identified 2,273 genes that tend (P-value < 0.05) to introgress and 2,362 genes that are more resistant to introgression (P-value < 0.05). In the non-radiating lineages, we identified 2,159 genes that are more likely (P-value < 0.05) to introgress and 4,001 genes that never introgressed in any of the 115 triplets. Functional analysis of genes that tend to introgress between distant relatives in radiating genera versus those that are resistant to introgression using GO term enrichment is shown in Table S13. Unexpectedly, species from radiating clusters tend to acquire genes encoding proteins that function in mate recognition and selection (GO terms: courtship behavior, odorant binding, neuropeptide hormone activity), and with roles in transcription and translation. Such genes resist introgression in non-radiating genera (Fig. 4E). Divergence in mate choice genes along with transcription/translation regulators is typically associated with speciation and may confer hybrid incompatibilities (*54, 57*). Introgression of such genes from distant species may facilitate speciation by promoting reproductive isolation.

Furthermore, radiating clusters introgress genes related to morphogenesis (GO terms: embryo development, pattern specification process, eye-antennal disc development, sex determination), while non-radiating genera do not. Acquiring new morphological traits by individual mutations may be slow, but introgressed alleles would immediately prompt a variety of phenotypes, e.g., introgression of wing pattern genes explains mimicry among *Heliconius* species (*3, 58*). Finally, the trend of radiating clusters to exchange genes with roles in salivary gland development and peritrophic matrix (*59*) (a membrane structure present between food and midgut tissue) formation may allow caterpillars’ adaptation to additional food plants, thus expanding ecological niches of these species. Similarly, preferred introgression of energy-producing mitochondrial genes, DNA repair factors, and starvation-resistance molecules is observed in radiating clusters. Such exchanges may increase the chance for a species to survive hard conditions by gathering advantageous alleles that originated in other species.

## Uneven evolutionary rates and positive selection

The phylogenetic tree constructed from all protein-coding genes reveals drastic variation in evolutionary rates between lineages of USC butterflies. Distances from the root to all leaves in this tree show a wide distribution, and some lineages evolve at least twice as fast as others (Fig. 5A). The clades with the largest rate (> 0.55 in Fig. 5A) prominently standing out from the rest are the blues (Polyommatinae) and the whites (Pierini). They may have experienced rapid evolution due to specialized interactions with ants (the blues) (*60, 61*) and adaptation to caterpillar feeding on mustards (the whites), which are toxic to many insects (*62*).

**Fig. 5.**
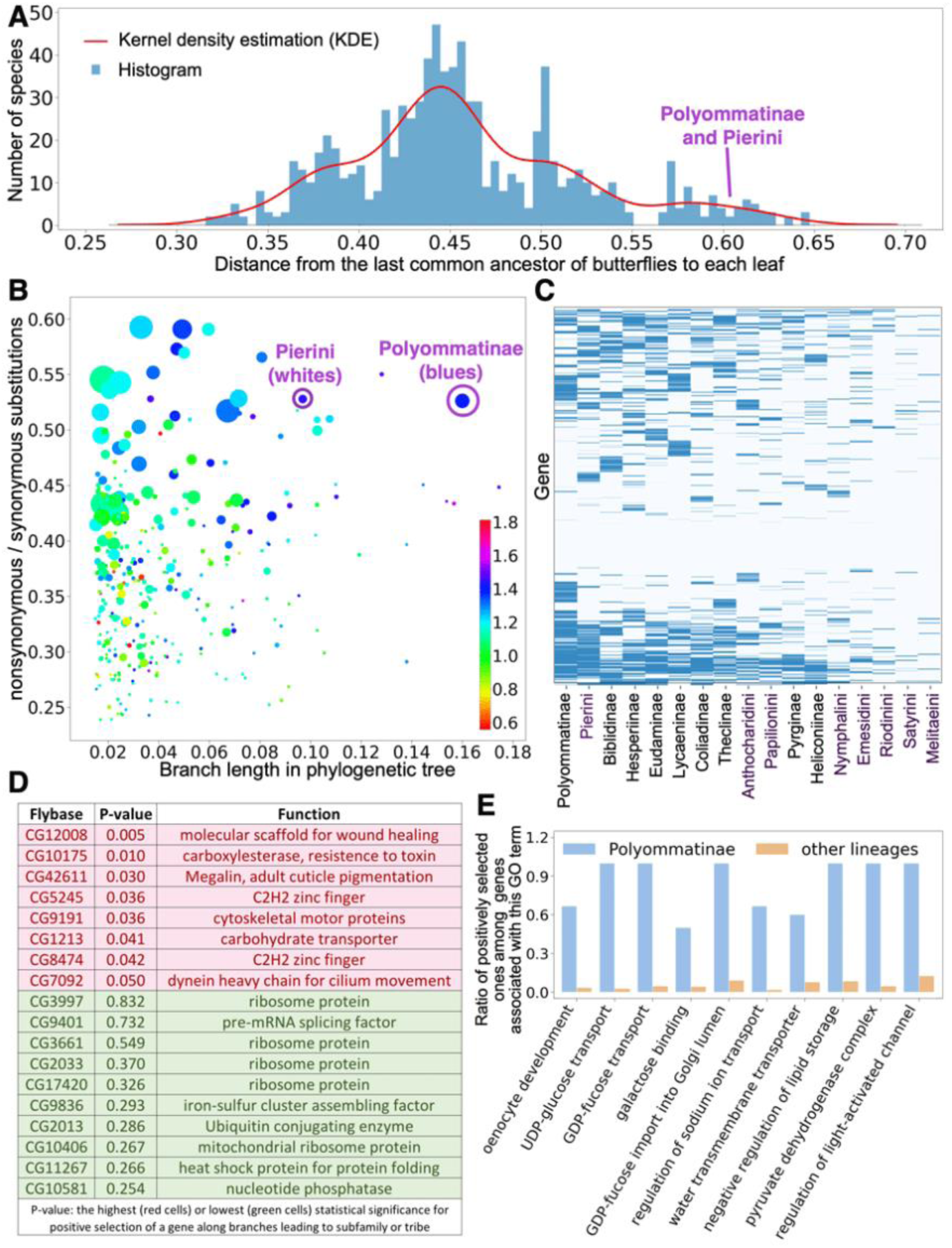
Patterns of positive selection in butterflies and adaptations unique to the blues (Polyommatinae). (A) Histogram and kernel density estimation of the distance from the last common ancestor of butterflies to each species in the USC butterfly tree based on nuclear genes. (B) Branch length and the Non-synonymous Substitution Rate (NSR) along each branch. The area of each dot is linearly correlated with the number of species originating from this branch and the color encodes the NSR along this branch divided by the average NSR in the branches originating from this branch. (C) Genes (rows) that are positively selected in major butterfly lineages (columns) colored (if P-value < 0.01) by statistical significance: darker blue indicates lower P-values (details in Methods). (D) Genes that are positively selected in all lineages (red) and those showing no positive selection in any lineage (green). (E) Gene Ontology terms that are enriched for the genes positively selected in the blues (Polyommatinae) but not in other lineages.

To better understand the reasons for the variation in evolutionary rate, we studied the effects of positive selection in each branch of the phylogenetic tree using reconstructed sequences of the internal nodes. The ratio of nonsynonymous and synonymous substitutions (Y-axis) is plotted versus the length of each tree branch (X-axis, Fig. 5B). The blues (Polyommatinae) are indeed a prominent outlier, indicating that they have been evolving under stronger positive selection. Notably, branches leading to more species (size of the circles in Fig. 5B is proportional to the number of species) generally show a higher rate of nonsynonymous substitutions. Apparently, stronger positive selection may lead to the development of adaptive traits giving advantage to a lineage and enabling it to diversify more than others. Furthermore, we find that positive selection is typically lowered in children of a long branch with strong positive selection (color of circles in Fig. 5B).

We identified genes under positive selection in the longest branches leading to diverse clades (9 subfamilies and 8 tribes) using modified McDonald-Kreitman tests (*63*) (Table S14). The blues (Polyommatinae) and the whites (Pierini) have the largest number of such genes. Biclustering partitions the genes into three groups: under strong positive selection in a lineage-specific fashion (top in Fig. 5C), not positively selected in any lineage (middle in Fig. 5C), and positively selected in multiple lineages (bottom in Fig. 5C). The genes that are positively selected in all lineages (Fig. 5D, red background) include a wound healing factor, a detoxification molecule and a carbohydrate transporter, which may participate in absorption of nutrients. These genes may have helped these lineages survive in tough conditions, e.g., during the Cretaceous-Paleogene extinction. In contrast, genes with the lowest positive selection (Fig. 5D, green background) function in fundamental processes, e.g., encode ribosomal proteins. Common to all life forms and polished thoroughly by evolution, these genes have few nonsynonymous substitutions.

The blues (Polyommatinae) exhibit the fastest evolution driven by strong positive selection in the largest number of genes. Their well-documented relationship with ants may be the driver. Ants protect caterpillars of blues from predators and feed on the liquid secreted by special glands of the caterpillars (*64*). Some blue species even evolved to feed on ant larvae, while fooling the ants by producing chemicals and sounds to accept them as their own kind (*65*). We identified GO terms associated with genes that have been positively selected only in the blues (Table S15, Fig. 5E). Many of these genes encode proteins possibly related to interactions with ants, e.g., proteins of oenocyte development. In ants, oenocytes secret cuticular hydrocarbons used to recognize their nestmates (*66*), and the blues may produce similar chemicals by oenocytes to trick the ants. The ability to secrete the ant-feeding liquid from special glands in a caterpillar should require a number of transporters, and we indeed observe strong positive selection in a number of transporters for sugars, ions, and water. Finally, we find unique positive selection in metabolic proteins such as regulators for lipid storage and enzymes for carbohydrate metabolism. Caterpillars of blues frequently feed on a nutrient-rich diet, such as flowers, fruits or even ants (*61*) instead of leaves, and therefore they may have altered their metabolism to adapt to this difference in food resource and achieve fast development.

## Evolutionary hypotheses: a broader perspective

A model for gene exchange between Eukaryotic species though introgression, butterflies, like the Darwin finches (*67*), hold the promise for discovering new general principles of evolution. Errors in replication generate variations for evolution to select from. Similarly, errors in mate selection introduce genomic segments from another species, providing a shortcut to accumulating mutations. A more efficient way to generate variation than point mutations, interspecies hybridization and introgression is emerging as a powerful evolutionary force to shape the adaptive landscape in multicellular organisms (*68*).

Our investigation into the diversification pattern of all USC butterflies provides direct evidence for the role of introgression in promoting radiation. We find that radiating genera show significantly higher introgression between distantly related (non-sister) species than non-radiating genera. Surprisingly, genes that are possibly related to mate recognition and speciation, such as those involved in courtship behavior and morphogenesis are preferably introgressed between distant species in radiating genera, while such genes resist introgression in non-radiating genera. Acquisition of speciation genes from a distant relative helps an incipient species to diverge from its sister by recombination rather than by point mutations, speeding up speciation and contributing to radiation.

We observe abundant bursts of radiations in the USC butterfly tree during the last 8 Myr. However, the scarcity of such radiations before 15 Mya suggests that only a small number of species from each radiating cluster persist in time. Each radiating cluster exists as a community of closely related species that exchange genes and compete with each other for resources, but eventually survive mostly as a single lineage. We find that the genes involved in food digestion, energy production, resistance to starvation, and DNA damage tend to introgress between species in radiating genera. Adapted alleles of such genes likely play a crucial role in the survival of a species in hard conditions, such as food shortage or temperature fluctuations. Therefore, the surviving lineage may gather advantageous alleles from other lineages that had become extinct in the past, allowing it to rapidly adapt to the changing environment and avoid extinction.

In sum, the patterns of diversification, radiation, introgression, and extinction observed in USC butterflies suggest the following evolutionary model. A species spreads over a large geographic area, increasing its population size and accumulating variations by mutations. Geographical isolation between populations drives them to speciate by accumulating Dobzhansky-Muller incompatibilities (*69*). Now, evolving as a set of closely related but reproductively semi-isolated species, these organisms further diverge and adapt to their local conditions. Still porous reproductive barriers between these incipient species allow them to exchange genes, and such exchanges speed up diversification by recombining speciation genes to generate new alleles, leading to radiation. While these exchanges are uncommon and not likely to propagate throughout the population, particularly beneficial alleles may get fixed due to selective sweeps (*70*). As a result of such introgression, each species can acquire beneficial traits from others and become adapted to more diverse conditions. Then, either as a result of direct competition or toughening environmental conditions, most species undergo extinction, and the species that gathered the most beneficial alleles moves forward in time. The cycle of diversification-radiation-introgression-extinction repeats, generating the diversification patterns we observe today.

Looking beyond butterflies, we see parallels in the evolution of Hominids. Diversified into several species including Neanderthals and Denisovans in the last 1 Myr, *Homo* experienced introgression as we see in butterflies. Most Non-African modern human populations contain about 2% DNA from Neanderthal (*71*), and the fraction of Denisovan genes varies (*72*) reaching 4-6% in Melanesians (*73*). Meanwhile, archaic human genomes also contain genes that are traced back to *Homo sapiens* (*74*). Nowadays, only modern humans survived, but genes of archaic humans stay in our genomes. Although many introgressed genes may be selected against and are being eliminated from modern human genomes with time (*75*), a fraction of them may be beneficial, increasing their frequency as a result of selection (*72*). These introgressed genes were proposed to help modern humans adapt to diverse climates (*76, 77*) and fight against pathogens (*78, 79*).

## MATERIALS AND METHODS

### Reference genome assembly and annotation

We sequenced, assembled and annotated genomes as previously described (*6, 9, 11*). Briefly, paired-end libraries with insert sizes 250 bp and 500 bp and mate-pair libraries with insert sizes 2 kb, 5 kb, and 10 kb were constructed and sequenced. All reads were processed by Trimmomatic (*80*) to remove adapter sequences and low-quality (quality score < 20) bases, and by Quake (*81*) to correct sequencing errors (*81*). We used Platanus (*82*) to assemble the genomes. The initial assemblies from Platanus were frequently redundant. The highly heterozygous equivalent segments in the paternal and maternal chromosomes were treated separately, and thus they were present twice in the assemblies. We detected and corrected such problems as described before (*6, 9, 11*).

The repeats in the genomes were identified by RepeatModeler (*83*). In addition, since repeats with highly similar sequences may be erroneously combined into one in the genome assemblies, we identified them using very high sequence depth (more than 4 times of the expected value) after all the sequence reads were mapped to the draft genomes using BWA (*84*). We combined the repeats identified by RepeatModeler and our sequence depth criteria with repeats in Repbase (*85*) to generate species-specific repeat libraries, and these libraries were supplied to RepeatMasker (*86*) to annotate repeats in the genomes.

We annotated protein-coding genes using three approaches: homology-based, transcript-based, and *de novo* gene prediction. We used protein sets from *Papilio machaon (21), Pieris rapae (9), Calycopis cecrops (10), Calephelis nemesis (8), Danaus plexippus (4), Cecropterus lyciades (7), Bombyx mori (87)*, and *Drosophila melanogaster (88)* as references for homology-based annotation. These references include one species from each of the 6 butterfly families and the established model organisms, silkworm and fruit fly with expected high quality of gene models. The reference protein sets were aligned to draft genomes using Exonerate (*89*). We had RNA-seq reads for 22 of the 23 new reference genomes, and we used TopHat/Cufflinks pipeline (*90, 91*) to perform transcript-based annotation for them. Three *de novo* gene prediction methods: Augustus (*92*), GeneMark_ES (*93*), and SNAP (*94*) were used to obtain *de novo* gene annotations. We trained these *de novo* predictors for each species using confident gene models derived from the consensus between transcript-based and homology-based annotations. Finally, annotations by different approaches were combined in EvidenceModeler (*95*) to obtain their consensus as the final gene predictions. We predicted the functions of these proteins by finding the closest sequence hits in Flybase (*96*) and Swissprot (*97*) using BLASTP (E-value < 0.00001) and transferred the Gene Ontology (GO) (*98*) terms and function annotations.

### Identification and analysis of Ultra Conserved Elements (UCE) in the butterfly genomes

A total of 36 reference genomes of USC butterflies were used in our study, and 31 of them were sequenced by us. The five genomes sequenced by others *(4, 21-23)* were obtained from LepBase v4 (http://ensembl.lepbase.org/index.html). We used the published gene models but annotated the protein function using our pipeline described above. For each butterfly family, we selected a representative genome with high N50: *Heliconius erato* (Nymphalidae) (*23*), *Megathymus ursus* (Hesperiidae) (*6*), *Apodemia nais* (Riodinidae), *Heraclides cresphontes* (Papilionidae), *Pieris rapae* (Pieridae), and *Feniseca tarquinius* (Lycaenidae). We masked the repetitive regions in these genomes using RepeatMasker (*86*) and removed short (less than 10 kb) scaffolds. We used *Heliconius erato* (assembly with the highest N50) as the primary reference, and aligned other five genomes to it by LASTZ (*27*). Aligned segments were processed sequentially by axtChain (*99*) and ChainNet (*100*) to generate pairwise whole genome alignments. These pairwise alignments were merged into a multiple genome alignment using MULTIZ (*101*). For segments where all six genomes were aligned, we counted identical positions in overlapping sliding windows of 50 bp. Windows with more than 96% identical positions in all six genomes were considered candidate UCEs, and adjacent candidate UCEs were merged.

We identified 764 candidate UCEs from the 6 selected genomes and searched for these UCEs in the remaining 30 genomes using BLASTN (*102*). A UCE was considered valid in a genome if a single confident hit (E-value < 0.001) can be found with higher than 96% sequence identity to the query UCE. As a result, we obtained 530 UCEs confirmed in at least 30 of the 36 genomes. We found genes that are less than 10 kb away from the UCEs in *Heliconius erato* genome, and detected GO terms that are preferably associated with these genes using binomial tests (p = probability for this GO term to be associated with any gene in the genome, m = number of genes near UCE that are associated with this GO term, N: total number of genes that are less than 10 kb away from UCEs). The most significant GO terms (false discovery rate (*103*) < 0.1) were visualized in REVIGO (*29*).

### Identification of lineage-specific gene expansions

We used OrthoMCL (*104*) to identify the groups of orthologous proteins encoded by the 36 reference genomes. We mapped proteins in each group to the closest protein (BLAST E-value < 0.00001) of *Drosophila melanogaster* from Flybase (*96*). We assigned a *Drosophila* protein to an orthologous group to if more than 50% members in this group mapped to the protein. Furthermore, we merged orthologous groups if at least 50% of *Drosophila* proteins in them were the same. After merging, 5089 orthologous groups were present in at least 50% of 36 butterfly species and included *Drosophila* proteins: these groups were used in the following analysis. We used the accumulative protein length instead of protein number to identify gene expansions because number of proteins can be more easily affected by scaffold discontinuity in draft genomes and errors in annotation. In addition, gene expansions tend to occur as tandem repeats, and we used this property to identify candidate gene expansions.

Among the 36 reference genomes, we have 5, 4, 6, 2, 11, and 8 species from Papilionidae, Pieridae, Lycaenidae, Riodinidae, Nymphalidae, and Hesperiidae families, respectively. We further divided three families with more than 5 members (Lycaenidae, Nymphalidae, and Hesperiidae) into smaller groups. Lycaenidae were partitioned into Polyommatinae (3 genomes) and the rest (3 genomes); Hesperiidae were partitioned into Hesperiinae (3 genomes) and the rest (5 genomes). We grouped Nymphalinae and Heliconiinae subfamilies from Nymphalidae because they both have scoli covering the caterpillars, and other Nymphalidae we sequenced (no scoli) were considered the other group. Thus, we partitioned the reference genomes into 9 lineages. We calculated the total length of proteins in each orthologous group for each lineage, and identified lineage-specific gene expansion using three criteria: (1) the average accumulative protein length for species within this lineage is at least twice the average for other species; (2) the minimal total protein length for species in this lineage is larger than 90% species from other lineages; (3) 50% of proteins in this lineage are encoded next to another protein from the same orthologous group in the genomes. A total of 22 cases passed all these criteria. We manually inspected them to remove 8 cases without functional annotation and those of possible transposon origins. The remaining cases are shown in Fig. 1C.

### Protein-coding sequence assembly for all USC butterflies

We developed a pipeline to assemble the protein-coding sequences from the whole genome shotgun reads of a target species using the protein sequences of a reference genome as baits, and the genome of a species more distant from the reference species than the target species as an outgroup. Different reference genomes were used for the same target species for different purposes. To obtain the most complete protein sets for each species, we used the closest reference genome; for phylogenetic analysis of a family, we selected a single reference for all species in that family; for phylogenetic analysis of all USC butterflies together, we used *Cecropterus lyciades* (*7*) as reference.

We split the reference proteins into exons and searched against sequence reads of a target species using DIAMOND (*105*) with the following parameters: -l 1 --comp-based-stats 1 --masking 0 - evalue 0.01. From the DIAMOND results of all exons in the reference, we kept the reads that could be unambiguously mapped to one locus by both E-value (< 1e-5 × E-value for other loci) and sequence identity (> identity for other loci + 10). We further filtered the alignments by requiring at least 80% coverage over the reads or the query exon and sequence identity higher than that between the reference and the outgroup. Because we used a number of old dry museum specimens whose DNA can be contaminated by fungi, bacteria and surrounding specimens, we applied the following protocol to detect and remove contaminants.

For each 30 bp sliding window applied to the alignment between the reference and the reads, we clustered all the reads into groups of similar sequences using the following procedure. We ranked reads by their sequence identity to the query from high to low. The first read initiated a cluster. Starting from the second read, a new read was compared to the first sequence of each cluster and assigned to the first cluster whose first sequence had no more than one mismatch from the current sequence. If a new read could not be assigned to existing clusters, a new cluster was initiated with this read as the first member. For each cluster, we computed its size and the average number of mismatches to the query, and we considered a cluster to be good if its size was at least half of the largest cluster size and number of mismatches was no larger than minimal mismatches + 2. If the number of good clusters was no more than 2 (diploid genome), we marked the reads that were not included in the good clusters as bad reads; otherwise, we marked all reads as bad. All the bad reads were discarded.

The dominant nucleotide (frequency > 0.6) at each position in the sequence alignment after this cleaning procedure was used to generate the exon sequences of the target species. The exon sequences were further translated to amino acid sequences and sequences of different exons of a protein were concatenated to obtain the protein sequence of the target species.

### Phylogenetic analysis of USC butterflies

Phylogenetic trees were constructed for each butterfly family from the protein-coding sequences using one reference genome per family: *Pterourus glaucus* (Papilionidae), *Phoebis sennae* (Pieridae), *Calycopis cecrops* (Lycaenidae), *Calephelis nemesis* (Riodinidae), *Heliconius melpomene* (Nymphalidae), and *Cecropterus lyciades* (Hesperiidae). Since the sequences for other samples were assembled using the reference as baits, they were all aligned to the reference and could be readily converted to multiple sequence alignments. Three datasets: autosomal, Z-linked and mitochondrial proteins, were used to construct phylogenetic trees. Strong conservation of gene content was reported for Lepidoptera Z chromosome (*106*). Therefore, we considered exons to be Z-linked if their best TBLASTN (*102*) hits were on *Heliconius erato* Z chromosome, and a gene to be Z-linked if more than 80% of its exons were Z-linked. Multiple sequence alignments of proteins were concatenated in each dataset, and positions containing more than 60% gaps were removed.

For autosome- and Z-chromosome-based phylogeny, we built trees for 100 partitions, and each partition was generated by randomly drawing 20 kb positions from the alignment. We used IQ-TREE (*107*) (model: GTR+I+G) to construct the maximum-likelihood trees for each partition and summarized them to obtain a consensus tree using sumtrees.py (-f 0.0) (*108*). For mitochondrial proteins, we used the entire alignment and applied IQ-TREE with model selection and 1000 fast bootstrap (-bb 1000) to construct the tree.

To resolve the relationship between families, we used a single reference genome, *Cecropterus lyciades*. Sequences for all other species were derived by mapping to this single reference, resulting in multiple sequence alignments of all USC butterflies. We constructed trees for autosomal, Z-linked and mitochondrial proteins as described above. These trees were expected to be less accurate in resolving shallower phylogeny due to the lower sequence similarity between *Cecropterus lyciades* and species in other families. Therefore, we replaced the clades for each family in these trees with the trees constructed for each individual butterfly family using python ETE3 module (*109*) to generate the USC butterfly trees used in this study. These trees were rescaled as previously described (*39, 110*) and the time axis was added to the tree constructed from all nuclear genes based on our published calibration (*8*) to match the ages of common nodes between the current and previous trees.

### Simulation of diversification process under constant speciation and extinction rate

We developed an in-house script to simulate species growth under constant speciation and extinction rates. Here, speciation rate (*R*_*S*_) is the probability for a taxon to split into two in 1 Myr, and extinction rate (*R*_*E*_) is the probability for a taxon to extinct after 1 Myr. Our simulation started with one taxon and iteratively introduced speciation and extinction events for 122 times, corresponding to the 122 Myr of butterfly evolution we observed from the data. To introduce random fluctuation, we used a random number to determine whether a taxon should undergo speciation or extinction with the expected probabilities defined by *R*_*S*_ and *R*_*E*_, respectively.

We computed the observed diversification rate per 5 Myr and the highest value is 2, which correspond to a speciation rate of 0.15 ((1 + 0.15)^5^ = 2.0) in the absence of extinctions. Therefore, we fixed *R*_*S*_ at 0.15 and tested different *R*_*E*_ in the range between 0.0 and 0.1 with an increment of 0.01. We ran 1000 simulations for each *R*_*E*_ value, and a value of 0.08 gave the best chance of producing about the same (+-10%) number of species as observed. We therefore ran 10000 simulations under these parameters (*R*_*S*_ = 0.15 and *R*_*E*_ = 0.08) and selected the trajectories that produced about the same (+-10%) number of species as observed. From these trajectories, we analyzed the apparent diversification rate (a result of both speciation and extinction) every 5 Myr and counted the number of radiation events at each time point for comparison with the observed data.

### Identification of radiation events and radiation-associated proteins

We identified radiating nodes in the tree as those generating at least 5 lineages in less than 2 Myr, and if one clade started with a radiating node and included at least 6 species, we consider it a radiation cluster. We thus identified 18 non-overlapping radiation clusters from 18 genera: *Pterourus, Colias, Callophrys, Satyrium, Euphilotes, Celastrina, Oeneis, Polygonia, Phyciodes, Chlosyne, Boloria, Speyeria, Cecropterus, Erynnis, Euphyes, Hesperia, Atrytonopsis*, and *Agathymus*. For each radiating genus, we used other closely related genera as external references to identify proteins with elevated divergence within this radiation cluster.

For each protein, we calculated its average divergence for any pair of species within the radiation cluster (*DIV*_*internal*_), and the average divergence between any species in the cluster and external references from other genera (*DIV*_*external*_). We mapped proteins in each genus to their closest (E-value < 0.00001) *Drosophila* proteins in Flybase. If multiple proteins were mapped to the same *Drosophila* protein, we computed the average *DIV*_*internal*_ and *DIV*_*external*_ weighted by the length of each protein. We detected proteins with elevated divergence in radiation clusters using two criteria: first, *DIV* _*internal*_ is significantly (P-value < 0.01) higher than the average *DIV* _*internal*_ over all proteins; second, *DIV*_*internal*_ is higher than *DIV* _*external*_ by at least 1.5 times. The second criterion ensured that we selected proteins that tended to diverge within radiation clusters instead of generally fast evolving ones, because *DIV*_*external*_ should be higher than *DIV*_*internal*_ for most proteins due to the larger evolutionary distance between genera than within a genus. These criteria selected 273–846 proteins in each radiating genus, and we considered them to be radiation-associated proteins.

We only considered proteins that were present in at least 9 radiating genera and we identified recurrent radiation-associated proteins using binomial tests (p = total number of radiation-associated proteins in all 18 genera / total number of proteins being analyzed in all 18 radiating genera, m = number of genera where this protein is among the radiation-associated proteins, N = number of genera where this protein is being analyzed, alternative hypothesis: greater). Proteins with P-values less than 0.05 were considered as recurrent radiation-associated proteins, and we further identified GO terms that were enriched among them using another binomial test (p = probability for this GO term to be associated with any protein, m = total number of recurrent radiation-associated proteins that are associated with this GO term, N = total number of recurrent radiation-associated proteins).

### Comparison of radiating and non-radiating lineages

To investigate the differences between radiating and non-radiating genera, we first compared sister species in both types of genera. We used the 18 radiating clusters identified above, and we found non-radiating genera using the following criteria: (1) the genus does not contain any consecutive speciation events separated by less than 0.67 Myr; (2) the genus is not rich in species south of the United States. We extracted 63 pairs of sister species from radiating genera. The distances between these pairs in the phylogenetic tree were mostly below 0.03 substitutions per position. We further selected 68 pairs of sister species whose distance in the tree were below 0.03 from non-radiating lineages. We binned the sister species pairs from radiating or non-radiating genera according to their average divergence in gene sequences to the following bins: 0.05 - 0.1, 0.1 - 0.15, 0.15 - 0.2, 0.2 - 0.25. We partitioned genes into autosomal and Z-linked ones. In each bin and each partition, we observed the distribution of sequence divergence (percentage of positions with different nucleotides) for individual genes in radiating and non-radiating genera by Python seaborn package (https://seaborn.pydata.org/).

The comparisons of sister species suggested a higher level of introgression in radiating genera, and thus we further tested the extent of introgression more rigorously using ABBA-BABA tests (*71*). ABBA-BABA test requires 4 taxa following a tree topology ((S1,S2),S3),O; where S1 and S2 are closely related, S3 is more distant and O is the outgroup. The test is used to identify introgression from the distant group S3 to either S1 or S2 based on excessive similarity between S3 and S1 or S2, respectively. We identified taxa following the topology of ((S1,S2),S3) among radiating genera and non-radiating genera, and we required the grouping of S1 with S2 to be strongly supported with a bootstrap value of 1. We obtained outgroups from the sister genera. The divergence between taxon S3 and taxon S1 or S2 in the cases we identified from radiating genera was mostly below 0.06, and therefore we selected the cases from non-radiating genera with the same level of divergence (< 0.06). As a result, we obtained 115 non-radiating cases and 2,557 radiating cases.

For each gene in each case, we counted the number of positions following the pattern of ABBA or BABA in taxa S1, S2, S3, and O. A pattern of ABBA means that taxa S1 and O share the same nucleotide, and taxa S2 and S3 share the same nucleotide that is different from S1. A pattern of BABA means that taxa S2 and O share the same nucleotide, and taxa S1 and S3 share the same nucleotide that is different from S2. Since we used multiple taxa as outgroups, and these outgroups may not support the same pattern, we counted the fraction of outgroups supporting a certain pattern at each position. In the absence of introgression (null hypothesis), the expected total number of ABBA positions should be equal to the total number of BABA positions. We tested significant deviation from the null hypothesis using binomial tests (p = 0.5, m = count of ABBA position, N = count of ABBA or BABA position). If a gene has significantly (P-value < 0.05) more ABBA position, we consider it to be introgression between S3 and S2, while significantly more BABA positions indicate introgression between S3 and S1.

We identified 2,273 genes that were significantly more likely (P-value < 0.05, alternative hypothesis: greater) to introgress and 2,362 genes that were resistant (P-value < 0.05, alternative hypothesis: less) to introgression among radiating genera using binomial tests (p = average fraction of introgressed genes in all cases, m = number of cases where this gene is introgressed, N = number of cases where this gene is being analyzed). Meanwhile, for the non-radiating genera, we identified 2,159 genes that were more likely (P-value < 0.05) to introgress and 4,001 genes that never introgressed in the 115 cases. We analyzed the functional enrichment of genes prone to introgression versus resistant ones in both radiating and non-radiating genera using GO terms as described above.

### Reconstruction of ancestral sequences and analysis of selection pressure

To study the evolutionary history and adaptation in different lineages of butterflies, we reconstructed the sequence for each gene at each node of the phylogenetic tree of USC butterflies. We derived the sequence of a target node based on its sister node and its two children using a fast in-house script. The probability of each nucleotide *i* at a position was computed using the following formula:

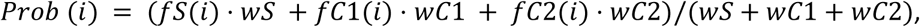

where *fS*(*i*), *fC*1(*i*), and *fC*2(*i*) were the frequencies of nucleotide *i* in the sister node, and the two children nodes, respectively; *wS, wC*1, and *wC*2 were the weights of the sister and two children nodes, and the weights were inversely correlated with the distances (in the USC butterfly tree of nuclear genes) between these nodes and the target node.

For each tree branch starting from node A and ending with node B, we compared each gene’s DNA sequences at nodes A and B. For each codon in a gene, we enumerated all possible single-substitution paths to change form the codon of node A to that of node B, and we considered the path with the lowest number of nonsynonymous substitutions (most parsimonious) to be the most likely path. We recorded these most likely substitution paths for each branch and each gene and counted the number of synonymous (*N*_*SS*_) and nonsynonymous substitutions (*N*_*NS*_) along the paths. The total *TN*_*SS*_ and *TN*_*NS*_ were obtained as the sum of *N*_*SS*_ and *N*_*NS*_ over all genes along a tree branch, and the ratio between them (*TN*_*NS*_/*TN*_*SS*_) was used as an indicator of the positive selection in the branch.

### Identification and analysis of positively selected proteins

To investigate both the universal and lineage-specific adaptation, we identified all the relatively long branches (branch length > 0.03 substitutions per position) leading to an entire subfamily, tribe or subtribe with at least 10 species in the United States and Canada. We manually inspected these branches to select a set of representatives following three rules: (1) these representatives should not overlap in species belonging to them; (2) a group originating after a longer branch and with more species is preferred. A total of 17 lineages were selected, including 9 subfamilies, 7 tribes and 1 subtribe. We identified genes showing significant positive selection along the branches leading to these lineages, respectively, using modified McDonald-Kreitman (MK) (*63*) tests.

A standard MK test compares the rate of nonsynonymous substitutions in a gene between species to the nonsynonymous polymorphisms rate within species (*63*). We generalized this method and instead evaluated if the nonsynonymous substitution rate of a gene along a tree branch was significantly higher than average nonsynonymous substitution rate in lineages originating from this branch. The rationale is that if some derived beneficial trait, such as mutualism between the blues and ants, originated in a branch, its offspring may tend to retain it by reducing the nonsynonymous substitution rate. Significance of higher nonsynonymous substitution rate in a gene along a branch was evaluated using a binomial test (p = rate of nonsynonymous substitution among species in a lineage, m = number of nonsynonymous substitutions along the branch leading to this lineage, N = number of all substitutions along the branch). To compare the positively selected genes in different lineages, we mapped genes in the 17 lineages to *Drosophila* genes, and rescaled the P-values for positive selection to significance scores as: *score* = 1 − *P* · 100, *if P* < 0.01; *score* = 0, *if P* > 0.01. We clustered the resulting score for each gene in each lineage using the clustermap function in Python seaborn package (https://seaborn.pydata.org/). In addition, GO term enrichment analysis of positively selected genes in each lineage was used to capture the functions of these genes.

## ACKNOWLEDGMENTS

We acknowledge Leina Song and Ping Chen for excellent technical assistance. We are grateful to David Grimaldi and Courtney Richenbacher (AMNH: American Museum of Natural History, New York, NY, USA), Jason Weintraub (ANSP: Academy of Natural Sciences of Drexel University, Philadelphia, PA, USA), Jonathan P. Pelham (BMUW: Burke Museum of Natural History and Culture, Seattle, WA, USA), Vince Lee and the late Norm Penny (CAS: California Academy of Sciences, San Francisco, CA, USA), Boris Kondratieff (CSUC: Colorado State University Collection, Fort Collins, CO, USA), Crystal Maier and Rebekah Baquiran (FMNH: Field Museum of Natural History, Chicago, FL, USA), Weiping Xie (LACM: Los Angeles County Museum of Natural History, Los Angeles, CA, USA), Andrew D. Warren and Debbie Matthews-Lott (MGCL: McGuire Center for Lepidoptera and Biodiversity, Gainesville, FL, USA), Edward G. Riley, Karen Wright, and John Oswald (TAMU: Texas A&M University Insect Collection, College Station, TX, USA), Alex Wild (TMMC: University of Texas Biodiversity Center, Austin, TX, USA), Jeff Smith and Lynn Kimsey (UCDC: Bohart Museum of Entomology, University of California, Davis, CA, USA), Robert K. Robbins, John M. Burns, and Brian Harris (USNM: National Museum of Natural History, Smithsonian Institution, Washington, DC, USA) for granting access to the collections under their care and for stimulating discussions; to Jim P. Brock, Jack S Carter, Bill R. Dempwolf, James McDermott, the late Edward C. Knudson (specimens now at MGCL), Harry Pavulaan, James A. Scott, John A. Shuey, and Mark Walker for specimens and leg samples. Greg Kareofelas and Matthew Garhart collected needed specimens and placed them in RNAlater for molecular analysis. Boris Kondratieff, Chuck Harp, and James Scott curated the butterfly collection at the C. P. Gillette Museum of Arthropod Diversity, Colorado State University, which facilitated the accurate sampling of remaining species needed to complete the analysis. Evi Buckner-Opler assisted by providing emotional and logistic support and helped to collect specimens. We are indebted to Texas Parks and Wildlife Department (Natural Resources Program Director David H. Riskind) for the research permit 08-02Rev, to U. S. National Park Service for the research permits: Big Bend (Raymond Skiles) for BIBE-2004-SCI-0011 and Yellowstone (Erik Oberg and Annie Carlson) for YELL-2017-SCI-7076 and to the National Environment & Planning Agency of Jamaica for the permission to collect specimens. We acknowledge the Texas Advanced Computing Center (TACC) at The University of Texas at Austin for providing HPC resources. The study has been supported in part by grants from the National Institutes of Health GM127390 and the Welch Foundation I-1505.

## REFERENCES

1. V. Nazari, L. Evans, Butterflies of Ancient Egypt. Journal of the Lepidopterists’ Society 69, 242–267 (2015).

2. J. A. Scott, The Butterflies of North America: A Natural History and Field Guide. (Standford University Press, Stanford, CA, 1986), pp. xiii + 583 pp.

3. Heliconius Genome Consortium, Butterfly genome reveals promiscuous exchange of mimicry adaptations among species. Nature 487, 94–98 (2012).

4. S. Zhan, C. Merlin, J. L. Boore, S. M. Reppert, The monarch butterfly genome yields insights into long-distance migration. Cell 147, 1171–1185 (2011).

5. S. Nallu et al., The molecular genetic basis of herbivory between butterflies and their host plants. Nat Ecol Evol 2, 1418–1427 (2018).

6. Q. Cong, W. Li, D. Borek, Z. Otwinowski, N. V. Grishin, The Bear Giant-Skipper genome suggests genetic adaptations to living inside yucca roots. Mol Genet Genomics 294, 211–226 (2019).

7. J. Shen, Q. Cong, D. Borek, Z. Otwinowski, N. V. Grishin, Complete Genome of Achalarus lyciades, The First Representative of the Eudaminae Subfamily of Skippers. Curr Genomics 18, 366–374 (2017).

8. Q. Cong et al., The first complete genomes of Metalmarks and the classification of butterfly families. Genomics 109, 485–493 (2017).

9. J. Shen et al., Complete genome of Pieris rapae, a resilient alien, a cabbage pest, and a source of anti-cancer proteins. F1000Res 5, 2631 (2016).

10. Q. Cong et al., Complete genomes of Hairstreak butterflies, their speciation, and nucleo-mitochondrial incongruence. Sci Rep 6, 24863 (2016).

11. Q. Cong, D. Borek, Z. Otwinowski, N. V. Grishin, Skipper genome sheds light on unique phenotypic traits and phylogeny. BMC genomics 16, 639 (2015).

12. A. Y. Kawahara, J. W. Breinholt, Phylogenomics provides strong evidence for relationships of butterflies and moths. Proceedings of the Royal Society B: Biological Sciences 281, 20140970 (2014).

13. M. Heikkila, L. Kaila, M. Mutanen, C. Pena, N. Wahlberg, Cretaceous origin and repeated tertiary diversification of the redefined butterflies. Proceedings. Biological sciences 279, 1093–1099 (2012).

14. J. P. Pelham, Catalogue of the Butterflies of the United States and Canada. Journal of Research on the Lepidoptera 40, 1–658 (2008).

15. J. P. Pelham. (http://www.butterfliesofamerica.com/US-Can-Cat.htm, 2019).

16. U. S. Fish and Wildlife Service. (https://ecos.fws.gov/ecp0/pub/SpeciesReport.do?groups=I&listingType=L, 2019).

17. S. Wang, S. Sun, Z. Li, R. Zhang, J. Xu, Accurate De Novo Prediction of Protein Contact Map by Ultra-Deep Learning Model. PLoS Comput Biol 13, e1005324 (2017).

18. Q. Cong, I. Anishchenko, S. Ovchinnikov, D. Baker, Protein interaction networks revealed by proteome coevolution. Science 365, 185–189 (2019).

19. A. J. Riesselman, J. B. Ingraham, D. S. Marks, Deep generative models of genetic variation capture the effects of mutations. Nat Methods 15, 816–822 (2018).

20. Q. Cong et al., When COI barcodes deceive: complete genomes reveal introgression in hairstreaks. Proceedings. Biological sciences 284, (2017).

21. X. Li et al., Outbred genome sequencing and CRISPR/Cas9 gene editing in butterflies. Nat Commun 6, 8212 (2015).

22. K. R. L. van der Burg et al., Contrasting Roles of Transcription Factors Spineless and EcR in the Highly Dynamic Chromatin Landscape of Butterfly Wing Metamorphosis. Cell Rep 27, 1027–1038 e1023 (2019).

23. N. J. Nadeau et al., Population genomics of parallel hybrid zones in the mimetic butterflies, *H. melpomene* and *erato*. Genome research 24, 1316–1333 (2014).

24. F. A. Simao, R. M. Waterhouse, P. Ioannidis, E. V. Kriventseva, E. M. Zdobnov, BUSCO: assessing genome assembly and annotation completeness with single-copy orthologs. Bioinformatics 31, 3210–3212 (2015).

25. A. Derti, F. P. Roth, G. M. Church, C. T. Wu, Mammalian ultraconserved elements are strongly depleted among segmental duplications and copy number variants. Nat Genet 38, 1216–1220 (2006).

26. D. Polychronopoulos, J. W. D. King, A. J. Nash, G. Tan, B. Lenhard, Conserved non-coding elements: developmental gene regulation meets genome organization. Nucleic Acids Res 45, 12611–12624 (2017).

27. R. S. Harris, Improved pairwise alignment of genomic DNA. Ph. D., (2007).

28. D. A. Petrov, Evolution of genome size: new approaches to an old problem. Trends Genet 17, 23–28 (2001).

29. F. Supek, M. Bosnjak, N. Skunca, T. Smuc, REVIGO summarizes and visualizes long lists of gene ontology terms. PLoS One 6, e21800 (2011).

30. M. K. Dhar, A. Koul, S. Kaul, Farnesyl pyrophosphate synthase: a key enzyme in isoprenoid biosynthetic pathway and potential molecular target for drug development. N Biotechnol 30, 114–123 (2013).

31. T. Eisner, Y. C. Meinwald, Defensive Secretion of a Caterpillar (*Papilio*). Science 150, 1733–1735 (1965).

32. N. E. Pierce, Lycaenid Butterflies and Ants: Selection for Nitrogen-Fixing and Other Protein-Rich Food Plants. The American Naturalist 125, 888–895 (1985).

33. P. Chelikani, I. Fita, P. C. Loewen, Diversity of structures and properties among catalases. Cell Mol Life Sci 61, 192–208 (2004).

34. A. D. Warren, J. R. Ogawa, A. V. Z. Brower, Revised classification of the family Hesperiidae (Lepidoptera: Hesperioidea) based on combined molecular and morphological data. Syst Entomol 34, 467–523 (2009).

35. R. K. Sahoo, A. D. Warren, S. C. Collins, U. Kodandaramaiah, Hostplant change and paleoclimatic events explain diversification shifts in skipper butterflies (Family: Hesperiidae). BMC Evolutionary Biology 17, 174 (2017).

36. M. Mutanen, N. Wahlberg, L. Kaila, Comprehensive gene and taxon coverage elucidates radiation patterns in moths and butterflies. Proceedings. Biological sciences 277, 2839–2848 (2010).

37. Q. Cong, J. Zhang, J. Shen, N. V. Grishin, Fifty new genera of Hesperiidae (Lepidoptera). Insecta Mundi 0731, 1–56 (2019).

38. J. Zhang, Q. Cong, J. Shen, P. A. Opler, N. V. Grishin, Changes to North American butterfly names. The Taxonomic Report of the International Lepidoptera Survey 8, 1–11 (2019).

39. W. Li et al., Genomes of skipper butterflies reveal extensive convergence of wing patterns. Proc Natl Acad Sci U S A 116, 6232–6237 (2019).

40. M. C. Fontaine et al., Mosquito genomics. Extensive introgression in a malaria vector species complex revealed by phylogenomics. Science 347, 1258524 (2015).

41. A. Suh, L. Smeds, H. Ellegren, The Dynamics of Incomplete Lineage Sorting across the Ancient Adaptive Radiation of Neoavian Birds. PLoS Biol 13, e1002224 (2015).

42. S. H. Martin et al., Genome-wide evidence for speciation with gene flow in Heliconius butterflies. Genome research 23, 1817–1828 (2013).

43. S. Sankararaman et al., The genomic landscape of Neanderthal ancestry in present-day humans. Nature 507, 354–357 (2014).

44. D. A. Turissini, D. R. Matute, Fine scale mapping of genomic introgressions within the *Drosophila yakuba* clade. PLoS Genet 13, e1006971 (2017).

45. M. L. M. Lalonde, B. S. McCullagh, J. M. Marcus, The taxonomy and population structure of the buckeye butterflies (Genus *Junonia*, Nymphalidae: Nymphalini) of Florida, USA. Journal of the Lepidopterists’ Society 72, 97–115 (2018).

46. M. M. L. Lalonde, J. M. Marcus, Getting western: biogeographical analysis of morphological variation, mitochondrial haplotypes and nuclear markers reveals cryptic species and hybrid zones in the *Junonia* butterflies of the American southwest and Mexico. Syst Entomol 44, 465–489 (2019).

47. J. T. Weir, D. Schluter, The latitudinal gradient in recent speciation and extinction rates of birds and mammals. Science 315, 1574–1576 (2007).

48. P. R. Renne et al., Time scales of critical events around the Cretaceous-Paleogene boundary. Science 339, 684–687 (2013).

49. Y. Terai, N. Morikawa, K. Kawakami, N. Okada, The complexity of alternative splicing of hagoromo mRNAs is increased in an explosively speciated lineage in East African cichlids. Proc Natl Acad Sci U S A 100, 12798–12803 (2003).

50. L. A. Smith, A. A. Peixoto, E. M. Kramer, A. Villella, J. C. Hall, Courtship and visual defects of cacophony mutants reveal functional complexity of a calcium-channel alpha1 subunit in *Drosophila*. Genetics 149, 1407–1426 (1998).

51. T. Sakai, N. Ishida, Circadian rhythms of female mating activity governed by clock genes in *Drosophila*. Proc Natl Acad Sci U S A 98, 9221–9225 (2001).

52. R. Allada, B. Y. Chung, Circadian organization of behavior and physiology in *Drosophila*. Annu Rev Physiol 72, 605–624 (2010).

53. K. L. Mack, M. W. Nachman, Gene Regulation and Speciation. Trends Genet 33, 68–80 (2017).

54. C. R. Landry et al., Compensatory cis-trans evolution and the dysregulation of gene expression in interspecific hybrids of Drosophila. Genetics 171, 1813–1822 (2005).

55. R. Storchova, J. Reif, M. W. Nachman, Female heterogamety and speciation: reduced introgression of the Z chromosome between two species of nightingales. Evolution 64, 456–471 (2010).

56. L. Cortes-Ortiz et al., Reduced Introgression of Sex Chromosome Markers in the Mexican Howler Monkey (*Alouatta palliata* x *A. pigra*) Hybrid Zone. Int J Primatol 40, 114–131 (2019).

57. A. Y. Tulchinsky, N. A. Johnson, W. B. Watt, A. H. Porter, Hybrid incompatibility arises in a sequence-based bioenergetic model of transcription factor binding. Genetics 198, 1155–1166 (2014).

58. A. V. Brower, Introgression of wing pattern alleles and speciation via homoploid hybridization in *Heliconius* butterflies: a review of evidence from the genome. Proceedings. Biological sciences 280, 20122302 (2013).

59. M. J. Lehane, Peritrophic matrix structure and function. Annu Rev Entomol 42, 525–550 (1997).

60. D. R. Nash, T. D. Als, R. Maile, G. R. Jones, J. J. Boomsma, A mosaic of chemical coevolution in a large blue butterfly. Science 319, 88–90 (2008).

61. N. E. Pierce et al., The ecology and evolution of ant association in the Lycaenidae (Lepidoptera). Annu Rev Entomol 47, 733–771 (2002).

62. F. Beran et al., Phyllotreta striolata flea beetles use host plant defense compounds to create their own glucosinolate-myrosinase system. Proc Natl Acad Sci U S A 111, 7349–7354 (2014).

63. J. H. McDonald, M. Kreitman, Adaptive protein evolution at the Adh locus in Drosophila. Nature 351, 652–654 (1991).

64. K. Fiedler, Ants That Associate with Lycaeninae Butterfly Larvae: Diversity, Ecology and Biogeography. Diversity and Distributions 7, 45–60 (2001).

65. F. Barbero, J. A. Thomas, S. Bonelli, E. Balletto, K. Schonrogge, Queen ants make distinctive sounds that are mimicked by a butterfly social parasite. Science 323, 782–785 (2009).

66. F. Barbero, Cuticular Lipids as a Cross-Talk among Ants, Plants and Butterflies. Int J Mol Sci 17, (2016).

67. C. R. Darwin, On the origin of species by means of natural selection, or preservation of favoured races in the struggle for life. (John Murray, London, 1859), pp. 512.

68. N. B. Edelman et al., Genomic architecture and introgression shape a butterfly radiation. Science 366, 594–599 (2019).

69. H. A. Orr, Dobzhansky, Bateson, and the genetics of speciation. Genetics 144, 1331–1335 (1996).

70. P. W. Messer, D. A. Petrov, Population genomics of rapid adaptation by soft selective sweeps. Trends Ecol Evol 28, 659–669 (2013).

71. R. E. Green et al., A draft sequence of the Neandertal genome. Science 328, 710–722 (2010).

72. A. B. Wolf, J. M. Akey, Outstanding questions in the study of archaic hominin admixture. PLoS Genet 14, e1007349 (2018).

73. D. Reich et al., Genetic history of an archaic hominin group from Denisova Cave in Siberia. Nature 468, 1053–1060 (2010).

74. M. Kuhlwilm et al., Ancient gene flow from early modern humans into Eastern Neanderthals. Nature 530, 429–433 (2016).

75. Q. Fu et al., The genetic history of Ice Age Europe. Nature 534, 200–205 (2016).

76. E. Huerta-Sanchez et al., Altitude adaptation in Tibetans caused by introgression of Denisovan-like DNA. Nature 512, 194–197 (2014).

77. R. M. Gittelman et al., Archaic Hominin Admixture Facilitated Adaptation to Out-of-Africa Environments. Curr Biol 26, 3375–3382 (2016).

78. O. Dolgova, O. Lao, Evolutionary and Medical Consequences of Archaic Introgression into Modern Human Genomes. Genes (Basel) 9, (2018).

79. D. Enard, D. A. Petrov, Evidence that RNA Viruses Drove Adaptive Introgression between Neanderthals and Modern Humans. Cell 175, 360–371 e313 (2018).

80. A. M. Bolger, M. Lohse, B. Usadel, Trimmomatic: a flexible trimmer for Illumina sequence data. Bioinformatics 30, 2114–2120 (2014).

81. D. R. Kelley, M. C. Schatz, S. L. Salzberg, Quake: quality-aware detection and correction of sequencing errors. Genome biology 11, R116 (2010).

82. R. Kajitani et al., Efficient de novo assembly of highly heterozygous genomes from whole-genome shotgun short reads. Genome research 24, 1384–1395 (2014).

83. A. F. A. Smit, R. Hubley, RepeatModeler Open-1.0. <http://www.repeatmasker.org>. (2008-2015).

84. H. Li, R. Durbin, Fast and accurate short read alignment with Burrows-Wheeler transform. Bioinformatics 25, 1754–1760 (2009).

85. W. Bao, K. K. Kojima, O. Kohany, Repbase Update, a database of repetitive elements in eukaryotic genomes. Mob DNA 6, 11 (2015).

86. A. F. A. Smit, R. Hubley, P. Green, RepeatMasker Open-4.0. <http://www.repeatmasker.org>. (2013-2015).

87. M. Kawamoto et al., High-quality genome assembly of the silkworm, *Bombyx mori*. Insect Biochem Mol Biol 107, 53–62 (2019).

88. G. dos Santos et al., FlyBase: introduction of the *Drosophila melanogaster* Release 6 reference genome assembly and large-scale migration of genome annotations. Nucleic Acids Res 43, D690–697 (2015).

89. G. S. Slater, E. Birney, Automated generation of heuristics for biological sequence comparison. BMC bioinformatics 6, 31 (2005).

90. C. Trapnell, L. Pachter, S. L. Salzberg, TopHat: discovering splice junctions with RNA-Seq. Bioinformatics 25, 1105–1111 (2009).

91. C. Trapnell et al., Transcript assembly and quantification by RNA-Seq reveals unannotated transcripts and isoform switching during cell differentiation. Nat Biotechnol 28, 511–515 (2010).

92. M. Stanke, R. Steinkamp, S. Waack, B. Morgenstern, AUGUSTUS: a web server for gene finding in eukaryotes. Nucleic Acids Res 32, W309–312 (2004).

93. A. Lomsadze, V. Ter-Hovhannisyan, Y. O. Chernoff, M. Borodovsky, Gene identification in novel eukaryotic genomes by self-training algorithm. Nucleic Acids Res 33, 6494–6506 (2005).

94. I. Korf, Gene finding in novel genomes. BMC bioinformatics 5, 59 (2004).

95. B. J. Haas et al., Automated eukaryotic gene structure annotation using EVidenceModeler and the Program to Assemble Spliced Alignments. Genome biology 9, R7 (2008).

96. J. Thurmond et al., FlyBase 2.0: the next generation. Nucleic Acids Res 47, D759–D765 (2019).

97. C. UniProt, UniProt: a worldwide hub of protein knowledge. Nucleic Acids Res 47, D506–D515 (2019).

98. C. Gene Ontology, Gene Ontology Consortium: going forward. Nucleic Acids Res 43, D1049–1056 (2015).

99. W. J. Kent, R. Baertsch, A. Hinrichs, W. Miller, D. Haussler, Evolution’s cauldron: duplication, deletion, and rearrangement in the mouse and human genomes. Proc Natl Acad Sci U S A 100, 11484–11489 (2003).

100. M. Hiller et al., Computational methods to detect conserved non-genic elements in phylogenetically isolated genomes: application to zebrafish. Nucleic Acids Res 41, e151 (2013).

101. M. Blanchette et al., Aligning multiple genomic sequences with the threaded blockset aligner. Genome research 14, 708–715 (2004).

102. S. F. Altschul, W. Gish, W. Miller, E. W. Myers, D. J. Lipman, Basic local alignment search tool. Journal of molecular biology 215, 403–410 (1990).

103. Y. Benjamini, Y. Hochberg, Controlling the False Discovery Rate: A Practical and Powerful Approach to Multiple Testing. Journal of the Royal Statistical Society. Series B (Methodological) 57, 289–300 (1995).

104. L. Li, C. J. Stoeckert, Jr., D. S. Roos, OrthoMCL: identification of ortholog groups for eukaryotic genomes. Genome research 13, 2178–2189 (2003).

105. B. Buchfink, C. Xie, D. H. Huson, Fast and sensitive protein alignment using DIAMOND. Nat Methods 12, 59–60 (2015).

106. C. Fraisse, M. A. L. Picard, B. Vicoso, The deep conservation of the Lepidoptera Z chromosome suggests a non-canonical origin of the W. Nat Commun 8, 1486 (2017).

107. L. T. Nguyen, H. A. Schmidt, A. von Haeseler, B. Q. Minh, IQ-TREE: a fast and effective stochastic algorithm for estimating maximum-likelihood phylogenies. Mol Biol Evol 32, 268–274 (2015).

108. J. Sukumaran, M. T. Holder, DendroPy: a Python library for phylogenetic computing. Bioinformatics 26, 1569–1571 (2010).

109. J. Huerta-Cepas, F. Serra, P. Bork, ETE 3: Reconstruction, Analysis, and Visualization of Phylogenomic Data. Mol Biol Evol 33, 1635–1638 (2016).

110. J. Zhang, Q. Cong, J. Shen, E. Brockmann, N. V. Grishin, Genomes reveal drastic and recurrent phenotypic divergence in firetip skipper butterflies (Hesperiidae: Pyrrhopyginae). Proceedings. Biological sciences 286, 20190609 (2019).

