## Supplementary figures and images for "Genomics of a complete butterfly continent"

### Figure 2

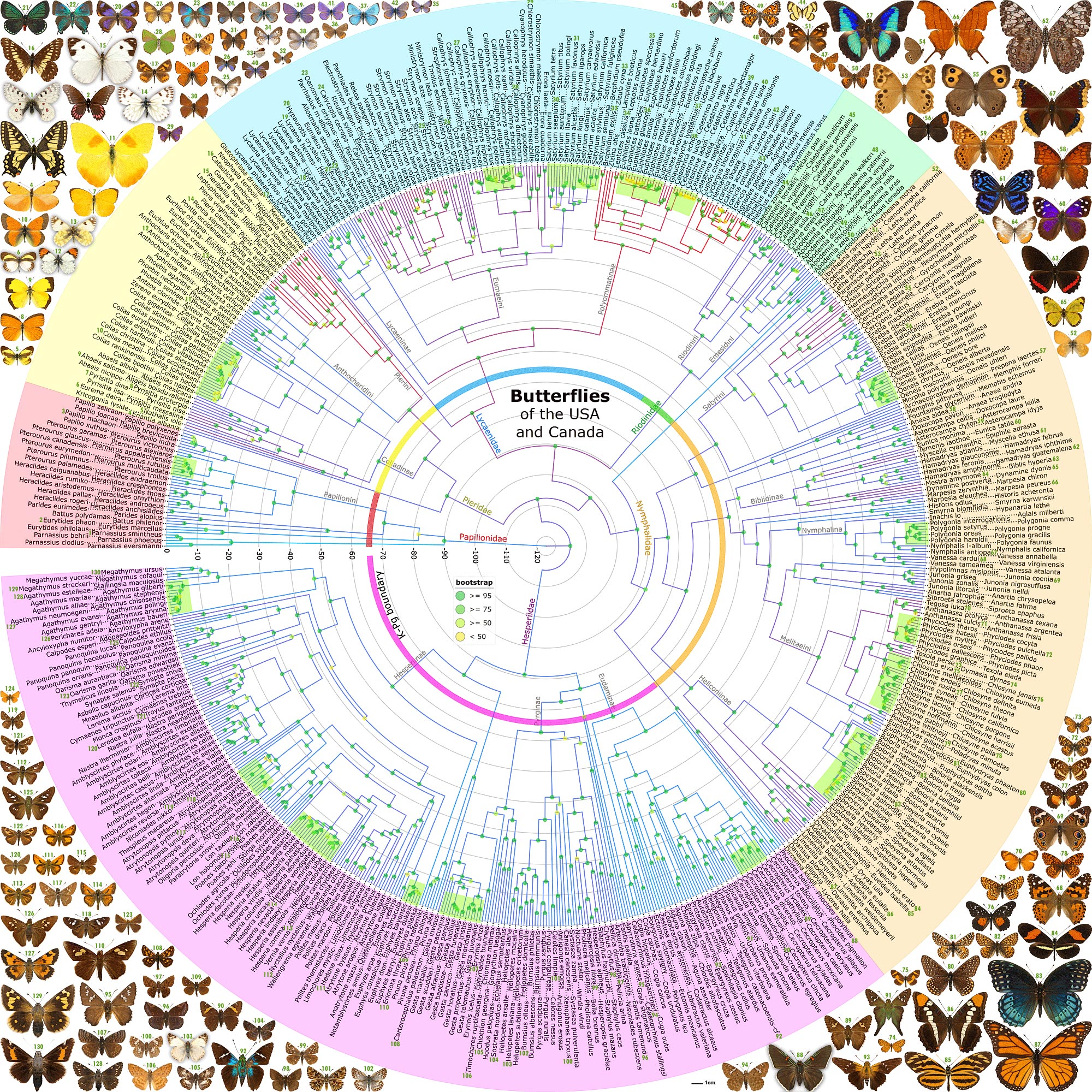

### Supplemental Figure S1

Nuclear genome tree

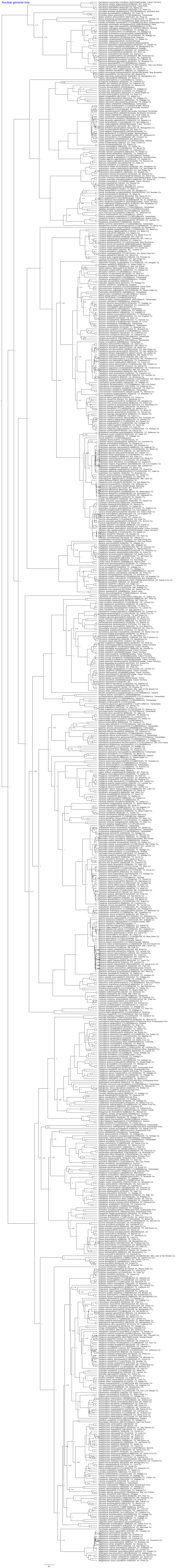

Z chromosome tree

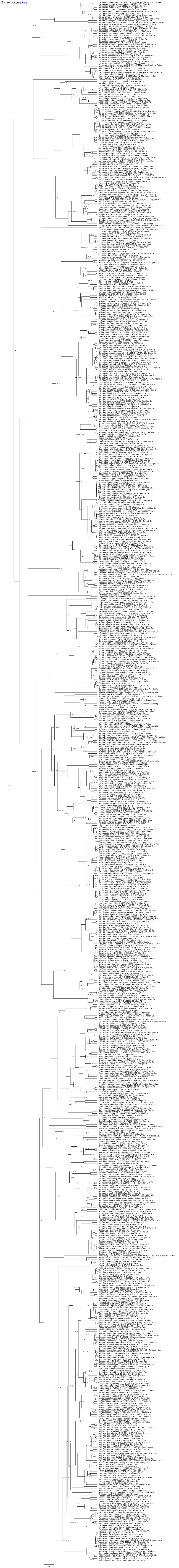

Mitochondrial genome tree

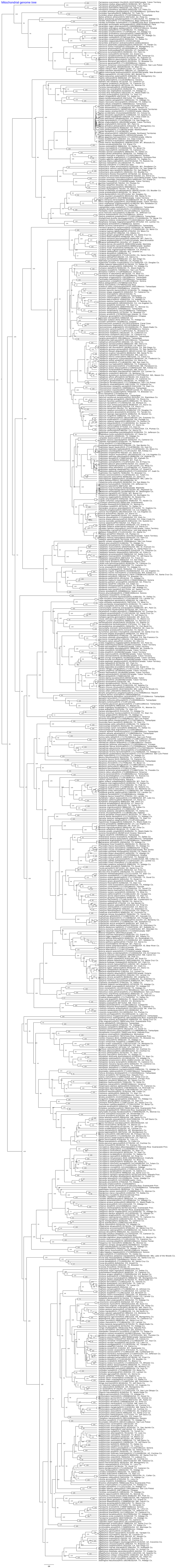
