## Supplemental Tables S1 for "Genomics of a complete butterfly continent"

### Table S1. Data for the butterfly specimens of 845 species we sequenced and the outgroup

Species are ordered according to our suggested taxonomic sequence that is suggested by the analysis of phylogenetic trees. The authors and dates of the taxa are not provided and can be found in the Pelham catalogue (2019), along with the synonymy. The names of genera applied to these species reflect our recently published results (Cong et al. 2019; Zhang et al. 2019). Names of genera recently described as new in connection with this study are highlighted in green, and those used in revised genus-species combinations are highlighted in yellow.

| # | DNA Voucher | Taxon name | Type | Sex | Locality and date | Repository | Genitalia # | Collection # |
|---|---|---|---|---|---|---|---|---|
| 1 | NVG-16107A06 | *Parnassius eversmanni thor* | | M | Canada: Yukon Territory, Montana Mountain, 24-Jun-2016 | WDempwolf | | |
| 2 | NVG-9436 | *Parnassius clodius altaurus* | | F | USA: WY, Park Co., Yellowstone National Park, 23-Jul-2017 | | | |
| 3 | PAO-E12 | *Parnassius phoebus* | | | Switzerland: Furka Pass, 27-Jul-2017 | | | |
| 4 | NVG-6662 | *Parnassius behrii* | | M | USA: CA, Alpine Co., Ebbetts Pass, 6-Jul-2015 | | | |
| 5 | NVG-6592 | *Parnassius smintheus sayii* | | M | USA: CO, Pitkin Co., White River National Forest, 11-Jul-2016 | | | |
| 6 | NVG-15112E04 | *Eurytides philolaus philolaus* | | M | Mexico: Quintana Roo, Tulun, 19-Apr-1990 | FMNH | | |
| 7 | NVG-3488 | *Eurytides marcellus* | | | USA: TX, Tyler Co., Kirby State Forest, 7-Jun-2015 | | | |
| 8 | NVG-17116F05 | *Eurytides phaon phaon* | | M | Mexico: Tamaulipas, 21-Feb-1975 | TAMU | | |
| 9 | NVG-5937 | *Battus philenor philenor* | | F | USA: AZ, Pima Co., Santa Rita Mts., 29-Mar-2016 | | | |
| 10 | NVG-17116E12 | *Battus polydamas polydamas* | | M | USA: TX, Hidalgo Co., 9-Sep-1972 | TAMU | | |
| 11 | NVG-17114C04 | *Parides alopius* | | M | Mexico: Baja California Sur, 27-Apr-1970 | CSUC | | CSU_ENT1004387 |
| 12 | NVG-17119D06 | *Parides eurimedes mylotes* | | | Costa Rica: Guanacaste Prov., ACG, Sector Santa Rosa, 2009 | USNM | | 09-SRNP-14759 |
| 13 | NVG-14081G03 | *Heraclides anchisiades anchisiades* | | F | Brazil: Para, Obidos, 3-Aug-1984 | CSUC | | |
| 14 | NVG-14082B06 | *Heraclides rogeri pharnaces* | | M | Mexico: El Rio, 10-Aug-1985 | CSUC | | |
| 15 | NVG-14081C07 | *Heraclides androgeus epidaurus* | | F | Mexico: Sinaloa, Chirimollos, Hwy 40, 23-Nov-2-Dec-2003 | CSUC | | |
| 16 | NVG-17116F08 | *Heraclides pallas* | | M | USA: TX, Hidalgo Co., 26-Jun-1975 | TAMU | | |
| 17 | NVG-17116F06 | *Heraclides ornythion* | | M | USA: TX, Hidalgo Co., 26-Jun-1975 | TAMU | | |
| 18 | NVG-14065C06 | *Heraclides aristodemus ponceana* | | M | USA: FL, Key Largo, 7-May-1972 | JAScott | | |
| 19 | NVG-14112C06 | *Heraclides thoas autocles* | | F | USA: TX, Hidalgo Co., Santa Ana NWR, 30-Jul-1972 | TAMU | | |
| 20 | NVG-2278 | *Heraclides rumiko* | PT | M | Costa Rica: Alajuela, Alajuela Adventist University, 15-20-May-1995 | TMMC | NVG140403-06 | |
| 21 | NVG-6035 | *Heraclides cresphontes* | | M | USA: TX, Dallas Co., Dallas, 7-Apr-2016 | | | |
| 22 | NVG-14102A10 | *Heraclides caiguanabus* | | | Cuba, old, before 1900 | FMNH | | |
| 23 | NVG-10301 | *Heraclides andraemon andraemon* | | F | Jamaica, 3-Dec-2017 | | | |
| 24 | NVG-6053 | *Pterourus palamedes palamedes* | | F | USA: TX, Hardin Co., FM1003 0.25 mi N of FM770, 24-Apr-2016 | | | |
| 25 | NVG-4262 | *Pterourus troilus troilus* | | F | USA: IN, Montgomery Co., 1.5 mi south of Deer Mill, 1-Aug-2015 | | | |
| 26 | NVG-16104H04 | *Pterourus pilumnus* | | F | Mexico: Hidalgo, 3-Mar-1978 | LACM | | |
| 27 | PAO-182 | *Pterourus multicaudata* | | | USA: UT, Garfield Co., Escalante, 4-Aug-2016 | | | |
| 28 | NVG-6526 | *Pterourus eurymedon* | | F | USA: CO, Park Co., 4.5 air mi SSE Florissant, 9-Jul-2016 | | | |
| 29 | NVG-6374 | *Pterourus rutulus* | | M | USA: CO, Grand Co., 13.5 air mi N Silverthorne, 5-Jul-2016 | | | |
| 30 | NVG-15116F05 | *Pterourus canadensis* | | M | USA: ME, Somerset Co., Hwy 201, 12-Jun-1993 | CSUC | | CSU_ENT1004548 |
| 31 | NVG-6172 | *Pterourus appalachiensis* | | M | USA: VA, Augusta Co., Calfpasture River, 5 mi SSW Deerfield, 11-May-2016 | | | |
| 32 | NVG-1670 | *Pterourus glaucus glaucus* | | M | USA: TX, Denton Co., Lake Ray Roberts State Park, 4-Aug-2013 | USNM | | |
| 33 | NVG-5522 | *Pterourus alexiares garcia* | | M | USA: TX, Brewster Co., Big Bend National Park, 12-Jun-2004 | TAMU | NVG160110-59 | |
| 34 | NVG-17113D03 | *Pterourus garamas* | | | Mexico: Oaxaca, 1988 | TMMC | | |
| 35 | NVG-17116F11 | *Pterourus victorinus victorinus* | | M | Mexico: San Luis Potosi, 21-Feb-1976 | TAMU | | |
| 36 | NVG-18042E04 | *Papilio xuthus* | | M | USA: HI, Maui Co., Kula near intersection Ainakula Rd and Hwy 377, 20-Mar-2004 | WDempwolf | | WRD 2781 |
| 37 | 11-BOA-13384A11 | *Papilio indra* | | M | USA: UT, Cache Co., Hyram, Left Hand Blacksmith Fork Canyon, 16-Jun-2011 | | | |
| 38 | PAO-188 | *Papilio machaon bairdii* | | | USA: UT, Garfield Co., Grand Staircase - Escalante National Monument, 5-Aug-2016 | | | |
| 39 | NVG-16104C09 | *Papilio brevicauda bretonensis* | | F | Canada: New Bruswick, 26-Mar-1953 | LACM | | |
| 40 | NVG-15116E11 | *Papilio joanae* | | M | USA: MO, Benton Co., near Warsaw, 16-May-1975 | CSUC | | CSU_ENT1006307 |
| 41 | NVG-4265 | *Papilio polyxenes asterius* | | M | USA: IN, Montgomery Co., 1.5 mi south of Deer Mill, 1-Aug-2015 | | | |
| 42 | NVG-9555 | *Papilio zelicaon* | | M | USA: UT, Davis Co., Wasatch National Forest, 28-Jul-2017 | | | |
| 43 | NVG-17114C05 | *Enantia albania* | | M | Mexico: Oaxaca, Candelaria Loxicha, 21-Oct-1983 | CSUC | | |
| 44 | NVG-3423 | *Kricogonia lyside* | | F | USA: TX, Cameron Co., 3 mi SW Sebastian, 30-May-2015 | | | |
| 45 | NVG-3572 | *Nathalis iole iole* | | F | USA: TX, Cameron Co., 3 mi SW Sebastian, 13-Jun-2015 | | | |
| 46 | NVG-4872 | *Eurema daira daira* | | | USA: FL, Monroe Co., Key West, 3-Oct-2015 | | | |
| 47 | NVG-10243 | *Pyrisitia messalina* | | | Jamaica, 2-Dec-2017 | | | |
| 48 | NVG-3545 | *Pyrisitia lisa centralis* | | F | USA: TX, Hidalgo Co., Old Rio Rico Rd. 1.5 air mi SE of Relampago, 13-Jun-2015 | | | |
| 49 | NVG-5142 | *Pyrisitia nise nelphe* | | M | USA: TX, Hidalgo Co., Chihuahua RR tracks, 15-Nov-2015 | | | |

| | | | | | | | | |
|---|---|---|---|---|---|---|---|---|
| 50 | NVG-5351 | *Pyrisitia dina helios* | | M | USA: FL, Miami-Dade Co., Homestead, 20-Dec-2015 | | | |
| 51 | NVG-5167 | *Pyrisitia proterpia* | | M | USA: TX, Starr Co., Rio Grande City, 15-Nov-2015 | | | |
| 52 | NVG-4057 | *Abaeis nicippe* | | M | USA: TX, Dallas Co., Dallas, Norbuck Park, 14-Jul-2015 | | | |
| 53 | NVG-3357 | *Abaeis boisduvaliana* | | M | USA: TX, Hidalgo Co., Old Rio Rico Rd. 1.5 air mi SE of Relampago, 23-May-2015 | | | |
| 54 | NVG-10664 | *Abaeis salome jamapa* | | M | USA: TX, Hidalgo Co., Edinburd, 5-Sep-1972 | TAMU | NVG180106-81 | |
| 55 | NVG-4202 | *Abaeis mexicana mexicana* | | F | USA: TX, Dallas Co., Dallas, Norbuck Park, 21-Jul-2015 | | | |
| 56 | NVG-10660 | *Abaeis albula celata* | | F | Mexico: Tamaulipas, 12-Dec-1975 | TAMU | NVG180106-77 | |
| 57 | NVG-17114D02 | *Colias nastes streckeri* | | M | Canada: Alberta, Division No. 15, 21-Jul-2003 | CSUC | | CSU_ENT1042199 |
| 58 | NVG-17114C12 | *Colias boothii boothii* | | M | Canada: Nunavut, 11-17-Jul-1988 | CSUC | | CSU_ENT1042144 |
| 59 | NVG-17114C09 | *Colias hecla* | | M | USA: AK, Yukon-Koyukuk, Eagle Summit, 2-Jul-1979 | CSUC | | CSU_ENT1042132 |
| 60 | NVG-19035F10 | *Colias rankinensis* | | M | Canada: Northwest Territories, Schultz Lake Rapids, 28-Jul-1965 | USNM | | |
| 61 | NVG-19062H07 | *Colias johanseni* | PT | F | Canada: Northwest Territories, District of Mackenzie, Bernard Harbour, 11-17-Jul-1988 | LACM | | |
| 62 | PAO-159 | *Colias meadii meadii* | | | USA: CO, Clear Creek Co., Loveland Pass, 1-Aug-2016 | | | |
| 63 | GK6 | *Colias occidentalis chrysomelas* | | | USA: CA, Colusa Co., Mill Valley Camp, 7-Jun-2017 | | | |
| 64 | NVG-19039A05 | *Colias christina krauthii* | | M | USA: SD, Black Hills, 2-Jul-1986 | USNM | | |
| 65 | NVG-11560 | *Colias alexandra apache* | | M | USA: AZ, Apache Co., 25-May-2018 | | | |
| 66 | NVG-17114C06 | *Colias harfordii* | | M | USA: CA, San Diego Co., 22-Jul-2008 | CSUC | | CSU_ENT1040534 |
| 67 | NVG-18042E08 | *Colias vitabunda* | | F | Canada: Yukon Territory, Dempster Highway, km 131, 5-Jul-2016 | WDempwolf | | WRD 11012 |
| 68 | NVG-9421 | *Colias eriphyle* | | M | USA: WY, Park Co., Yellowstone National Park, 23-Jul-2017 | | | |
| 69 | NVG-4261 | *Colias philodice philodice* | | M | USA: IN, Montgomery Co., 1.5 mi south of Deer Mill, 1-Aug-2015 | | | |
| 70 | NVG-6530 | *Colias eurytheme* | | F | USA: CO, Park Co., 4.5 air mi SSE Florissant, 9-Jul-2016 | | | |
| 71 | NVG-19034A10 | *Colias palaeno chippewa* | | M | Canada: YT, Milepost 1024 Hwy 1, 4-Jul-1982 | USNM | | |
| 72 | NVG-17119E03 | *Colias pelidne* | | M | Canada: Newfoundland, Hwy 1, 2.8 mi SW Mummichog Park, 8-Jul-1975 | USNM | | |
| 73 | NVG-17114D06 | *Colias skinneri* | | F | USA: WY, Fremont Co., 30-Jul-2007 | CSUC | | CSU_ENT1041861 |
| 74 | NVG-18062B11 | *Colias interior dornfeldi* | | M | USA: MT, Missoula Co., 28-Jun-1994 | USNM | | |
| 75 | NVG-18038A05 | *Colias behrii* | | M | USA: CA, Mono Co., 19-Jul-2009 | CSUC | | |
| 76 | NVG-17114D04 | *Colias gigantea harroweri* | | M | Canada: Alberta, Division No. 15, 19-Jul-2003 | CSUC | | CSU_ENT1041736 |
| 77 | PAO-160 | *Colias scudderii* | | | USA: CO, Clear Creek Co., Loveland Pass, 1-Aug-2016 | | | |
| 78 | GK8 | *Zerene eurydice* | | | USA: CA, Placer Co., Placer land Trust, 28-Jun-2017 | | | |
| 79 | NVG-3980 | *Zerene cesonia* | | F | USA: TX, Dallas Co., Dallas, Norbuck Park, 8-Jul-2015 | | | |
| 80 | NVG-17116H03 | *Anteos clorinde* | | F | USA: TX, Bexar Co., 29-Jul-1986 | TAMU | | |
| 81 | NVG-10234 | *Anteos maerula* | | F | USA: TX, Cameron Co., Harlingen, 14-Oct-2017 | | | |
| 82 | NVG-3950 | *Phoebis sennae eubule* | | M | USA: TX, Dallas Co., Coppell, 5-Jul-2015 | | | |
| 83 | NVG-3829 | *Phoebis philea philea* | | M | USA: TX, Hidalgo Co., Old Rio Rico Rd. 1.5 air mi SE of Relampago, 27-Jun-2015 | | | |
| 84 | NVG-17114D07 | *Phoebis neocypris virgo* | | M | Mexico: Sinaloa, Concordia, 23-Nov-2-Dec-2003 | CSUC | | CSU_ENT1049496 |
| 85 | NVG-17116H05 | *Phoebis argante argante* | | M | Mexico: Quintana Roo, 24-Oct-1979 | TAMU | | |
| 86 | NVG-4925 | *Phoebis agarithe agarithe* | | M | USA: TX, Zapata Co., Morales-Sanchez, 12-Nov-2015 | | | |
| 87 | NVG-16107D12 | *Aphrissa statira statira* | | M | USA: TX, Nueces Co., Blucher Park, 5-Aug-2013 | JMcDermott | | |
| 88 | NVG-14114E02 | *Aphrissa neleis* | | M | USA: FL, Dade Co., Homestead, 18-Jul-2013 | TLS | | |
| 89 | NVG-17119E07 | *Aphrissa orbis* | | F | no data, old | USNM | | |
| 90 | NVG-6134 | *Anthocharis midea annickae* | | M | USA: WV, Pendleton Co., FS112, 8-May-2016 | | | |
| 91 | NVG-6667 | *Anthocharis cethura cethura* | | M | USA: CA, San Diego Co., 6-Mar-2004 | | | |
| 92 | NVG-5638 | *Anthocharis sara sara* | | | USA: CA, San Diego Co., 11-Feb-2016 | | | |
| 93 | NVG-1952 | *Anthocharis julia julia* | | M | USA: CO, Boulder Co., 1.7 mi WSW Univeristy of Colorado, 22-Apr-2006 | | | |
| 94 | NVG-1970 | *Anthocharis thoosa thoosa* | | M | USA: AZ, Mohave Co., Virgin Mountains, 14-Mar-2008 | | | |
| 95 | PAO-63 | *Anthocharis lanceolata lanceolata* | | | USA: CA, Sierra Co., Tahoe National Forest, 19-Jun-2016 | | | |
| 96 | NVG-16107B05 | *Euchloe ochracea kamchatkensis* | | M | Canada: Yukon Territory, Dempster Hwy km 131 (west), 6-Jul-2016 | WDempwolf | | |
| 97 | NVG-6465 | *Euchloe ausonides coloradensis* | | M | USA: CO, Grand Co., 4.3 air mi SE Hot Sulphur Springs, 6-Jul-2016 | | | |
| 98 | NVG-17114D09 | *Euchloe creusa* | | M | Canada: Yukon Territory, Ogilvie Mtns., Windy Pass, 28-Jun-2003 | CSUC | | CSU_ENT1042040 |
| 99 | PAO-70 | *Euchloe hyantis* | | | USA: CA, Plumas Co., Plumas National Forest, 19-Jun-2016 | | | |
| 100 | PAO-391 | *Euchloe lotta* | | | USA: NV, Elko Co., IH80, 14-Jun-2017 | | | |
| 101 | NVG-6663 | *Euchloe guaymasensis* | | M | Mexico: Sonora, Camino Al Panteon, 29-Jan-2012 | | | |
| 102 | NVG-74 | *Euchloe olympia* | | F | USA: TX, Wise Co., LBJ National Grassland, 16-Mar-2004 | | | |
| 103 | NVG-15112F11 | *Pontia beckerii* | | F | USA: NV, Douglas Co., 19-Aug-1993 | FMNH | | |
| 104 | NVG-18042E12 | *Pontia occidentalis occidentalis* | | F | USA: CO, Boulder Co., Haystack Mountain, 2-Jun-1957 | WDempwolf | | WRD 4080 |
| 105 | NVG-8433 | *Pontia protodice* | | F | USA: TX, Wise Co., LBJ National Grassland, 4-Apr-2017 | | | |

| | | | | | | | | |
|---|---|---|---|---|---|---|---|---|
| 106 | NVG-8922 | *Pontia sisymbrii elivata* | | F | USA: NM, Santa Fe Co., Santa Fe National Forest, 18-May-2017 | | | |
| 107 | NVG-16107B06 | *Pieris angelika* | | M | Canada: Yukon Territory, Dempster Hwy km 151, 6-Jul-2016 | WDempwolf | | |
| 108 | NVG-16105B04 | *Pieris oleracea oleracea* | | M | USA: MI, St. Joseph Co., 1-Mar-1984 | LACM | | |
| 109 | NVG-16107B09 | *Pieris marginalis guppyi* | | M | Canada: Yukon Territory, Montana Mountain, 24-Jun-2016 | WDempwolf | | |
| 110 | NVG-6138 | *Pieris virginiensis virginiensis* | | M | USA: WV, Pendleton Co., FS274, 8-May-2016 | | | |
| 111 | NVG-3537 | *Pieris rapae* | | F | USA: TX, Dallas Co., Dallas, 5-Jun-2015 | | | |
| 112 | NVG-17116G10 | *Leptophobia aripa elodia* | | M | Mexico: Tamaulipas, 14-Feb-1980 | TAMU | | |
| 113 | NVG-17116H09 | *Itaballia demophile centralis* | | F | Venezuela, 30-Oct-1980 | TAMU | | |
| 114 | NVG-15112F10 | *Pieriballia viardi viardi* | | M | Mexico: Guerrero, vic. Ixtapa-Zihautenejo, 12-Jan-1984 | FMNH | | |
| 115 | NVG-4148 | *Ascia monuste monuste* | | F | USA: TX, Jefferson Co., S of Sabine Pass, 18-Jul-2015 | | | |
| 116 | NVG-15117H10 | *Ganyra howarthi* | | F | Mexico: Sonora, 2 mi E San Carlos, 29-Jan-2003 | CSUC | | |
| 117 | NVG-17116G12 | *Ganyra josephina josepha* | | F | Mexico: Tamaulipas, 22-Jan-1974 | TAMU | | |
| 118 | NVG-17116G01 | *Catasticta nimbice nimbice* | | M | Mexico: Nuevo Leon, 23-Oct-1979 | TAMU | | |
| 119 | PAO-257 | *Neophasia menapia magnamenapia* | | | USA: CO, Larimer Co., Roosevelt National Forest, 10-Sep-2016 | | | |
| 120 | NVG-2768 | *Neophasia terlooii* | | M | Mexico: Sonora, 10.9 miles S Huachinera, 4-Jul-1979 | CSUC | | |
| 121 | NVG-17116G03 | *Melete lycimnia isandra* | | M | Mexico: Tamaulipas, 21-Nov-1974 | TAMU | | |
| 122 | NVG-17116G06 | *Glutophrissa drusilla neumoegenii* | | F | USA: TX, Bexar Co., 31-Oct-1996 | TAMU | | |
| 123 | NVG-4334 | *Feniseca tarquinius tarquinius* | | M | USA: IN, Newton Co., 2.5 mi S Lake Village, 2-Aug-2015 | | | |
| 124 | NVG-17115H02 | *Lycaena phlaeas hypophlaeas* | | M | USA: CA, Mono Co., Sonora Pass, 12-Jul-1994 | CSUC | | |
| 125 | PAO-475 | *Lycaena cupreus lapidicola* | | | USA: CA, Mono Co., Mosquito flat, 6-Jul-2017 | | | |
| 126 | PAO-286 | *Lycaena hyllus* | | | USA: CO, Adams Co., Barr Lake State Park, 8-Oct-2016 | | | |
| 127 | PAO-346 | *Lycaena gorgon gorgon* | | | USA: CA, Santa Cruz Co., Santa Cruz Mountains, 16-May-2017 | | | |
| 128 | NVG-9348 | *Lycaena heteronea* | | F | USA: WY, Park Co., Yellowstone National Park, 24-Jul-2017 | | | |
| 129 | NVG-9440 | *Lycaena mariposa penroseae* | | M | USA: WY, Park Co., Yellowstone National Park, 23-Jul-2017 | | | |
| 130 | PAO-191 | *Lycaena helloides* | | | USA: UT, Grand Co., E Moab, 7-Aug-2016 | | | |
| 131 | NVG-17114D12 | *Lycaena dospassosi* | | F | Canada: New Brunswick, Bathurst, 5-Aug-1983 | CSUC | | |
| 132 | NVG-19081G05 | *Lycaena dorcas dorcas* | | M | Canada: Manitoba, Pine Ridge, 30-Jul-1979 | HPavulaan | | |
| 133 | NVG-6686 | *Lycaena epixanthe epixanthe* | | M | Canada: Quebec, Laurentides, 17-Jul-2011 | | | |
| 134 | NVG-6459 | *Lycaena nivalis browni* | | M | USA: CO, Grand Co., 4 air mi SSE Hot Sulphur Springs, 6-Jul-2016 | | | |
| 135 | PAO-126 | *Lycaena rubidus* | | | USA: WY, Laramie Co., Little Bear community, 16-Jul-2016 | | | |
| 136 | NVG-10599 | *Lycaena dione* | | M | USA: TX, 31-May-1977 | TAMU | NVG180106-16 | |
| 137 | NVG-17114D11 | *Lycaena xanthoides* | | M | USA: CA, Santa Clara Co., East of Mt. Hamilton, 18-Jul-2007 | CSUC | | |
| 138 | PAO-60 | *Lycaena editha* | | | USA: CA, Sierra Co., Tahoe National Forest, 18-Jun-2016 | | | |
| 139 | PAO-449 | *Lycaena arota virginiensis* | | | USA: CA, Plumas Co., Fether River north of Clio, 3-Jul-2017 | | | |
| 140 | NVG-6685 | *Lycaena hermes* | | M | USA: CA, San Diego Co., Roberts Ranch, 1-Jul-2011 | | | |
| 141 | PAO-466 | *Habrodais grunus* | | | USA: CA, Plumas Co., 2 mi E Twain, 4-Jul-2017 | | | |
| 142 | NVG-17117A05 | *Hypaurotis crysalus crysalus* | | M | USA: CO, Douglas Co., 28-Apr-1993 | TAMU | | |
| 143 | NVG-5636 | *Eumaeus atala* | | F | USA: FL, Miami-Dade Co., Miami, 9-Feb-2016 | | | |
| 144 | NVG-7284 | *Eumaeus toxea* | | F | Mexico: San Luis Potosi, 10 mi SW Tamazunchale, 12-Aug-1972 | USNM | NVG161007-11 | |
| 145 | NVG-9179 | *Atlides halesus halesus* | | F | USA: TX, Dallas Co., Dallas, 9-Jun-2017 | | | |
| 146 | NVG-10604 | *Allosmaitia strophius* | | M | USA: TX, Hidalgo Co., Madero, 14-Nov-1993 | TAMU | NVG180106-21 | |
| 147 | NVG-17119F05 | *Michaelus ira* | | | Mexico: Veracruz, 21-Jul-1-Aug-1985 | USNM | | |
| 148 | NVG-8673 | *Parrhasius m-album* | | F | USA: TX, Wise Co., LBJ National Grassland, 7-May-2017 | | | |
| 149 | NVG-10623 | *Parrhasius moctezuma* | | M | Mexico: Nuevo Leon, 26-Oct-1979 | TAMU | NVG180106-40 | |
| 150 | NVG-10615 | *Oenomaus ortygnus* | | M | Mexico: Tamaulipas, 21-Feb-1974 | TAMU | NVG180106-32 | |
| 151 | NVG-15115H06 | *Pendantus guzanta* | | F | Mexico: Chiapas, Oxchuc, 20-25 Sep 1975 | USNM | | USNMENT00149445 |
| 152 | NVG-3306 | *Calycopis cecrops* | | F | USA: LA, Natchitoches Pa, Kisatchie NF, 12-Apr-2015 | | | |
| 153 | NVG-3431 | *Calycopis isobeon* | | M | USA: TX, Hidalgo Co., Penitas RR tracks, 30-May-2015 | | | |
| 154 | NVG-7423 | *Kisutam syllis* | | M | Mexico: Chiapas, 10 km S. San Cristobal d. l. Casas, 21-May-1981 | USNM | NVG161105-96 | |
| 155 | NVG-15115F07 | *Electrostrymon joya* | | M | Panama: Canal Zone, 27-Feb-1979 | USNM | | USNMENT00149422 |
| 156 | NVG-15115F02 | *Electrostrymon hugon* | | M | Belize: Cayo, Macaw River , 27-Jun-2003 | USNM | | USNMENT00149417 |
| 157 | NVG-15115G04 | *Electrostrymon angelia* | | M | USA: FL, Miami-Dade Co., Homestead, 19-May-1982 | USNM | | USNMENT00149431 |
| 158 | NVG-10625 | *Panthiades bathildis* | | M | Mexico: Tamaulipas, 8-Jan-1974 | TAMU | NVG180106-42 | |
| 159 | NVG-17114E12 | *Hypostrymon critola* | | F | Mexico: Sonora, 24-Mar-2004 | CSUC | | |
| 160 | NVG-17076B03 | *Rekoa palegon* | | M | Costa Rica | USNM | | USNMENT00181220 |
| 161 | NVG-17076B04 | *Rekoa marius* | | F | Costa Rica | USNM | | USNMENT00181221 |

| | | | | | | | | |
|---|---|---|---|---|---|---|---|---|
| 162 | NVG-10621 | *Arawacus jada (=Dolymorpha)* | | F | Mexico: Tamaulipas, 6-May-1978 | TAMU | NVG180106-38 | |
| 163 | NVG-10593 | *Strymon cestri* | | M | Mexico: Veracruz, 25-Mar-1977 | TAMU | NVG180106-10 | |
| 164 | NVG-5216 | *Strymon bazochii bazochii* | | F | USA: TX, Hidalgo Co., Penitas RR tracks, 25-Nov-2015 | | | |
| 165 | NVG-5264 | *Strymon istapa istapa* | | M | USA: TX, Cameron Co., Port Isabel, 27-Nov-2015 | | | |
| 166 | NVG-18014C02 | *Strymon limenia* | | | USA: FL, no data, old | USNM | | |
| 167 | NVG-10591 | *Strymon yojoa* | | F | Mexico: Tamaulipas, 29-Dec-1974 | TAMU | NVG180106-08 | |
| 168 | NVG-10585 | *Strymon rufofusca* | | F | USA: TX, Hidalgo Co., Bentsen-Rio Frande Valley State Park, 28-Dec-1973 | TAMU | NVG180106-02 | |
| 169 | NVG-10584 | *Strymon bebrycia* | | F | USA: TX, Hidalgo Co., Santa Ana NWR, 1-Dec-1968 | TAMU | NVG180106-01 | |
| 170 | NVG-5334 | *Strymon melinus melinus* | | M | USA: FL, Monroe Co., Key West, 19-Dec-2015 | | | |
| 171 | NVG-18033G02 | *Strymon avalona* | | M | USA: CA, Los Angeles Co., Avalon, 12-Mar-2013 | MWalker | | |
| 172 | NVG-5242 | *Strymon alea* | | F | USA: TX, Starr Co., Rio Grande City, 26-Nov-2015 | | | |
| 173 | NVG-18064H05 | *Strymon martialis* | | M | USA: FL, Monroe Co., Big Pine Key, 14-May-1986 | USNM | | |
| 174 | NVG-17076E03 | *Strymon acis* | | F | Virgin Islands: St. Croix, | USNM | | USNMENT00181255 |
| 175 | NVG-10589 | *Strymon albata* | | F | Mexico: Tamaulipas, 2-Jan-1975 | TAMU | NVG180106-06 | |
| 176 | NVG-14112C08 | *Strymon serapio* | | | Mexico: Tamaulipas, | TAMU | | |
| 177 | NVG-1573 | *Strymon solitario* | | M | USA: TX, Brewster Co., Big Bend National Park, 26-Mar-2005 | | | |
| 178 | NVG-17119F07 | *Ministrymon azia* | | | USA: FL, Broward Co., Coral Springs, 24-Aug-1989 | USNM | | |
| 179 | NVG-17119F06 | *Ministrymon janevicroy* | | | Mexico: Sonora, 3-Dec-1995 | USNM | | |
| 180 | NVG-10610 | *Ministrymon leda* | | F | USA: TX, Culberson Co., 15 mi S of Van Horn, 17-Jul-1978 | TAMU | NVG180106-27 | |
| 181 | NVG-4021 | *Ministrymon clytie* | | F | USA: TX, Cameron Co., River Dr., 1 mi S of Santa Maria, 11-Jul-2015 | | | |
| 182 | NVG-10614 | *Tmolus echion echiolus* | | M | Mexico: Tamaulipas, 12-Feb-1974 | TAMU | NVG180106-31 | |
| 183 | NVG-10613 | *Gargina gnosia* | | M | Mexico: Tamaulipas, 7-Mar-1975 | TAMU | NVG180106-30 | |
| 184 | NVG-10612 | *Strephonota tephraeus* | | F | Mexico: Tamaulipas, 4-Dec-1974 | TAMU | NVG180106-29 | |
| 185 | NVG-10609 | *Ocaria ocrisia* | | F | Mexico: Tamaulipas, 19-Nov-1974 | TAMU | NVG180106-26 | |
| 186 | PAO-440 | *Callophrys johnsoni* | | | USA: CA, Sierra Co., FR54, 25-Jun-2017 | | | |
| 187 | NVG-8956 | *Callophrys spinetorum* | | F | USA: NM, Otero Co., Lincoln National Forest, 21-May-2017 | | | |
| 188 | NVG-18022C07 | *Callophrys muiri* | ST | M | USA: CA, Marin Co., Tamalpais, Jun-1880 | AMNH | | |
| 189 | NVG-18036G12 | *Callophrys loki thornei* | | F | USA: CA, San Diego Co., 9-Feb-2011 | MWalker | | |
| 190 | NVG-3521 | *Callophrys gryneus gryneus* | | M | USA: TX, San Jacinto Co., Sam Houston NF, 6.4 mi W Shepherd, 7-Jun-2015 | | | |
| 191 | NVG-18014C11 | *Callophrys hesseli angulata* | | | USA: NC, Gates Co., Great Dismal Swamp, 8-Jun-1983 | USNM | | |
| 192 | NVG-8849 | *Callophrys eryphon fusca* | | F | USA: NM, Santa Fe Co., Santa Fe National Forest, 15-May-2017 | | | |
| 193 | NVG-16107A03 | *Callophrys niphon* | | F | USA: MN, Lake Co., Lake One, 3-Jun-2016 | WDempwolf | | |
| 194 | NVG-17114E07 | *Callophrys lanoraieensis* | | M | USA: ME, Penobscot Co., Enfield, 19-May-1983 | CSUC | | |
| 195 | NVG-5862 | *Callophrys irus hadros* | | M | USA: TX, Freestone Co., 1 mi NE SH164, 5-Mar-2016 | | | |
| 196 | NVG-5807 | *Callophrys henrici solatus* | | M | USA: TX, Travis Co., Austin, 25-Feb-2016 | | | |
| 197 | NVG-17108A01 | *Callophrys polios obscurus* | | F | USA: WA, Kittitas Co., 13-Apr-1991 | BMUW | | |
| 198 | PAO-441 | *Callophrys mossii windi* | | | USA: CA, Sierra Co., Frazier Falls trail, 25-Jun-2017 | | | |
| 199 | NVG-8626 | *Callophrys augustinus croesioides* | | M | USA: VA, Frederick Co., near Hayfield, 19-Apr-2017 | | | |
| 200 | NVG-17114E06 | *Callophrys fotis* | | M | USA: NM, San Juan Co., Chuska Mts., 1-May-1980 | CSUC | | |
| 201 | NVG-8944 | *Callophrys mcfarlandi* | | F | USA: NM, Otero Co., Lincoln National Forest, 21-May-2017 | | | |
| 202 | NVG-5428 | *Callophrys xami texami* | | M | USA: TX, Cameron Co., 7mi SW Port Isabel, 25-Dec-2015 | | | |
| 203 | PAO-296 | *Callophrys sheridanii sheridanii* | | | USA: CO, Larimer Co., Cherokee Park state wildlife area, 6-May-2017 | | | |
| 204 | NVG-965 | *Callophrys viridis* | | | USA: CA, Monterey Co., Marina Dunes, 6-Mar-2012 | | | |
| 205 | NVG-17107F08 | *Callophrys dumetorum dumetorum* | | M | USA: WA, Mason Co., 29-Apr-1989 | BMUW | | |
| 206 | NVG-6479 | *Callophrys affinis affinis* | | F | USA: CO, Grand Co., 4.3 air mi SE Hot Sulphur Springs, 6-Jul-2016 | | | |
| 207 | NVG-10618 | *Cyanophrys goodsoni* | | M | USA: TX, Hidalgo Co., Santa Ana NWR, 16-Oct-1976 | TAMU | NVG180106-35 | |
| 208 | NVG-17117A08 | *Cyanophrys herodotus* | | F | Mexico: San Luis Potosi, 23-Dec-1972 | TAMU | | |
| 209 | NVG-10617 | *Cyanophrys miserabilis* | | F | USA: TX, Cameron Co., Los Palomas WMA, 29-Jul-1965 | TAMU | NVG180106-34 | |
| 210 | NVG-5161 | *Chlorostrymon simaethis sarita* | | F | USA: TX, Starr Co., Rio Grande City, 15-Nov-2015 | | | |
| 211 | NVG-10605 | *Chlorostrymon telea* | | F | Mexico: Tamaulipas, 8-Jan-1975 | TAMU | NVG180106-22 | |
| 212 | NVG-18033F11 | *Chlorostrymon maesites* | | | USA: FL, Monroe Co., Big Pine Key, 13-Jun-2014 | MWalker | | |
| 213 | NVG-7650 | *Erora quaderna quaderna* | | M | USA: TX, Brewster Co., Big Bend National Park, 22-Jun-1976 | TAMU | NVG170107-81 | |
| 214 | NVG-17076H04 | *Erora laeta* | | M | USA: VT | USNM | | USNMENT00181291 |
| 215 | PAO-92 | *Satyrium behrii behrii* | | | USA: CA, Sierra Co., Tahoe National Forest, 28-Jun-2016 | | | |
| 216 | PAO-26 | *Satyrium tetra* | | | USA: CA, Monterey Co., Chews Ridge, 13-Jun-2016 | | | |
| 217 | NVG-10188 | *Satyrium saepium provo* | | M | USA: CO, Douglas Co., 31-Jul-2017 | | | |

| | | | | | | | | |
|---|---|---|---|---|---|---|---|---|
| 218 | NVG-9638 | *Satyrium titus immaculosus* | | F | USA: UT, Davis Co., Wasatch National Forest, 28-Jul-2017 | | | |
| 219 | PAO-351 | *Satyrium auretorum auretorum* | | | USA: CA, Stanislaus Co., Del Puerto Canyon, 17-May-2017 | | | |
| 220 | NVG-10622 | *Satyrium polingi polingi* | | M | USA: TX, Pecos Co., 20-May-1973 | TAMU | NVG180106-39 | |
| 221 | NVG-17114E04 | *Satyrium ilavia* | | M | USA: AZ, Mohave Co., Hualpai Mountains, 26-Jun-1968 | CSUC | | |
| 222 | NVG-6080 | *Satyrium favonius autolycus* | | F | USA: TX, Wise Co., LBJ National Grassland, 1-May-2016 | | | |
| 223 | NVG-6224 | *Satyrium alcestis alcestis* | | F | USA: TX, Tarrant Co., Benbrook, 28-May-2016 | | | |
| 224 | NVG-4306 | *Satyrium liparops liparops* | | F | USA: IN, Newton Co., 2.5 mi S Lake Village, 2-Aug-2015 | | | |
| 225 | NVG-3486 | *Satyrium kingi* | | F | USA: TX, Tyler Co., Kirby State Forest, 7-Jun-2015 | | | |
| 226 | NVG-18014C09 | *Satyrium caryaevorus* | | | USA: OH, Guemsey Co., Salt Fork State Park, 27-Jun-1982 | USNM | | |
| 227 | NVG-8691 | *Satyrium calanus falacer* | | M | USA: TX, Denton Co., Grapevine Lake, 7-May-2017 | | | |
| 228 | NVG-17114E03 | *Satyrium edwardsii* | | M | USA: NJ, Sussex Co., Newton, 7-Jul-1971 | CSUC | | |
| 229 | NVG-17114E02 | *Satyrium acadica* | | F | USA: OH, Portage Co., Hiram Township, 11-Jul-1998 | CSUC | | |
| 230 | NVG-9560 | *Satyrium californica* | | F | USA: UT, Davis Co., Wasatch National Forest, 28-Jul-2017 | | | |
| 231 | PAO-477 | *Satyrium sylvinus* | | | USA: CA, Inyo Co., Lower Rock Creek Rd., 6-Jul-2017 | | | |
| 232 | NVG-17114E01 | *Satyrium fuliginosa fuliginosa* | | M | USA: CA, Plumas Co., Plumas National Forest, 7-Jul-2012 | CSUC | | |
| 233 | PAO-89 | *Satyrium semiluna* | | | USA: CA, Sierra Co., 1 mi E of Calpine, 28-Jun-2016 | | | |
| 234 | NVG-4159 | *Brephidium pseudofea pseudofea* | | M | USA: TX, Jefferson Co., S of Sabine Pass, 18-Jul-2015 | | | |
| 235 | NVG-8262 | *Brephidium exilis exilis* | | M | USA: UT, Kane Co., west of Big Water, 24-Mar-2017 | | | |
| 236 | NVG-10594 | *Zizula cyna* | | F | USA: TX, Hidalgo Co., Bentsen-Rio Frande Valley State Park, 13-Jul-1972 | TAMU | NVG180106-11 | |
| 237 | NVG-17119F02 | *Zizina otis* | | | Myanmar, 2-9-May-2003 | USNM | | |
| 238 | NVG-17114F04 | *Lampides boeticus* | | M | USA: HI, Kauai, 28-Jul-1982 | CSUC | | |
| 239 | NVG-3444 | *Leptotes cassius cassidula* | | M | USA: TX, Cameron Co., 3 mi SW Sebastian, 31-May-2015 | | | |
| 240 | NVG-8703 | *Leptotes marina* | | F | USA: TX, Randall Co., Palo Duro Canyon State Park, 13-May-2017 | | | |
| 241 | NVG-17066C04 | *Euphilotes leona* | | | USA: OR, Klamath Co., Sand Creek, Hwy 97, 6-Jul-2010 | CSUC | | |
| 242 | NVG-6698 | *Euphilotes speciosa speciosa* | | M | USA: CA, Imperial Co., In Koh Pah Gorge, 3-Apr-2013 | | | |
| 243 | NVG-17114F06 | *Euphilotes battoides battoides* | | M | USA: CA, Mono Co., Minaret Vista, 15-Jun-2007 | CSUC | | |
| 244 | PAO-6 | *Euphilotes glaucon glaucon* | | | USA: CA, Lassen Co., quarry N of Hallelujah Junction on USH395, 10-Jun-2016 | | | |
| 245 | PAO-378 | *Euphilotes bernardino* | | | USA: CA, San Benito Co., Panoche Pass, 19-May-2017 | | | |
| 246 | NVG-17114F09 | *Euphilotes baueri baueri* | | M | USA: CA, Inyo Co., White Mts., Westgard Pass, 29-May-2003 | CSUC | | |
| 247 | NVG-9456 | *Euphilotes ancilla* | | F | USA: WY, Park Co., Yellowstone National Park, 23-Jul-2017 | | | |
| 248 | NVG-17114F11 | *Euphilotes stanfordorum* | | M | USA: CO, Mesa Co., west of Mack, 11-May-2013 | CSUC | | |
| 249 | NVG-17114F07 | *Euphilotes centralis* | | M | USA: CO, El Paso Co., Colorado Springs, 17-Jul-1994 | CSUC | | |
| 250 | NVG-17114F08 | *Euphilotes ellisii ellisii* | | F | USA: UT, Grand Co., Castleton Road, 16-Aug-1996 | CSUC | | |
| 251 | NVG-18014D03 | *Euphilotes mojave mojave* | | | USA: CA, Los Angeles Co., 26-Mar-1964 | USNM | | |
| 252 | NVG-17114F12 | *Euphilotes columbiae* | | M | USA: WA, Kittitas Co., Wenatchee National Forest, 10-Jul-2010 | CSUC | | |
| 253 | PAO-65 | *Euphilotes enoptes* | | | USA: CA, Sierra Co., Tahoe National Forest, 19-Jun-2016 | | | |
| 254 | PAO-192 | *Euphilotes spaldingi spaldingi* | | | USA: UT, Grand Co., E Moab, 7-Aug-2016 | | | |
| 255 | NVG-17114G01 | *Euphilotes pallescens pallescens* | | M | USA: UT, Juab Co., Little Sahara Recreation Area, 23-Jul-2006 | CSUC | | |
| 256 | NVG-7628 | *Euphilotes rita rita* | | M | USA: TX, Brewster Co., Big Bend National Park, 27-Sep-1971 | TAMU | NVG170107-59 | |
| 257 | NVG-17114F05 | *Philotes sonorensis* | | M | USA: CA, Tulare Co., Camp Nelson, 19-Feb-2015 | CSUC | | |
| 258 | NVG-9559 | *Glaucopsyche piasus* | | M | USA: UT, Davis Co., Wasatch National Forest, 28-Jul-2017 | | | |
| 259 | NVG-16106E04 | *Glaucopsyche lygdamus* | | M | USA: MN, Lake Co., Lake One, 3-Jun-2016 | WDempwolf | | |
| 260 | NVG-18014D06 | *Udara blackburni* | | | USA: HI, Maui, 30-Apr-1988 | USNM | | |
| 261 | NVG-5918 | *Celastrina echo echo* | | F | USA: CA, San Mateo Co., SE of Moss Beach, 24-Mar-2016 | | | |
| 262 | NVG-7324 | *Celastrina humulus* | | F | USA: CO, Jefferson Co., Tinytown, 17-Jun-1997 | | | |
| 263 | NVG-6246 | *Celastrina lucia lucia* | | | Canada: Manitoba, Winnipeg, 29-May-1989 | | | |
| 264 | NVG-6887 | *Celastrina ladon* | | M | USA: MD, Cedarville, 23-Mar-2012 | | | |
| 265 | NVG-16106D09 | *Celastrina serotina* | HT | M | USA: RI, Washington Co., Great Swamp, 14-May-1990 | ANSP | | |
| 266 | NVG-6275 | *Celastrina neglectamajor* | | | USA: NC, Haywood Co., Maggie Valley, 12-May-2010 | | | |
| 267 | NVG-16106D08 | *Celastrina idella* | HT | M | USA: NJ, Burlington Co., 2.2 km S Chatsworth, 11-May-1987 | ANSP | | |
| 268 | NVG-4329 | *Celastrina neglecta* | | F | USA: IN, Newton Co., 2.5 mi S Lake Village, 2-Aug-2015 | | | |
| 269 | NVG-6277 | *Celastrina nigra* | | M | USA: WV, Boone Co., Fork Creek PMA, 20-Apr-1990 | | | |
| 270 | NVG-9365 | *Cupido amyntula* | | M | USA: WY, Park Co., Yellowstone National Park, 24-Jul-2017 | | | |
| 271 | NVG-3506 | *Cupido comyntas comyntas* | | F | USA: TX, Hardin Co., 4.4 mi SW Kountze, 7-Jun-2015 | | | |
| 272 | LEP17800 | *Cyclargus ammon* | | M | Cuba: Guantanamo, GTMO Naval Base, site 15 marsh area nr. Ridge trail, 4-Oct-2013 | MGCL | DM 1678 | MGCL 230739 |
| 273 | LEP22359 | *Cyclargus thomasi* | | M | USA: FL, Monroe Co., Bahia Honda State Park, Jul-Aug-2008 | MGCL | | |

| | | | | | | | | |
|---|---|---|---|---|---|---|---|---|
| 274 | NVG-3975 | *Echinargus isola* | | F | USA: TX, Dallas Co., Dallas, Norbuck Park, 7-Jul-2015 | | | |
| 275 | NVG-6771 | *Hemiargus ceraunus astenidas* | | M | USA: TX, Hopkins Co., Cooper Lake State Park, 13-Aug-2016 | | | |
| 276 | NVG-6697 | *Plebulina emigdionis* | | M | USA: CA, Los Angeles Co., Smokey Bear Rd., 16-May-2000 | | | |
| 277 | PAO-161 | *Icaricia shasta pitkinsensis* | | | USA: CO, Clear Creek Co., Loveland Pass, 1-Aug-2016 | | | |
| 278 | NVG-18014D01 | *Icaricia neurona* | | F | USA: CA, Los Angeles Co., 2-Jun-1981 | USNM | | |
| 279 | PAO-78 | *Icaricia acmon* | | | USA: CA, Sierra Co., Tahoe National Forest, 25-Jun-2016 | | | |
| 280 | PAO-5 | *Icaricia lupini lupini* | | | USA: CA, Lassen Co., quarry N of Hallelujah Junction on USH395, 10-Jun-2016 | | | |
| 281 | NVG-17114H03 | *Icaricia cotundra* | | M | USA: CO, Park Co., 10 mi. W Fairplay | CSUC | | |
| 282 | NVG-6351 | *Icaricia icarioides pembina* | | F | USA: CO, Summit Co., 12 air mi NNW Silverthorne, 5-Jul-2016 | | | |
| 283 | PAO-50 | *Icaricia saepiolus* | | | USA: CA, Placer Co., Blackwood Canyon, 17-Jun-2016 | | | |
| 284 | NVG-6409 | *Agriades glandon rustica* | | M | USA: CO, Grand Co., 2 air mi SW Hot Sulphur Springs, 6-Jul-2016 | | | |
| 285 | PAO-94 | *Agriades podarce podarce* | | | USA: CA, Sierra Co., Tahoe National Forest, 28-Jun-2016 | | | |
| 286 | NVG-16107C04 | *Agriades optilete yukona* | | M | Canada: Yukon Territory, Goring Creek, 7-Jul-2016 | WDempwolf | | |
| 287 | NVG-16107B12 | *Plebejus idas alaskensis* | | M | Canada: Yukon Territory, Carcross, 26-Jun-2016 | WDempwolf | | |
| 288 | NVG-17114G12 | *Plebejus fridayi* | | F | USA: CA, Alpine Co., Sonora Pass, 27-Jul-2009 | CSUC | | |
| 289 | PAO-455 | *Plebejus anna* | | | USA: CA, Plumas Co., Frazier Falls Rd., 3-Jul-2017 | | | |
| 290 | NVG-6544 | *Plebejus melissa melissa* | | M | USA: CO, Park Co., 7.9 air mi SW Florissant, 9-Jul-2016 | | | |
| 291 | NVG-7627 | *Plebejus samuelis* | | M | USA: NY, Albany Co., Colonie, 21-Jul-1963 | TAMU | NVG170107-58 | |
| 292 | PAO-E03 | *Polyommatus icarus* | | M | France: rest stop along A41, 15-Aug-2017 | | | |
| 293 | NVG-10595 | *Baeotis zonata zonata* | | F | Mexico: San Luis Potosi, 5-Feb-1980 | TAMU | NVG180106-12 | |
| 294 | NVG-5148 | *Melanis pixe pixe* | | | USA: TX, Hidalgo Co., Penitas RR tracks, 15-Nov-2015 | | | |
| 295 | NVG-7147 | *Calephelis borealis* | | F | USA: MD, Allegany Co., Green Ridge State Forest, 22-Jul-1984 | USNM | NVG161004-74 | |
| 296 | NVG-18042F04 | *Calephelis muticum* | | M | USA: MO, St. Francois Co., St. Francois State Park, 6-Aug-2010 | WDempwolf | | WRD 3900 |
| 297 | NVG-8109 | *Calephelis virginiensis* | | M | USA: FL, Santa Rosa Co., 2.7 air mi E of Munson, 11-Mar-2017 | | | |
| 298 | NVG-5254 | *Calephelis perditalis perditalis* | | F | USA: TX, Cameron Co., E of Brownsville, 27-Nov-2015 | | | |
| 299 | NVG-5951 | *Calephelis nemesis nemesis* | | M | USA: AZ, Pima Co., Santa Rita Mts., 29-Mar-2016 | | | |
| 300 | NVG-5983 | *Calephelis arizonensis* | | M | USA: AZ, Santa Cruz Co., Pajarito Mts., 30-Mar-2016 | | | |
| 301 | NVG-15102B07 | *Calephelis freemani* | HT | | USA: TX, Jeff Davis Co., ca. 12 miles Northwest of Alpine on State Hwy. 118, 5-Jun-1942 | USNM | Slide N109 | |
| 302 | NVG-5887 | *Calephelis rawsoni* | | M | USA: TX, Travis Co., Austin, 5-Mar-2016 | | | |
| 303 | NVG-6683 | *Calephelis wrighti* | | M | USA: CA, San Diego Co., Shelter Valley, 22-Sep-2006 | | | |
| 304 | NVG-10596 | *Lasaia maria anna* | | M | Mexico: Nuevo Leon, 24-Oct-1979 | TAMU | NVG180106-13 | |
| 305 | NVG-4026 | *Lasaia sula peninsularis* | | M | USA: TX, Cameron Co., USH 281, 0.9 mi ESE Encantada-Ranchito El Calaboz, 12-Jul-2015 | | | |
| 306 | NVG-3562 | *Caria ino melicerta* | | F | USA: TX, Cameron Co., 3 mi SW Sebastian, 13-Jun-2015 | | | |
| 307 | NVG-5245 | *Curvie emesia* | | F | USA: TX, Hidalgo Co., Los Ebanos, 25-Nov-2015 | | | |
| 308 | NVG-4514 | *Apodemia walkeri* | | M | USA: TX, Hidalgo Co., Old Rio Rico Rd. 1.5 air mi SE of Relampago, 16-Aug-2015 | | | |
| 309 | NVG-17066E05 | *Apodemia hepburni hepburni* | | | Mexico: Sonora, Rio Sonora Route, 10-Mar-2003 | CSUC | | |
| 310 | NVG-10226 | *Apodemia palmerii arizona* | | | USA: AZ, Pima Co., 21-Sep-2017 | | | |
| 311 | NVG-7635 | *Apodemia multiplaga* | | F | USA: TX, Cameron Co., Brownsville, 24-Oct-1974 | TAMU | NVG170107-66 | |
| 312 | NVG-15105E04 | *Apodemia virgulti virgulti* | NT | M | USA: CA, Los Angeles Co., La Tuna Canyon, 27-Sep-1951 | CAS | | |
| 313 | NVG-17107E02 | *Apodemia mormo mormo* | | M | USA: WA, Kittitas Co., 25-Aug-1990 | BMUW | | |
| 314 | 11-BOA-13383BrockC10 | *Apodemia mejicanus mejicanus* | | F | USA: AZ, Santa Cruz Co., Peck Canyon, 22-Jul-1990 | JimPBrock | | |
| 315 | NVG-700 | *Apodemia duryi* | | M | USA: TX, Brewster Co., Big Bend National Park, 10-Oct-2009 | | | |
| 316 | PAO-145 | *Apodemia nais* | | | USA: CO, Gilpin Co., Golden Gate Canyon State Park, 24-Jul-2016 | | | |
| 317 | NVG-154 | *Apodemia chisosensis* | | F | USA: TX, Brewster Co., Big Bend National Park, 2-May-2004 | | | |
| 318 | NVG-17114H01 | *Apodemia ares* | | M | USA: NM, Hidalgo Co., Pine Canyon, Animas Mountains, 21-Jul-1991 | CSUC | | |
| 319 | NVG-8217 | *Apodemia zela cleis* | | F | USA: AZ, Santa Cruz Co., Coronado N.F., Santa Rita Mts., 25-Mar-2017 | | | |
| 320 | NVG-10598 | *Emesis tenedia* | | F | Mexico: Nuevo Leon, 12-Nov-1980 | TAMU | NVG180106-15 | |
| 321 | NVG-17066E10 | *Emesis phyciodoides* | | M | Mexico: Sonora, Mesa Grande, 10 Mi. NW Yecora on dirt Road, 12-Sep-2004 | CSUC | | |
| 322 | NVG-7278 | *Libytheana motya* | | | Cuba, old | USNM | NVG161007-05 | |
| 323 | NVG-4190 | *Libytheana carinenta bachmanii* | | M | USA: TX, Dallas Co., Dallas, White Rock Creek, 18-Jul-2015 | | | |
| 324 | NVG-9388 | *Coenonympha california ochracea* | | F | USA: WY, Park Co., Yellowstone National Park, 24-Jul-2017 | | | |
| 325 | NVG-9370 | *Coenonympha haydenii* | | M | USA: WY, Park Co., Yellowstone National Park, 24-Jul-2017 | | | |
| 326 | NVG-17115C09 | *Lethe eurydice fumosa* | | M | USA: CO, Larimer Co., LaPorte, 28-Jun-1985 | CSUC | | |
| 327 | NVG-4359 | *Lethe appalachia leeuwi* | | F | USA: IN, Lake Co., Gary, 2-Aug-2015 | | | |
| 328 | NVG-7687 | *Lethe anthedon anthedon* | | M | USA: TX, Wise Co., 9 mi N. Decatur, 31-Aug-1997 | TAMU | NVG170108-18 | |
| 329 | NVG-4416 | *Lethe portlandia missarkae* | | F | USA: TX, Lamar Co., FM1499 at Craddock Creek, 8-Aug-2015 | | | |

| | | | | | | | |
|---|---|---|---|---|---|---|---|
| 330 | NVG-3317 | *Lethe creola* | | M | USA: TX, San Jacinto Co., Sam Houston NF, Big Creek Scenic Area, 12-Apr-2015 | | |
| 331 | NVG-17068E09 | *Paramacera xicaque allyni* | | M | USA: AZ, Cochise Co., Chiricahua Mts., 2-Jul-2007 | CSUC | |
| 332 | NVG-17115C10 | *Cyllopsis pyracmon* | | M | USA: AZ, Santa Cruz Co., Santa Rita Mountains, 24-Jun-1968 | CSUC | |
| 333 | NVG-9714 | *Cyllopsis pertepida* | | M | USA: TX, Jeff Davis Co., Davis Mnts., 4-Aug-2017 | | |
| 334 | NVG-9181 | *Cyllopsis gemma gemma* | | F | USA: AR, Scott Co., Ouachita National Forest, 6-Jul-2017 | | |
| 335 | NVG-8382 | *Cissia rubricata rubricata* | | M | USA: TX, Blanco Co., Middle Creek Rd. nr. USH290, 31-Mar-2017 | | |
| 336 | NVG-3689 | *Megisto cymela cymela* | | F | USA: TX, Harrison Co., Caddo Lake State Park, 20-Jun-2015 | | |
| 337 | NVG-3310 | *Hermeuptychia sosybius* | | M | USA: TX, Sabine Co., Sabine NF, 12-Apr-2015 | | |
| 338 | NVG-1747 | *Hermeuptychia hermybius* | PT | | USA: TX, Starr Co., Salineno at Rio Grande, 23-Oct-2013 | | |
| 339 | NVG-1563 | *Hermeuptychia intricata* | PT | | USA: TX, Fort Bend Co., Brazos Bend SP, 17-Aug-2013 | | |
| 340 | NVG-3269 | *Neonympha mitchellii mitchellii* | | M | USA: MI, Cass Co., 1 mi NE Wakelee, 3-Jul-1971 | TAMU | NVG15011-85 |
| 341 | NVG-2121 | *Neonympha areolatus* | | | USA: TX, Hardin Co., Kountze, 2009 | | |
| 342 | NVG-10223 | *Gyrocheilus patrobas tritonia* | | F | USA: AZ, Santa Cruz Co., Coronado N.F., Santa Rita Mts., 21-Sep-2017 | | |
| 343 | NVG-4506 | *Cercyonis pegala texana* | | F | USA: TX, Wise Co., LBJ National Grassland, 9-Aug-2015 | | |
| 344 | NVG-9708 | *Cercyonis meadii* | | M | USA: TX, Jeff Davis Co., Davis Mnts., 4-Aug-2017 | | |
| 345 | NVG-7259 | *Cercyonis sthenele sthenele* | ST | F | USA: CA, San Fransico Co., San Fransico, very old | USNM | NVG161005-86 |
| 346 | PAO-435 | *Cercyonis incognita* | | | USA: CA, Mendocino Co., 2 mi W Mendocino Pass, 23-Jun-2017 | | |
| 347 | PAO-112 | *Cercyonis oetus oetus* | | | USA: NV, Elko Co., Pequop Summit, 1-Jul-2016 | | |
| 348 | NVG-6549 | *Erebia magdalena magdalena* | | M | USA: CO, Park Co., 9.5 air mi W Fairplay, 10-Jul-2016 | | |
| 349 | NVG-16106E11 | *Erebia mackinleyensis* | | M | Canada: Yukon Territory, Dempster Hwy km 465, 3-Jul-2016 | WDempwolf | |
| 350 | NVG-16106F02 | *Erebia fasciata* | | M | Canada: Yukon Territory, Montana Mountain, 24-Jun-2016 | WDempwolf | |
| 351 | NVG-7999 | *Erebia discoidalis discoidalis* | | M | USA: WI, Oneida Co., Bog 0.8 mi S of Villas Co. line on Hwy17, 17-May-1988 | USNM | NVG170207-84 |
| 352 | NVG-16106E05 | *Erebia rossii* | | M | Canada: Yukon Territory, Dempster Hwy km 155, 5-Jul-2016 | WDempwolf | |
| 353 | NVG-16106E08 | *Erebia disa* | | M | Canada: Yukon Territory, Dempster Hwy km 464, 4-Jul-2016 | WDempwolf | |
| 354 | NVG-16106F06 | *Erebia mancinus* | | M | Canada: Yukon Territory, Dempster Hwy km 151, 5-Jul-2016 | WDempwolf | |
| 355 | NVG-16106G06 | *Erebia lafontainei* | | M | Canada: Yukon Territory, Montana Mountain, 24-Jun-2016 | WDempwolf | |
| 356 | NVG-16106G03 | *Erebia youngi youngi* | | M | Canada: Yukon Territory, Dempster Hwy km 465, 2-Jul-2016 | WDempwolf | |
| 357 | NVG-16106G09 | *Erebia occulta occulta* | | M | Canada: Yukon Territory, Dempster Hwy km 464, 4-Jul-2016 | WDempwolf | |
| 358 | NVG-16106G01 | *Erebia pawloskii alaskensis* | | M | Canada: Yukon Territory, Fish Lake Road, 28-Jun-2016 | WDempwolf | |
| 359 | NVG-6457 | *Erebia epipsodea brucei* | | F | USA: CO, Grand Co., 4 air mi SSE Hot Sulphur Springs, 6-Jul-2016 | | |
| 360 | NVG-8001 | *Erebia vidleri* | | | USA: WA, Whatcom Co., Skyline Ridge, 30-Jul-1980 | USNM | NVG170207-86 |
| 361 | NVG-9376 | *Erebia callias* | | F | USA: WY, Park Co., Yellowstone National Park, 24-Jul-2017 | | |
| 362 | NVG-17115C11 | *Oeneis ridingsii ridingsii* | | M | USA: CO, Larimer Co., Horsetooth Mountain Park, 22-Jun-1991 | CSUC | |
| 363 | NVG-16106H04 | *Oeneis jutta* | | M | Canada: Yukon Territory, Dempster Hwy km 131 (west), 6-Jul-2016 | WDempwolf | |
| 364 | NVG-16106H06 | *Oeneis melissa* | | F | Canada: Yukon Territory, Dempster Hwy km 131 (west), 5-Jul-2016 | WDempwolf | |
| 365 | NVG-16106G12 | *Oeneis polixenes* | | M | Canada: Yukon Territory, Montana Mountain, 27-Jun-2016 | WDempwolf | |
| 366 | NVG-17068D10 | *Oeneis philipi* | PT | M | USA: AK, Ballaine Rd., 14-Jun-1985 | CSUC | |
| 367 | NVG-8003 | *Oeneis alpina excubitor* | | | Canada: Yukon, 4 km E My. Chambers, 16-Jun-1981 | USNM | NVG170207-88 |
| 368 | NVG-16106H08 | *Oeneis bore* | | M | Canada: Yukon Territory, Montana Mountain, 27-Jun-2016 | WDempwolf | |
| 369 | NVG-11482 | *Oeneis alberta daura* | | F | USA: AZ, Apache Co., 25-May-2018 | | |
| 370 | NVG-5203 | *Oeneis tanana* | | F | USA: AK, Alaska Hwy., mi. 1410, 12 mi SE Delta Jct., 15-Jun-2001 | MGCL | |
| 371 | NVG-6464 | *Oeneis chryxus chryxus* | | F | USA: CO, Grand Co., 4.3 air mi SE Hot Sulphur Springs, 6-Jul-2016 | | |
| 372 | PAO-73 | *Oeneis nevadensis* | | | USA: CA, Plumas Co., Plumas National Forest, 23-Jun-2016 | | |
| 373 | NVG-16107A02 | *Oeneis macounii* | | M | USA: MN, Lake of the Woods Co., Norris Camp, 4-Jun-2016 | WDempwolf | |
| 374 | NVG-6583 | *Oeneis uhleri uhleri* | | M | USA: CO, Park Co., 3 air mi SW Fairplay, 10-Jul-2016 | | |
| 375 | NVG-17117D08 | *Morpho polyphemus polyphemus* | | M | Mexico: Nayarit, 11-Oct-1975 | TAMU | |
| 376 | NVG-19065A09 | *Prepona laertes octavia* | | | Belize, 9-16-Jun-2016 | UCDC | |
| 377 | NVG-17117F10 | *Archaeoprepona demophon centralis* | | M | Mexico: Tamaulipas, 24-Feb-1974 | TAMU | |
| 378 | NVG-17118A09 | *Memphis forreri* | | M | Mexico: Veracruz, 19-Jul-1980 | TAMU | |
| 379 | NVG-17118A08 | *Memphis pithyusa pithyusa* | | M | Mexico: Tamaulipas, 18-Jul-1994 | TAMU | |
| 380 | NVG-17119F12 | *Memphis echemus* | | | Cuba, 7-Oct-1962 | USNM | |
| 381 | NVG-17118A11 | *Fountainea glycerium glycerium* | | | Mexico: Tamaulipas, 18-Jul-1994 | TAMU | |
| 382 | NVG-3982 | *Anaea andria* | | F | USA: TX, Marion Co., FM805 near the lake, 9-Jul-2015 | | |
| 383 | NVG-3787 | *Anaea aidea* | | F | USA: TX, Starr Co., Roma International Bridge, 27-Jun-2015 | | |
| 384 | NVG-17115F10 | *Anaea troglodyta floridalis* | | M | USA: FL, Monroe Co., 7-Aug-1967 | CSUC | |
| 385 | NVG-17117H07 | *Doxocopa pavon theodora* | | F | Mexico: Tamaulipas, 2-Mar-1974 | TAMU | |

| | | | | | | | |
|---|---|---|---|---|---|---|---|
| 386 | NVG-3838 | *Doxocopa laure laure* | | M | USA: TX, Starr Co., Roma International Bridge, 28-Jun-2015 | | |
| 387 | NVG-4189 | *Asterocampa celtis celtis* | | M | USA: TX, Dallas Co., Coppell, 18-Jul-2015 | | |
| 388 | NVG-4022 | *Asterocampa leilia* | | F | USA: TX, Cameron Co., 3 mi SW Sebastian, 12-Jul-2015 | | |
| 389 | NVG-4068 | *Asterocampa clyton clyton* | | F | USA: TX, Dallas Co., Dallas, 14-Jul-2015 | | |
| 390 | NVG-17112H07 | *Asterocampa idyja argus* | | | Mexico, 8-Aug-1974 | TMMC | |
| 391 | NVG-17118B05 | *Epiphile adrasta adrasta* | | | Mexico: Tamaulipas, 26-27-Aug-1985 | TAMU | |
| 392 | NVG-17118B03 | *Temenis laothoe hondurensis* | | | Mexico: Tamaulipas, 18-Jul-1994 | TAMU | |
| 393 | NVG-10640 | *Eunica monima* | | F | USA: TX, Cameron Co., Brownsville, 27-Jun-1969 | TAMU | NVG180106-57 |
| 394 | NVG-17117E10 | *Eunica tatila tatila* | | M | Mexico: San Luis Potosi, 18-Feb-1976 | TAMU | |
| 395 | NVG-17118B11 | *Myscelia cyananthe cyananthe* | | | Mexico: Oaxaca, 13-Jul-1987 | TAMU | |
| 396 | NVG-4049 | *Myscelia ethusa ethusa* | | M | USA: TX, Cameron Co., USH 281, 0.9 mi ESE Encantada-Ranchito El Calaboz, 12-Jul-2015 | | |
| 397 | NVG-17118B10 | *Hamadryas atlantis lelaps* | | | Mexico: Oaxaca, 11-17-Jul-1981 | TAMU | |
| 398 | NVG-17117G06 | *Hamadryas februa ferentina* | | F | Mexico: Tamaulipas, 19-Nov-1980 | TAMU | |
| 399 | NVG-17117G07 | *Hamadryas glauconome glauconome* | | M | Mexico: Tamaulipas, 24-Feb-1974 | TAMU | |
| 400 | NVG-17119D09 | *Hamadryas iphthime joannae* | | | Mexico: Yucatan, Sep-1978 | USNM | |
| 401 | NVG-17117G09 | *Hamadryas feronia farinulenta* | | M | USA: TX, Hidalgo Co., 15-Jul-1975 | TAMU | |
| 402 | NVG-17117G11 | *Hamadryas guatemalena marmarice* | | M | Mexico: San Luis Potosi, 20-Feb-1980 | TAMU | |
| 403 | NVG-17113A09 | *Hamadryas amphinome mexicana* | | | Mexico, Aug-1976 | TMMC | |
| 404 | NVG-17117F03 | *Biblis hyperia* | | M | USA: TX, Hidalgo Co., 19-Oct-1973 | TAMU | |
| 405 | NVG-3631 | *Mestra amymone* | | F | USA: TX, Cameron Co., 3 mi SW Sebastian, 13-Jun-2015 | | |
| 406 | NVG-3563 | *Dynamine dyonis* | | M | USA: TX, Cameron Co., 3 mi SW Sebastian, 13-Jun-2015 | | |
| 407 | NVG-10642 | *Dynamine postverta mexicana* | | M | Mexico: Tamaulipas, 14-Nov-1974 | TAMU | NVG180106-59 |
| 408 | NVG-17117F06 | *Marpesia chiron* | | F | USA: TX, Bexar Co., 28-Oct-1983 | TAMU | |
| 409 | NVG-17118B01 | *Marpesia zerynthia dentiger* | | | Mexico: Oaxaca, 12-Jul-1987 | TAMU | |
| 410 | NVG-4890 | *Marpesia petreus* | | M | USA: FL, Miami-Dade Co., Homestead, 4-Oct-2015 | | |
| 411 | NVG-17119D12 | *Marpesia eleuchea* | | | Cuba, 4-9-May-2010 | USNM | |
| 412 | NVG-17118B07 | *Historis acheronta acheronta* | | M | USA: TX, Presidio Co., Shafter, 13-Aug-1969 | TAMU | |
| 413 | NVG-17118B08 | *Historis odius odius* | | | Mexico: Veracruz, 19-Jul-1980 | TAMU | |
| 414 | NVG-17118B06 | *Smyrna karwinskii* | | | Mexico: Tamaulipas, 26-30-Jul-1993 | TAMU | |
| 415 | NVG-17117H09 | *Smyrna blomfildia* | | F | USA: TX, Nueces Co., 28-Oct-1984 | TAMU | |
| 416 | NVG-17118C08 | *Hypanartia lethe* | | | Mexico: Oaxaca, 12-Jul-1987 | TAMU | |
| 417 | PAO-E20 | *Inachis io* | | | Germany: Bavaria, near Grasswang, 12-Aug-2017 | | |
| 418 | NVG-9460 | *Aglais milberti subpallida* | | M | USA: WY, Park Co., Yellowstone National Park, 23-Jul-2017 | | |
| 419 | NVG-6030 | *Polygonia interrogationis* | | F | USA: TX, Dallas Co., Dallas, Norbuck Park, 7-Apr-2016 | | |
| 420 | NVG-17117D09 | *Polygonia comma* | | F | USA: TX, Polk Co., 25-Oct-1971 | TAMU | |
| 421 | PAO-42 | *Polygonia satyrus satyrus* | | | USA: CA, Monterey Co., Old Coast Road off SH1, 14-Jun-2016 | | |
| 422 | NVG-17115B04 | *Polygonia progne* | | M | USA: SD, Roberts Co., 7-Jul-1981 | CSUC | |
| 423 | NVG-17115B05 | *Polygonia oreas silenus* | | M | USA: OR, Yamhill Co., 21-Jul-1983 | CSUC | |
| 424 | PAO-246 | *Polygonia gracilis zephyrus* | | | USA: CO, Larimer Co., 2 mi N Virginia Dale, 3-Sep-2016 | | |
| 425 | NVG-17115B06 | *Polygonia haroldii* | | M | Mexico: Hidalgo, 18 miles northeast of Zimapan, 4-Aug-1981 | CSUC | |
| 426 | PAO-270 | *Polygonia faunus hylas* | | | USA: CO, Larimer Co., Sky Ranch Road off Pingree Park Road, 10-Sep-2016 | | |
| 427 | NVG-17115B03 | *Nymphalis l-album watsoni* | | M | USA: MT, Lake Co., Flathead Lake, 26-May-1976 | CSUC | |
| 428 | NVG-9635 | *Nymphalis californica* | | | USA: UT, Davis Co., Wasatch National Forest, 28-Jul-2017 | | |
| 429 | NVG-8980 | *Nymphalis antiopa* | | F | USA: NM, Otero Co., Lincoln National Forest, 21-May-2017 | | |
| 430 | NVG-9614 | *Vanessa annabella* | | M | USA: UT, Davis Co., Wasatch National Forest, 28-Jul-2017 | | |
| 431 | NVG-8641 | *Vanessa cardui* | | F | USA: TX, Dallas Co., Dallas, 23-Apr-2017 | | |
| 432 | NVG-3866 | *Vanessa virginiensis* | | F | USA: TX, Dallas Co., Dallas, White Rock Creek, 3-Jul-2015 | | |
| 433 | NVG-17115B02 | *Vanessa tameamea* | | M | USA: HI, Hawaii, 5-Mar-1974 | CSUC | |
| 434 | NVG-9826 | *Vanessa atalanta rubria* | | F | USA: TX, Dallas Co., Dallas, White Rock Lake Park, 12-Aug-2017 | | |
| 435 | NVG-5356 | *Hypolimnas misippus* | | M | USA: FL, Miami-Dade Co., 6 mi SW of Florida City, 20-Dec-2015 | | |
| 436 | NVG-3935 | *Junonia coenia* | | F | USA: AR, Scott Co., Ouachita National Forest, 4-Jul-2015 | | |
| 437 | NVG-18098H10 | *Junonia grisea* | | F | USA: CA, Los Angeles Co., Pico Rivera, 10-Sep-2016 | | |
| 438 | NVG-5950 | *Junonia nigrosuffusa* | | M | USA: AZ, Pima Co., Santa Rita Mts., 29-Mar-2016 | | |
| 439 | NVG-8165 | *Junonia zonalis zonalis* | | M | USA: FL, Miami-Dade Co., 6 mi SW of Florida City, 19-Mar-2017 | | |
| 440 | NVG-4818 | *Junonia neildi* | | M | USA: FL, Collier Co., Everglades City Airport, 2-Oct-2015 | | |
| 441 | NVG-6649 | *Junonia litoralis* | | F | French Guiana: Macouria, 17-Jun-2016 | | |

| | | | | | | | | |
|---|---|---|---|---|---|---|---|---|
| 442 | NVG-17119E08 | *Anartia chrysopelea* | | | Cuba: Habana, 5-Sep-1959 | USNM | | |
| 443 | NVG-3615 | *Anartia jatrophae luteipicta* | | M | USA: TX, Starr Co., Roma International Bridge, 14-Jun-2015 | | | |
| 444 | NVG-17117D12 | *Anartia fatima fatima* | | M | USA: TX, Hidalgo Co., 22-May-1972 | TAMU | | |
| 445 | NVG-3989 | *Siproeta stelenes biplagiata* | | F | USA: TX, Starr Co., Roma International Bridge, 10-Jul-2015 | | | |
| 446 | NVG-17112F12 | *Siproeta epaphus epaphus* | | | Ecuador, 30-May-1994 | TMMC | | |
| 447 | NVG-10639 | *Tegosa luka* | | M | Mexico: Tamaulipas, 12-Jan-1974 | TAMU | NVG180106-56 | |
| 448 | NVG-4378 | *Anthanassa texana texana* | | F | USA: TX, Dallas Co., Dallas, 7-Aug-2015 | | | |
| 449 | NVG-10631 | *Anthanassa ptolyca ptolyca* | | M | Mexico: Tamaulipas, 29-Nov-1986 | TAMU | NVG180106-48 | |
| 450 | NVG-17118C05 | *Anthanassa argentea* | | M | Mexico: Tamaulipas, 4-Jul-1986 | TAMU | | |
| 451 | NVG-10634 | *Anthanassa tulcis* | | F | USA: TX, Hidalgo Co., Santa Ana NWR, 12-Dec-1970 | TAMU | NVG180106-51 | |
| 452 | NVG-10629 | *Anthanassa frisia frisia* | | F | USA: FL, Monroe Co., Key Largo, 13-Jul-1983 | TAMU | NVG180106-46 | |
| 453 | NVG-3395 | *Phyciodes tharos tharos* | | F | USA: TX, Hidalgo Co., Old Rio Rico Rd. 1.5 air mi SE of Relampago, 30-May-2015 | | | |
| 454 | NVG-18022C03 | *Phyciodes cocyta cocyta* | NT | M | Canada: Nov Scotia, Cape Breton, 23-Jun-1983 | AMNH | | |
| 455 | NVG-15098D04 | *Phyciodes batesii batesii* | ST | M | USA: Virginia, Winchester, prior to 1866 | FMNH | | |
| 456 | PAO-52 | *Phyciodes pulchella montana* | | | USA: CA, Placer Co., Blackwood Canyon, 17-Jun-2016 | | | |
| 457 | NVG-17115E12 | *Phyciodes mylitta arizonensis* | | F | USA: NM, Colfax Co., Raton Pass, 5-Oct-1997 | CSUC | | |
| 458 | NVG-17115F01 | *Phyciodes pallida pallida* | | F | USA: CO, Larimer Co., Fort Collins, 21-Jun-1988 | CSUC | | |
| 459 | NVG-17115E10 | *Phyciodes orseis orseis* | | M | USA: CA, Siskiyou, Shasta City area, no date | CSUC | | |
| 460 | NVG-3405 | *Phyciodes phaon phaon* | | M | USA: TX, Hidalgo Co., Old Rio Rico Rd. 1.5 air mi SE of Relampago, 30-May-2015 | | | |
| 461 | NVG-17115C03 | *Phyciodes pallescens* | | M | Mexico: Sinaloa, 28-Apr-1998 | CSUC | | |
| 462 | NVG-10637 | *Phyciodes picta picta* | | F | USA: TX, Randall Co., 28-Jun-1975 | TAMU | NVG180106-54 | |
| 463 | NVG-3386 | *Phyciodes graphica vesta* | | M | USA: TX, Hidalgo Co., Edinburg, 24-May-2015 | | | |
| 464 | NVG-3468 | *Texola elada ulrica* | | F | USA: TX, Duval Co., Parilla Creek, North Fork at SH16, 31-May-2015 | | | |
| 465 | NVG-5971 | *Texola perse* | | M | USA: AZ, Santa Cruz Co., Pajarito Mtns., 30-Mar-2016 | | | |
| 466 | NVG-5963 | *Dymasia dymas chara* | | M | USA: AZ, Pima Co., Santa Rita Mts., 29-Mar-2016 | | | |
| 467 | NVG-10626 | *Microtia elva elva* | | M | USA: TX, Cameron Co., Brownsville, 20-Jul-1973 | TAMU | NVG180106-43 | |
| 468 | NVG-17117C11 | *Chlosyne melitaeoides* | | M | USA: TX, Starr Co., 22-Oct-1974 | TAMU | | |
| 469 | NVG-17119E12 | *Chlosyne eumeda* | | | Mexico: Nayarit, 5-Sep-1932 | USNM | | |
| 470 | NVG-17117C05 | *Chlosyne endeis pardelina* | | F | USA: TX, Duval Co., 13-Sep-1980 | TAMU | | |
| 471 | NVG-3462 | *Chlosyne definita* | | F | USA: TX, Duval Co., Parilla Creek, South Fork at SH16, 31-May-2015 | | | |
| 472 | NVG-5108 | *Chlosyne janais janais* | | | USA: TX, Starr Co., Rio Grande City, 14-Nov-2015 | | | |
| 473 | NVG-16107E02 | *Chlosyne rosita browni* | | M | USA: TX, Hidalgo Co., La Lomita historical site, 7-Aug-2013 | JMcDermott | | |
| 474 | NVG-5209 | *Chlosyne theona* | | M | USA: TX, Starr Co., Roma Creek, 25-Nov-2015 | | | |
| 475 | NVG-17117D01 | *Chlosyne cyneas* | | F | Mexico: Tlaxcala, 31-Mar-1977 | TAMU | | |
| 476 | NVG-17117D04 | *Chlosyne fulvia fulvia* | | M | USA: TX, Terrell Co., 3-Oct-1988 | TAMU | | |
| 477 | PAO-352 | *Chlosyne leanira leanira* | | | USA: CA, Stanislaus Co., Del Puerto Canyon, 17-May-2017 | | | |
| 478 | NVG-17115B11 | *Chlosyne californica* | | M | USA: AZ, Yavapai Co., 8-Sep-2007 | CSUC | | |
| 479 | NVG-3753 | *Chlosyne lacinia adjutrix* | | F | USA: TX, Starr Co., Roma International Bridge, 26-Jun-2015 | | | |
| 480 | NVG-8406 | *Chlosyne gorgone carlota* | | F | USA: TX, Wise Co., LBJ National Grassland, 4-Apr-2017 | | | |
| 481 | NVG-4444 | *Chlosyne nycteis nycteis* | | F | USA: TX, Wise Co., LBJ National Grassland, 9-Aug-2015 | | | |
| 482 | NVG-17115B12 | *Chlosyne harrisii* | | M | USA: ME, Cumberland Co., 7-Jul-1982 | CSUC | | |
| 483 | PAO-79 | *Chlosyne hoffmanni* | | | USA: CA, Sierra Co., Tahoe National Forest, 26-Jun-2016 | | | |
| 484 | NVG-8314 | *Chlosyne acastus sabina* | | F | USA: AZ, Pinal Co., Coronado N.F., Catalina Mts., S of Oracle, 26-Mar-2017 | | | |
| 485 | PAO-375 | *Chlosyne gabbii* | | | USA: CA, San Benito Co., BLM land, 19-May-2017 | | | |
| 486 | PAO-335 | *Chlosyne palla palla* | | | USA: CA, Stanislaus Co., Del Puerto Canyon, 15-May-2017 | | | |
| 487 | NVG-17115C01 | *Chlosyne whitneyi* | | M | USA: CA, Mono Co., Sonora Pass, 25-Aug-1974 | CSUC | | |
| 488 | NVG-17115C02 | *Chlosyne damoetas* | | M | USA: CO, Gilpin Co., Rollins Pass, 23-Jul-1997 | CSUC | | |
| 489 | NVG-9586 | *Poladryas arachne* | | F | USA: AZ, Coconino Co., Coconino National Forest, 11.3 air mi N of Flagstaff, 29-Jul-2017 | | | |
| 490 | NVG-7659 | *Poladryas minuta minuta* | | M | USA: TX, Terrell Co., 9 mi west of Dryden, 8-Oct-1987 | TAMU | NVG170107-90 | |
| 491 | NVG-17115B07 | *Euphydryas gillettii* | | M | USA: MT, Cascade Co., Little Belt Mountains, 16-Jul-1996 | CSUC | | |
| 492 | NVG-17115B08 | *Euphydryas phaeton* | | M | USA: OH, Portage Co., Pioneer Trail Road, 11-Jul-1983 | CSUC | | |
| 493 | PAO-31 | *Euphydryas chalcedona chalcedona* | | | USA: CA, Monterey Co., Chews Ridge, 13-Jun-2016 | | | |
| 494 | NVG-17115B09 | *Euphydryas colon* | | M | USA: OR, Josephine Co., Little Grayback Pass, 9-Jul-2008 | CSUC | | CSU-CPG-LEP-0520 |
| 495 | NVG-9446 | *Euphydryas anicia windi* | | M | USA: WY, Park Co., Yellowstone National Park, 23-Jul-2017 | | | |
| 496 | PAO-334 | *Euphydryas editha luestherae* | | | USA: CA, Stanislaus Co., Del Puerto Canyon, 15-May-2017 | | | |
| 497 | NVG-17114H07 | *Boloria eunomia caelestis* | | F | USA: WY, Sheridan Co., Big Horn Mts., 18-Jul-1999 | CSUC | | |

| | | | | | | | | |
|---|---|---|---|---|---|---|---|---|
| 498 | NVG-17115A07 | *Boloria alaskensis halli* | | F | USA: WY, Sublette Co., N. Wind River Mts., 4-Aug-1992 | CSUC | | |
| 499 | NVG-17115A02 | *Boloria natazhati natazhati* | | M | Canada: NWT, Bernard Harbour, 1-18-July-1988 | CSUC | | |
| 500 | NVG-18037H01 | *Boloria freija* | | M | Canada: YT, Nickel Creek, 23-Jun-1992 | CSUC | | |
| 501 | NVG-6356 | *Boloria chariclea helena* | | F | USA: CO, Grand Co., 13.5 air mi N Silverthorne, 5-Jul-2016 | | | |
| 502 | NVG-17115A09 | *Boloria improba harryi* | | M | USA: WY, Sublette Co., Wind River Range, 12-Jul-1990 | CSUC | | |
| 503 | NVG-17114H09 | *Boloria frigga saga* | | M | USA: WY, Albany Co., Snowy Range, 13-Jun-1996 | CSUC | | |
| 504 | NVG-6098 | *Boloria bellona bellona* | | F | USA: VA, Rockingham Co., FS85, 7-May-2016 | | | |
| 505 | PAO-77 | *Boloria epithore epithore* | | | USA: CA, Sierra Co., Tahoe National Forest, 25-Jun-2016 | | | |
| 506 | NVG-9479 | *Boloria kriemhild* | | M | USA: WY, Park Co., Yellowstone National Park, 23-Jul-2017 | | | |
| 507 | NVG-17114H08 | *Boloria selene tollandensis* | | F | USA: ID, Bear River Co., NNE Geneva, 16-Jun-2003 | CSUC | | |
| 508 | NVG-17114H11 | *Boloria polaris* | | M | Canada: YT, 10-Jun-1987 | CSUC | | |
| 509 | NVG-17119E11 | *Boloria alberta* | | | Canada: Alberta, Plateau Mountain, 22-Jul-1967 | USNM | | |
| 510 | NVG-17115A01 | *Boloria astarte astarte* | | F | Canada: Alberta, Plateau Mountain, 21-Jul-2003 | CSUC | | |
| 511 | NVG-17115A04 | *Speyeria idalia occidentalis* | | M | USA: CO, Kit Carson Co., 4 miles east of Flagler, 7-Jul-1987 | CSUC | | |
| 512 | NVG-9190 | *Speyeria diana* | | M | USA: AR, Polk Co., Ouachita National Forest, 6-Jul-2017 | | | |
| 513 | NVG-18062B01 | *Speyeria nokomis nitocris* | | F | USA: NM, Catron Co., Mogollon Mts., 12-Sep-1978 | USNM | | |
| 514 | NVG-9191 | *Speyeria cybele cybele* | | F | USA: AR, Scott Co., Ouachita National Forest, 6-Jul-2017 | | | |
| 515 | PAO-220 | *Speyeria aphrodite ethne* | | | USA: CO, Larimer Co., 2 miles NNE Virginia Dale, 14-Aug-2016 | | | |
| 516 | PAO-143 | *Speyeria coronis halcyone* | | | USA: WY, Goshen Co., Lone Tree Canyon, 16-Jul-2016 | | | |
| 517 | NVG-18028A02 | *Speyeria carolae* | | M | USA: NV, Clark Co., 15-Jun-1967 | USNM | | |
| 518 | NVG-9626 | *Speyeria zerene picta* | | M | USA: OR, Umatilla Co., Umatilla National Forest, 27-Jul-2017 | | | |
| 519 | NVG-9516 | *Speyeria callippe* | | M | USA: UT, Davis Co., Wasatch National Forest, 28-Jul-2017 | | | |
| 520 | NVG-9532 | *Speyeria egleis utahensis* | | F | USA: UT, Davis Co., Wasatch National Forest, 28-Jul-2017 | | | |
| 521 | NVG-19051H09 | *Speyeria adiaste adiaste* | | M | USA: CA, Santa Cruz Co., Saratoga Gap, 9-Jul-1959 | CSUC | | |
| 522 | NVG-9433 | *Speyeria hydaspe rhodope* | | | USA: WY, Park Co., Yellowstone National Park, 23-Jul-2017 | | | |
| 523 | NVG-19065C03 | *Speyeria atlantis atlantis* | | | Canada: Ontario, 20-Aug-2013 | UCDC | | |
| 524 | NVG-11628 | *Speyeria hesperis nausicaa* | | M | USA: AZ, Greenlee Co., 27-May-2018 | | | |
| 525 | PAO-218 | *Speyeria edwardsii* | | | USA: CO, Larimer Co., 2 miles NNE Virginia Dale, 14-Aug-2016 | | | |
| 526 | NVG-9380 | *Speyeria mormonia eurynome* | | | USA: WY, Park Co., Yellowstone National Park, 24-Jul-2017 | | | |
| 527 | NVG-3706 | *Euptoieta claudia* | | F | USA: TX, Dallas Co., Dallas, 21-Jun-2015 | | | |
| 528 | NVG-3374 | *Euptoieta hegesia meridiania* | | F | USA: TX, Hidalgo Co., Penitas RR tracks, 23-May-2015 | | | |
| 529 | NVG-3451 | *Dione vanillae incarnata* | | M | USA: TX, Hidalgo Co., Old Rio Rico Rd. 1.5 air mi SE of Relampago, 31-May-2015 | | | |
| 530 | NVG-14114D10 | *Dione moneta* | | M | USA: TX, Hidalgo Co., Mission, ecl. on 13-Jan-2014 | TLS | | |
| 531 | NVG-17117B01 | *Dione juno huascuma* | | M | Mexico: San Luis Potosi, 28-Nov-1978 | TAMU | | |
| 532 | NVG-17117B05 | *Heliconius erato petiverana* | | M | Mexico: San Luis Potosi, 11-Feb-1976 | TAMU | | |
| 533 | NVG-4695 | *Heliconius charithonia tuckeri* | | | USA: FL, Levy Co., Yankeetown, 26-Sep-2015 | | | |
| 534 | NVG-17117B03 | *Eueides isabella eva* | | F | USA: TX, San Patricio Co., 20-Apr-1968 | TAMU | | |
| 535 | NVG-17117H03 | *Dryadula phaetusa* | | M | USA: TX, Dallas Co., Garland, 19-Jul-1981 | TAMU | | |
| 536 | NVG-4900 | *Dryas iulia largo* | | | USA: FL, Miami-Dade Co., Homestead, west of Turkey Point, 4-Oct-2015 | | | |
| 537 | PAO-34 | *Adelpha californica* | | | USA: CA, Monterey Co., Chews Ridge, 13-Jun-2016 | | | |
| 538 | NVG-8977 | *Adelpha eulalia* | | | USA: NM, Otero Co., Lincoln National Forest, 21-May-2017 | | | |
| 539 | NVG-17118B12 | *Adelpha basiloides* | | | Mexico: Campeche, 26-27-Jul-1980 | TAMU | | |
| 540 | NVG-10006 | *Adelpha fessonia fessonia* | | F | USA: TX, Cameron Co., Resaca de la Palma State Park, 20-Aug-2017 | | | |
| 541 | NVG-15113D11 | *Limenitis arthemis arthemis* | | F | USA: WI, Vilas Co., Lac du Flambeau, 11-Jul-2001 | FMNH | | |
| 542 | NVG-15113D09 | *Limenitis weidemeyerii* | | M | USA: AZ, Sedona, 16-Jun-1975 | FMNH | | |
| 543 | NVG-15113E05 | *Limenitis lorquini* | | M | USA: CA, Plumas Co., Butterfly Valley, 24-Jun-1993 | FMNH | | |
| 544 | NVG-15113D01 | *Limenitis archippus* | | M | USA: IL, Lake Co., Lake Forest, Skokie River Nature Preserve, Jun-2003 | FMNH | | |
| 545 | NVG-5141 | *Danaus plexippus* | | F | USA: TX, Hidalgo Co., Chihuahua RR tracks, 15-Nov-2015 | | | |
| 546 | NVG-5111 | *Danaus eresimus montezuma* | | F | USA: TX, Starr Co., Rio Grande City, 14-Nov-2015 | | | |
| 547 | NVG-3554 | *Danaus gilippus* | | F | USA: TX, Hidalgo Co., Old Rio Rico Rd. 1.5 air mi SE of Relampago, 13-Jun-2015 | | | |
| 548 | NVG-17118A03 | *Lycorea halia atergatis* | | | Mexico: Veracruz, 16-Apr-1979 | TAMU | | |
| 549 | NVG-17117H05 | *Dircenna klugii* | | F | Mexico: San Luis Potosi, 6-Feb-1976 | TAMU | | |
| 550 | NVG-17113G02 | *Phocides polybius lilea* | | F | USA: TX, Cameron Co., 20-Dec-1986 | TAMU | | |
| 551 | NVG-17097F12 | *Phocides belus* | | | Costa Rica, 2009 | USNM | | 09-SRNP-55421 |
| 552 | NVG-5316 | *Phocides pigmalion okeechobee* | | M | USA: FL, Monroe Co., Key West, 19-Dec-2015 | | | |
| 553 | NVG-5749 | *Cecropterus albociliatus albociliatus* | | M | Costa Rica: Guanacaste Prov., ACG, Sector Horizontes, ecl. on 08-Jun-2007 | USNM | NVG160217-10 | 07-SRNP-12984 |

| | | | | | | | | |
|---|---|---|---|---|---|---|---|---|
| 554 | NVG-17115E06 | *Cecropterus jalapus* | | F | Mexico: Sinaloa, Concordia, 26-Nov-2005 | CSUC | | CSU_ENT1045688 |
| 555 | NVG-3830 | *Cecropterus toxeus* | | F | USA: TX, Starr Co., Roma, 28-Jun-2015 | | | |
| 556 | NVG-3610 | *Cecropterus doryssus* | | M | USA: TX, Starr Co., Roma International Bridge, 14-Jun-2015 | | | |
| 557 | NVG-3311 | *Cecropterus lyciades* | | M | USA: TX, Sabine Co., Sabine NF, 12-Apr-2015 | | | |
| 558 | NVG-8269 | *Cecropterus casica* | | M | USA: AZ, Pinal Co., Coronado N.F., Catalina Mts., S of Oracle, 26-Mar-2017 | | | |
| 559 | 11-BOA-15609H02 | *Cecropterus tehuacana* | | M | Mexico: Coahuila, Cuesta La Muralla, 12-Sep-1976 | USNM | X-849 J. M. Burns 1978 | |
| 560 | NVG-4185 | *Cecropterus confusis* | | M | USA: TX, Wise Co., LBJ National Grassland, 19-Jul-2015 | | | |
| 561 | NVG-4539 | *Cecropterus bathyllus* | | F | USA: OK, Atoka Co., McGee Creek Recreation Area, 22-Aug-2015 | | | |
| 562 | NVG-14114A09 | *Cecropterus mexicana nevada* | | | USA: CA, Tulare Co., 6-Jul-2003 | LACM | | |
| 563 | NVG-17114A01 | *Cecropterus diversus* | | M | USA: CA, Madera Co., 30-May-2007 | CSUC | | CSU_ENT1036820 |
| 564 | NVG-3313 | *Cecropterus pylades* | | M | USA: TX, Sabine Co., Sabine NF, 12-Apr-2015 | | | |
| 565 | NVG-9688 | *Cecropterus drusius* | | F | USA: AZ, Santa Cruz Co., Coronado National Forest, Harshaw Creek Rd, 31-Jul-2017 | | | |
| 566 | NVG-14114A03 | *Cecropterus cincta* | | | Mexico: Hidalgo, Puerto del Caballo, 2-Sep-1982 | LACM | | |
| 567 | NVG-17109A11 | *Cecropterus pseudocellus* | ST | M | USA: AZ, Cochise Co., Huachuca Mts., 7-Jun-1910 | LACM | | |
| 568 | NVG-5075 | *Cecropterus egregius egregius* | | M | Costa Rica: Alajuela Prov., ACG, Sector Rincon Rain Forest, ecl. on 26-Mar-2013 | USNM | NVG151102-30 | 13-SRNP-40562 |
| 569 | NVG-4822 | *Cecropterus dorantes dorantes* | | | USA: FL, Collier Co., Chokoloskee Island, 2-Oct-2015 | | | |
| 570 | NVG-14112D08 | *Spicauda teleus* | | M | USA: Texas, Hidalgo Co., McAllen, 18-Oct-1973 | TAMU | | |
| 571 | NVG-14112D06 | *Spicauda tanna* | | M | Ecuador: Coco, 25-Jun-1980 | TAMU | | |
| 572 | NVG-3754 | *Spicauda procne* | | F | USA: TX, Hidalgo Co., Old Rio Rico Rd. 1.5 air mi SE of Relampago, 27-Jun-2015 | | | |
| 573 | NVG-17103C05 | *Spicauda simplicius* | | | Peru, 2-Feb-2013 | USNM | | |
| 574 | NVG-14112E09 | *Urbanus esmeraldus* | | M | USA: TX, Hidalgo Co., Estero Llano SP, 31-Oct-2011 | TLS | | |
| 575 | NVG-4894 | *Urbanus proteus proteus* | | | USA: FL, Miami-Dade Co., Homestead, 4-Oct-2015 | | | |
| 576 | NVG-14111G01 | *Urbanus pronus* | | | USA: TX, Hidalgo Co., SH1016 S of Mission, 19-Oct-1971 | TAMU | | |
| 577 | NVG-14121D06 | *Urbanus alva* | | | Belize: Cayo, Douglas de Silva, 7-Aug-2014 | JAShuey | | |
| 578 | NVG-5070 | *Urbanus viterboana* | | M | Costa Rica: Guanacaste Prov., ACG, Sector Cacao, ecl. on 12-May-2008 | USNM | NVG151102-25 | 08-SRNP-35218 |
| 579 | NVG-5074 | *Telegonus talus* | | M | Costa Rica: Guanacaste Prov., ACG, Sector Del Oro, ecl. on 28-May-2014 | USNM | NVG151102-29 | 14-SRNP-20343 |
| 580 | NVG-3427 | *Telegonus catemacoensis*-cf | | M | USA: TX, Cameron Co., 3 mi SW Sebastian, 30-May-2015 | | | |
| 581 | NVG-14111E04 | *Telegonus alector hoppferi* | | M | USA: TX, Hidalgo Co., Mission, 29-Oct-1971 | TAMU | | |
| 582 | NVG-14111E03 | *Telegonus alardus latia* | | M | USA: TX, Hidalgo Co., Santa Ana NWR, 31-Aug-1973 | TAMU | | |
| 583 | NVG-14111H04 | *Telegonus anaphus* | | M | USA: TX, Starr Co., Ft. Ringgold, 4-Nov-2014 | TAMU | | |
| 584 | NVG-14061D07 | *Telegonus cellus* | | | USA: AL, Marion Co., Hackleburg, 1-May-1974 | USNM | | |
| 585 | NVG-5715 | *Autochton potrillo potrillo* | | F | Costa Rica: Guanacaste Prov., ACG, Sector Mundo Nuevo, ecl. on 29-Aug-2009 | USNM | NVG160214-76 | 09-SRNP-57000 |
| 586 | NVG-3835 | *Spathilepia clonius* | | M | USA: TX, Starr Co., Roma International Bridge, 28-Jun-2015 | | | |
| 587 | NVG-17113G06 | *Proteides mercurius mercurius* | | M | Mexico: Tamaulipas, 19-Nov-1974 | TAMU | | |
| 588 | NVG-17108B08 | *Epargyreus zestos zestos* | | F | USA: FL, Monroe Co., Lower Matacumbe Key, 18-Feb-1982 | BMUW | | |
| 589 | NVG-4192 | *Epargyreus clarus clarus* | | F | USA: TX, Dallas Co., Dallas, White Rock Lake Park, 21-Jul-2015 | | | |
| 590 | NVG-16107G12 | *Epargyreus cruza* | | | Costa Rica, 2009 | USNM | | 09-SRNP-73319 |
| 591 | NVG-17097D12 | *Chioides zilpa* | | | USA: TX, Leakey, 20-Sep-1990 | USNM | | |
| 592 | NVG-3840 | *Chioides albofasciatus* | | M | USA: TX, Hidalgo Co., Old Rio Rico Rd. 1.5 air mi SE of Relampago, 28-Jun-2015 | | | |
| 593 | NVG-5526 | *Aguna metophis* | | M | USA: TX, Cameron Co., Brownsville, 20-Oct-1973 | TAMU | NVG160110-62 | |
| 594 | NVG-17098C02 | *Aguna asander* | | | Costa Rica, 2007 | USNM | | 07-SRNP-56519 |
| 595 | NVG-5525 | *Aguna claxon* | | F | USA: TX, Hidalgo Co., Loop 374, 6 mi west of Mission, 19-Oct-1973 | TAMU | | |
| 596 | NVG-3279 | *Codatractus alcaeus* | | M | USA: TX, Hidalgo Co., Penitas, 19-Oct-1973 | TAMU | NVG15011-95 | |
| 597 | NVG-9703 | *Codatractus arizonensis* | | M | USA: AZ, Santa Cruz Co., Coronado National Forest, Harshaw Creek Rd, 31-Jul-2017 | | | |
| 598 | NVG-15105B05 | *Lobotractus valeriana* | | M | USA: AZ, Santa Cruz Co., Harshaw Cr. Rd., 8-Aug-2005 | CAS | | |
| 599 | NVG-5680 | *Zestusa dorus* | | M | USA: AZ, Coconino Co., Oak Creek Canyon, 3-May-1984 | USNM | NVG160214-41 | |
| 600 | NVG-17113G12 | *Ectomis mexicanus* | | F | USA: TX, Cameron Co., Brownsville, 20-Oct-1973 | TAMU | | |
| 601 | NVG-18033D12 | *Ectomis octomaculata* | | | Mexico: Gomez Farias, 23-Oct-2002 | MWalker | | |
| 602 | NVG-5338 | *Polygonus leo histrio* | | M | USA: FL, Monroe Co., Key West, 19-Dec-2015 | | | |
| 603 | NVG-5029 | *Polygonus savigny savigny* | | | Mexico: Yucatan, 15-Aug-1962 | USNM | NVG151101-80 | |
| 604 | NVG-17113G10 | *Cogia undulatus* | | M | USA: TX, Hidalgo Co., Relampago, 12-Oct-1975 | TAMU | | |
| 605 | NVG-3354 | *Cogia calchas* | | M | USA: TX, Hidalgo Co., Old Rio Rico Rd. 1.5 air mi SE of Relampago, 23-May-2015 | | | |
| 606 | NVG-18033E12 | *Cogia caicus moschus* | | | Mexico: Sonora, 15-Sep-2010 | MWalker | | |
| 607 | NVG-17113D06 | *Cogia hippalus hippalus* | | | Mexico: Sonora, 1987 | TMMC | | |
| 608 | NVG-17116A05 | *Cogia outis* | | M | USA: TX, Cooke Co., 28-Jul-1973 | TAMU | | |
| 609 | NVG-17116A02 | *Celaenorrhinus fritzgaertneri* | | F | Mexico: Tamaulipas, 1-Feb-1975 | TAMU | | |

| | | | | | | | | |
|---|---|---|---|---|---|---|---|---|
| 610 | NVG-18013G09 | *Celaenorrhinus stallingsi* | | | Costa Rica: Guanacaste Prov., ACG, Sector Del Oro, ecl. on 11-Jan-2011 | USNM | | 10-SRNP-22671 |
| 611 | NVG-4469 | *Apyrrothrix araxes arizonae* | | F | USA: AZ, Santa Cruz Co., NF Road 58, at 9 miles se of Patagonia, 11-Aug-2015 | | | |
| 612 | NVG-7880 | *Grais stigmaticus* | | | Costa Rica: Guanacaste Prov., ACG, Sector Pitilla, ecl. on 16-Mar-2014 | USNM | NVG170206-65 | 14-SRNP-30242 |
| 613 | NVG-1931 | *Eantis pallida* | | M | Mexico: Tamaulipas, Villa Gomez Farias, 27-Jan-1974 | TAMU | NVG140104-70 | |
| 614 | NVG-3758 | *Eantis tamenund* | | F | USA: TX, Hidalgo Co., Old Rio Rico Rd. 1.5 air mi SE of Relampago, 27-Jun-2015 | | | |
| 615 | NVG-5485 | *Arteurotia tractipennis tractipennis* | | M | USA: TX, Hidalgo Co., Mission, 2-Sep-1972 | TAMU | NVG160110-26 | |
| 616 | NVG-17116A07 | *Nisoniades rubescens* | | M | Mexico: Tamaulipas, 26-Jan-1974 | TAMU | | |
| 617 | NVG-17116A09 | *Pellicia dimidiata dimidiata* | | M | Mexico: Tamaulipas, 8-Jan-1974 | TAMU | | |
| 618 | NVG-7877 | *Pellicia arina* | | | Costa Rica: Alajuela Prov., ACG, Sector Rincon Rain Forest, ecl. on 18-Mar-2013 | USNM | NVG170206-62 | 13-SRNP-67590 |
| 619 | NVG-17108F02 | *Clytius clytius* | | F | USA: AZ, Santa Cruz Co., 4-May-1991 | LACM | | |
| 620 | NVG-7245 | *Bolla brennus brennus* | | M | Panama: Chiriqui Prov., El Volcan, 16-Apr-1973 | USNM | NVG161005-72 | |
| 621 | NVG-9694 | *Staphylus ceos* | | F | USA: AZ, Santa Cruz Co., Coronado National Forest, Harshaw Creek Rd, 31-Jul-2017 | | | |
| 622 | NVG-7039 | *Staphylus hayhurstii* | | F | USA: TX, Delta Co., 5 air mi WNW of Commerce, 24-Sep-2016 | | | |
| 623 | NVG-18058E10 | *Staphylus mazans* | | M | Mexico: Veracruz, 23-May-1979 | USNM | | |
| 624 | NVG-5594 | *Hesperopsis alpheus alpheus* | | F | USA: TX, Cameron Co., E of Brownsville, 8-Feb-2016 | | | |
| 625 | NVG-17067B02 | *Hesperopsis gracielae* | | | USA: CA, Riverside Co., Blythe, 15-Apr-1997 | CSUC | | CSU_ENT1039348 |
| 626 | NVG-17067A09 | *Hesperopsis libya libya* | | | USA: CA, Inyo Co., Argus Mts., Homewood Canyon, 24-Aug-2009 | CSUC | | CSU_ENT1039161 |
| 627 | NVG-7976 | *Pholisora mejicanus* | | | USA: NM, Colfax Co., Cimarron, 7-Aug-1989 | USNM | NVG170207-61 | |
| 628 | NVG-3990 | *Pholisora catullus* | | F | USA: TX, Starr Co., Roma International Bridge, 10-Jul-2015 | | | |
| 629 | NVG-17116A10 | *Noctuana stator* | | F | Mexico: San Luis Potosi, 17-Nov-1974 | TAMU | | |
| 630 | NVG-15102D03 | *Windia windi* | | M | Mexico: Sonora, Rte 16, 8.5 mi W. Rio Yaqui, 26-Aug-1984 | USNM | X-2052 J. M. Burns 1985 | |
| 631 | NVG-6008 | *Systasea zampa* | | F | USA: TX, El Paso Co., El Paso, Franklin Mts. State Park, 31-Mar-2016 | | | |
| 632 | NVG-3621 | *Systasea pulverulenta* | | M | USA: TX, Duval Co., Parilla Creek, North Fork at SH16, 14-Jun-2015 | | | |
| 633 | NVG-17116B06 | *Canesia canescens canescens* | | F | Mexico: Tamaulipas, 28-Jan-1974 | TAMU | | |
| 634 | NVG-7906 | *Xenophanes tryxus* | | M | Costa Rica: Alajuela Prov., ACG, Sector San Cristobal, 26-Mar-2010 | USNM | NVG170206-91 | 10-SRNP-103428 |
| 635 | NVG-10646 | *Antigonus emorsa* | | M | Mexico: Michoacan, 21-Aug-1994 | TAMU | NVG180106-63 | |
| 636 | NVG-7907 | *Antigonus erosus* | | | Costa Rica: Guanacaste Prov., ACG, Sector Mundo Nuevo, ecl. on 18-Oct-2013 | USNM | NVG170206-92 | 13-SRNP-56479 |
| 637 | 11-BOA-13383BrockB12 | *Celotes limpia* | | F | USA: TX, Brewster Co., 2005 | JimPBrock | | |
| 638 | NVG-3956 | *Celotes nessus* | | M | USA: TX, Hidalgo Co., Penitas RR tracks, 28-Jun-2015 | | | |
| 639 | NVG-17069B07 | *Pyrgus centaureae freija* | | M | USA: AK, Dalton Hwy, mi 274, 5-Jul-1991 | USNM | | |
| 640 | NVG-17067G07 | *Pyrgus ruralis ruralis* | | | USA: CA, Mariposa Co., Miami Mountain Rd., Hwy 41, 14-Jun-2009 | CSUC | | CSU_ENT1030397 |
| 641 | NVG-17067H09 | *Pyrgus xanthus* | | | USA: CO, San Juan Co., Cunningham Creek, below Road 4A, 23-Jun-2002 | CSUC | | CSU_ENT1036166 |
| 642 | PAO-187 | *Pyrgus scriptura* | | | USA: UT, Garfield Co., Grand Staircase - Escalante National Monument, 5-Aug-2016 | | | |
| 643 | NVG-18018E04 | *Burnsius communis* | | M | USA: AZ, Cochise Co., Portal, 11-Jul-1974 | USNM | | |
| 644 | NVG-18018E01 | *Burnsius albescens* | | M | USA: AZ, Cochise Co., Portal, 17-Jul-1974 | USNM | | |
| 645 | NVG-3375 | *Burnsius philetas* | | M | USA: TX, Starr Co., 0.5 W Roma Creek, along FM650, 23-May-2015 | | | |
| 646 | NVG-3542 | *Burnsius oileus* | | M | USA: TX, Hidalgo Co., Old Rio Rico Rd. 1.5 air mi SE of Relampago, 13-Jun-2015 | | | |
| 647 | NVG-5229 | *Heliopetes domicella domicella* | | M | USA: TX, Starr Co., Rio Grande City, 26-Nov-2015 | | | |
| 648 | NVG-14114E04 | *Heliopetes sublinea* | | M | USA: TX, Hidalgo Co., Alamo, 1-Nov-2014 | TLS | | |
| 649 | 11-BOA-13385C12 | *Heliopetes ericetorum* | | M | USA: AZ, Gila Co., Washington Park, 24-Apr-2012 | | | |
| 650 | NVG-3338 | *Heliopetes laviana laviana* | | F | USA: TX, Cameron Co., River Dr., 1.4 mi S of Santa Maria, 23-May-2015 | | | |
| 651 | NVG-5250 | *Heliopetes macaira macaira* | | F | USA: TX, Cameron Co., E of Brownsville, 27-Nov-2015 | | | |
| 652 | NVG-7557 | *Heliopetes arsalte* | | M | USA: TX, Cameron Co., Boca Chica, 20-Oct-1973 | TAMU | NVG170107-13 | |
| 653 | NVG-17109G07 | *Heliopetes alana* | | M | Guatemala, 5-10-Jun-2003 | LACM | | |
| 654 | NVG-5494 | *Sostrata nordica* | | M | USA: TX, Hidalgo Co., 26-Oct-1973 | TAMU | NVG160110-35 | |
| 655 | NVG-7884 | *Echelatus sempiternus* | | | Costa Rica: Guanacaste Prov., ACG, Sector Santa Elena, ecl. on 16-Mar-2007 | USNM | NVG170206-69 | 07-SRNP-12147 |
| 656 | NVG-7882 | *Hoodus pelopidas* | | | Costa Rica: Guanacaste Prov., ACG, Sector Mundo Nuevo, ecl. on 09-Mar-2008 | USNM | NVG170206-67 | 08-SRNP-55556 |
| 657 | NVG-5493 | *Gorgythion begga pyralina* | | M | USA: TX, Hidalgo Co., 28-Dec-1973 | TAMU | NVG160110-34 | |
| 658 | NVG-5100 | *Chiothion georgina* | | F | USA: TX, Starr Co., Rio Grande City, 14-Nov-2015 | | | |
| 659 | NVG-17109F12 | *Chiomara mithrax* | | F | USA: AZ, Santa Cruz Co., 19-Sep-1992 | LACM | | |
| 660 | NVG-17116A12 | *Timochares ruptifasciata* | | F | USA: TX, Bexar Co., 17-Jul-1997 | TAMU | | |
| 661 | NVG-17095E06 | *Ephyriades brunnea floridensis* | | | USA: FL, Monroe Co., Stock Island, 18-Mar-1987 | USNM | | |
| 662 | NVG-6151 | *Erynnis icelus* | | F | USA: VA, Rockbridge Co., 7 air mi NNW of Lexington, 11-May-2016 | | | |
| 663 | NVG-6120 | *Erynnis brizo* | | F | USA: WV, Pendleton Co., FS112, 8-May-2016 | | | |
| 664 | NVG-8904 | *Gesta telemachus* | | M | USA: NM, Santa Fe Co., Santa Fe National Forest, 15-May-2017 | | | |
| 665 | NVG-16107D08 | *Gesta juvenalis juvenalis* | | M | USA: TX, Jeff Davis Co., Davis Mts., 19-Apr-2015 | JSCarter | | |

| | | | | | | | | |
|---|---|---|---|---|---|---|---|---|
| 666 | PAO-21 | *Gesta propertius* | | | USA: CA, Sierra Co., N fork Yuba River, 11-Jun-2016 | | | |
| 667 | NVG-18022E03 | *Gesta meridianus meridianus* | HT | M | USA: AZ, White Mts., 1-Jul-1915 | AMNH | | |
| 668 | NVG-3678 | *Gesta horatius* | | F | USA: TX, Marion Co., Caddo Lake region, 20-Jun-2015 | | | |
| 669 | NVG-9715 | *Gesta tristis tatius* | | M | USA: TX, Jeff Davis Co., Davis Mnts., 4-Aug-2017 | | | |
| 670 | NVG-4753 | *Gesta zarucco* | | M | USA: FL, Levy Co., NW of Williston Highlands, along CR337, 27-Sep-2015 | | | |
| 671 | NVG-4208 | *Gesta funeralis* | | F | USA: TX, Collin Co., Richardson, 24-Jul-2015 | | | |
| 672 | NVG-3907 | *Gesta baptisiae* | | M | USA: AR, Montgomery Co., Ouachita National Forest, 4-Jul-2015 | | | |
| 673 | NVG-18014B07 | *Gesta lucilius* | | M | USA: MI, Presque Isle Co., 25-May-1995 | USNM | | |
| 674 | NVG-17114A03 | *Gesta afranius* | | M | USA: CO, Montezuma Co., 27-Jun-1999 | CSUC | | CSU_ENT1034909 |
| 675 | NVG-6567 | *Gesta persius fredericki* | | F | USA: CO, Park Co., 3 air mi SW Fairplay, 10-Jul-2016 | | | |
| 676 | NVG-9725 | *Gesta scudderi* | | M | USA: AZ, Cochise Co., Huachuca Mtns., Coronado National Forest, 3-Aug-2017 | | | |
| 677 | NVG-9770 | *Gesta pacuvius pacuvius* | | M | USA: AZ, Cochise Co., Chiricahua Mtns., Coronado National Forest, 3-Aug-2017 | | | |
| 678 | NVG-3900 | *Gesta martialis* | | M | USA: AR, Montgomery Co., Ouachita National Forest, 4-Jul-2015 | | | |
| 679 | NVG-6818 | *Gesta invisus* | | M | USA: TX, Aransas Co., Rockport, 21-Aug-2016 | | | |
| 680 | NVG-16106E03 | *Carterocephalus palaemon mandan* | | F | USA: MN, Lake of the Woods Co., Norris Camp, 5-Jun-2016 | WDempwolf | | |
| 681 | NVG-17068A11 | *Piruna aea mexicana* | | | USA: AZ, Santa Cruz Co., Ruby Road, 1-2 Miles W of Calir Culch Road, 27-Aug-2016 | CSUC | | CSU_ENT1033276 |
| 682 | NVG-17114A05 | *Piruna penaea* | | M | USA: AZ, Santa Cruz Co., 27-Aug-2016 | CSUC | | CSU_ENT1033276 |
| 683 | NVG-10649 | *Piruna polingii* | | M | Mexico: Nuevo Leon, 23-Oct-1979 | TAMU | NVG180106-66 | |
| 684 | NVG-6454 | *Piruna pirus* | | M | USA: CO, Grand Co., 4 air mi SSE Hot Sulphur Springs, 6-Jul-2016 | | | |
| 685 | NVG-492 | *Piruna haferniki* | | | USA: TX, Brewster Co., Big Bend National Park, 15-Sep-2007 | | | |
| 686 | NVG-7910 | *Erionota thrax* | | | USA: HI, Molokai, Kamakou Preserve, 2-Nov-2005 | USNM | NVG170206-95 | |
| 687 | NVG-8176 | *Euphyes pilatka pilatka* | | F | USA: FL, Miami-Dade Co., 4 mi SW of Florida City, 19-Mar-2017 | | | |
| 688 | NVG-17116E01 | *Euphyes berryi* | | M | USA: FL, Volusia Co., 23-Sep-1993 | TAMU | | |
| 689 | NVG-15113F07 | *Euphyes dukesi* | | M | USA: TX, Tyler Co., Steinhagen lake, 11-Sep-1993 | FMNH | | |
| 690 | NVG-4534 | *Euphyes dion* | | F | USA: TX, Lamar Co., FM1499 at Craddock Creek, 20-Aug-2015 | | | |
| 691 | NVG-932 | *Euphyes bayensis* | | | USA: LA, Cameron Pa., Grand Chenier, 1-Oct-2011 | | | |
| 692 | NVG-17114B08 | *Euphyes conspicua* | | M | USA: IA, Johnson Co., 10-Jul-1984 | CSUC | | CSU_ENT1023047 |
| 693 | NVG-17114B10 | *Euphyes bimacula* | | M | USA: CO, Larimer Co., 21-Jun-1986 | CSUC | | CSU_ENT1038759 |
| 694 | NVG-17114B11 | *Euphyes arpa* | | M | USA: GA, Mcintosh Co., 23-Aug-2014 | CSUC | | CSU_ENT1038765 |
| 695 | NVG-7581 | *Euphyes vestris kiowah* | | M | USA: TX, Brewster Co., Big Bend National Park, 23-Sep-1971 | TAMU | NVG170107-37 | |
| 696 | NVG-17098F11 | *Notamblyscirtes simius* | | F | USA: CO, Larimer Co., 14-Jun-1987 | USNM | | |
| 697 | NVG-3649 | *Quasimellana eulogius* | | F | USA: TX, Cameron Co., River Dr., 1.4 mi S of Santa Maria, 14-Jun-2015 | | | |
| 698 | NVG-17119D04 | *Anatrytone mazai* | | | Costa Rica, Oct-1982 | USNM | | |
| 699 | NVG-4059 | *Anatrytone logan logan* | | F | USA: TX, Dallas Co., Dallas, Norbuck Park, 14-Jul-2015 | | | |
| 700 | NVG-17095H05 | *Atrytone arogos iowa* | | F | USA: KS, Barber Co., 29-Aug-1992 | USNM | | |
| 701 | NVG-17095H06 | *Atrytone bulenta* | | M | USA: NJ, Cumberland Co., 28-Jun-1998 | USNM | | |
| 702 | NVG-3986 | *Atrytone byssus kumskaka* | | F | USA: AR, Montgomery Co., Ouachita National Forest, 4-Jul-2015 | | | |
| 703 | NVG-5174 | *Hylephila phyleus* | | F | USA: TX, Hidalgo Co., Chihuahua RR tracks, 15-Nov-2015 | | | |
| 704 | NVG-4942 | *Hedone vibex praeceps* | | F | USA: TX, Hidalgo Co., Old Rio Rico Rd. 1.5 air mi SE of Relampago, 13-Nov-2015 | | | |
| 705 | NVG-9320 | *Limochores sonora* | | F | USA: WY, Park Co., Yellowstone National Park, 24-Jul-2017 | | | |
| 706 | NVG-18042G01 | *Limochores mystic* | | F | USA: WI, Dunn Co., Muddy Creek State WMA, 13-Jun-2016 | WDempwolf | | WRD 10960 |
| 707 | NVG-4547 | *Limochores origenes origenes* | | M | USA: OK, Atoka Co., McGee Creek Natural Scenic Recreation Area, 22-Aug-2015 | | | |
| 708 | NVG-8978 | *Polites themistocles turneri* | | | USA: NM, Otero Co., Lincoln National Forest, 21-May-2017 | | | |
| 709 | NVG-4276 | *Polites peckius peckius* | | F | USA: IN, Montgomery Co., 1.5 mi south of Deer Mill, 1-Aug-2015 | | | |
| 710 | NVG-17114B05 | *Polites mardon* | | M | USA: CA, Del Norte Co., 27-Aug-1992 | CSUC | | CSU_ENT1034299 |
| 711 | PAO-395 | *Polites sabuleti sabuleti* | | | USA: CA, Monterey Co., Pacific Grove, 16-Jun-2017 | | | |
| 712 | NVG-6474 | *Polites draco* | | M | USA: CO, Grand Co., 4.3 air mi SE Hot Sulphur Springs, 6-Jul-2016 | | | |
| 713 | NVG-5968 | *Polites carus* | | F | USA: AZ, Santa Cruz Co., Pajarito Mtns., 30-Mar-2016 | | | |
| 714 | NVG-17114B04 | *Polites rhesus* | | M | USA: CO, Larimer Co., 22-May-1996 | CSUC | | CSU_ENT1025525 |
| 715 | NVG-8170 | *Polites baracoa baracoa* | | M | USA: FL, Miami-Dade Co., 4.8 air mi E of Florida City, 19-Mar-2017 | | | |
| 716 | NVG-3987 | *Wallengrenia egeremet* | | F | USA: AR, Montgomery Co., Ouachita National Forest, 4-Jul-2015 | | | |
| 717 | NVG-3571 | *Wallengrenia otho clavus* | | F | USA: TX, Cameron Co., 3 mi SW Sebastian, 13-Jun-2015 | | | |
| 718 | NVG-4070 | *Nyctelius nyctelius* | | F | USA: TX, Hidalgo Co., Old Rio Rico Rd. 1.5 air mi SE of Relampago, 11-Jul-2015 | | | |
| 719 | NVG-7963 | *Conga chydaea* | | | Costa Rica: Alajuela Prov., ACG, Sector Rincon Rain Forest, ecl. on 11-Dec-2009 | USNM | NVG170207-48 | 09-SRNP-68418 |
| 720 | NVG-18014H01 | *Vernia verna* | | M | USA: OH, Summit Co., Liberty Metropark S of Rt.82, 8-Jun-2012 | USNM | | |
| 721 | NVG-3718 | *Atalopedes campestris huron* | | F | USA: TX, Dallas Co., Dallas, 23-Jun-2015 | | | |

| | | | | | | | | |
|---|---|---|---|---|---|---|---|---|
| 722 | NVG-16108F07 | *Hesperia comma mixta* | | | Russia: Tien-Shan, 26-Jul-1991 | USNM | | |
| 723 | NVG-17107E01 | *Hesperia colorado oregonia* | | M | USA: WA, Thurston Co., Mima Mounds National Area Preserve, 9-Aug-1992 | BMUW | | |
| 724 | NVG-18027B08 | *Hesperia assiniboia* | PLT | M | Canada: Saskatchewan, Regina, 5-Aug-1890 | USNM | | |
| 725 | PAO-14 | *Hesperia juba* | | | USA: CA, Sierra Co., S of Loyalton, 9-Jun-2016 | | | |
| 726 | PAO-364 | *Hesperia lindseyi lindseyi* | | | USA: CA, Colusa Co., Mendocino National Forest, 18-May-2017 | | | |
| 727 | NVG-17114B03 | *Hesperia miriamae miriamae* | | M | USA: CA, Inyo Co., 13-Aug-2006 | CSUC | | CSU_ENT1028355 |
| 728 | NVG-6612 | *Hesperia nevada nevada* | | F | USA: CO, Lake Co., 1.3 air mi NE Twin Lakes, 11-Jul-2016 | | | |
| 729 | NVG-17113E02 | *Hesperia woodgatei* | | F | USA: AZ, Yavapai Co., 23-Sep-2015 | WDempwolf | | |
| 730 | PAO-114 | *Hesperia uncas* | | | USA: WY, Sweetwater Co., Pilot Butte Rd., 2-Jul-2016 | | | |
| 731 | NVG-17116D01 | *Hesperia leonardus montana* | | M | USA: CO, Jefferson Co., 19-Aug-1987 | TAMU | | |
| 732 | PAO-368 | *Hesperia columbia* | | | USA: CA, Colusa Co., Mendocino National Forest, 18-May-2017 | | | |
| 733 | NVG-8279 | *Hesperia pahaska williamsi* | | M | USA: AZ, Pinal Co., Coronado N.F., Catalina Mts., S of Oracle, 26-Mar-2017 | | | |
| 734 | NVG-8392 | *Hesperia viridis* | | M | USA: TX, Blanco Co., Pedernales Falls State Park, 31-Mar-2017 | | | |
| 735 | NVG-17116D04 | *Hesperia metea licinus* | | F | USA: TX, Wheeler Co., 25-Apr-1987 | TAMU | | |
| 736 | NVG-4650 | *Hesperia attalus slossonae* | | M | USA: FL, Citrus Co., Withlacoochie State Forest, 25-Sep-2015 | | | |
| 737 | NVG-17114A10 | *Hesperia ottoe* | | M | USA: CO, Boulder Co., 10-Jul-2008 | CSUC | | CSU_ENT1032139 |
| 738 | NVG-6510 | *Hesperia meskei straton* | | M | USA: FL, Levy Co., NW of Williston Highlands, 7-Sep-2015 | | | |
| 739 | NVG-17114B01 | *Hesperia sassacus* | | M | USA: ME, Cumberland Co., 7-Jul-1982 | CSUC | | CSU_ENT1028320 |
| 740 | NVG-17114A12 | *Hesperia dacotae* | | M | USA: ND, Mchenry Co., 28-Jul-2006 | CSUC | | CSU_ENT1028253 |
| 741 | NVG-8050 | *Pseudocopaeodes eunus* | | F | USA: CA, Inyo Co., 20-Jun-1950 | USNM | NVG170208-35 | |
| 742 | NVG-17114B06 | *Ochlodes yuma* | | M | USA: CO, Mesa Co., 6-Sep-2001 | CSUC | | CSU_ENT1024498 |
| 743 | PAO-263 | *Ochlodes sylvanoides* | | | USA: CO, Larimer Co., Roosevelt National Forest, 10-Sep-2016 | | | |
| 744 | PAO-23 | *Ochlodes agricola* | | | USA: CA, Sierra Co., N fork Yuba River, 11-Jun-2016 | | | |
| 745 | NVG-6016 | *Stinga morrisoni* | | M | USA: TX, Jeff Davis Co., Davis Mts., 31-Mar-2016 | | | |
| 746 | NVG-7046 | *Poanes yehl* | | F | USA: TX, Hopkins Co., Cooper Lake State Park, 24-Sep-2016 | | | |
| 747 | NVG-4704 | *Poanes aaroni howardi* | | F | USA: FL, Levy Co., Yankeetown, 26-Sep-2015 | | | |
| 748 | NVG-6711 | *Poanes viator zizaniae* | | F | USA: TX, Dallas Co., Dallas, White Rock Lake Park, 26-Jul-2016 | | | |
| 749 | NVG-17114B07 | *Poanes massasoit* | | M | USA: MD, Dorchester Co., 2-Jul-1976 | CSUC | | CSU_ENT1031028 |
| 750 | NVG-6175 | *Lon hobomok hobomok* | | M | USA: VA, Augusta Co., Calfpasture River, 11-May-2016 | | | |
| 751 | NVG-9209 | *Lon zabulon* | | F | USA: AR, Montgomery Co., Ouachita National Forest, 6-Jul-2017 | | | |
| 752 | NVG-9774 | *Lon taxiles* | | M | USA: AZ, Cochise Co., Chiricahua Mtns., Coronado National Forest, 3-Aug-2017 | | | |
| 753 | NVG-17111H01 | *Lon melane melane* | | F | USA: CA, San Luis Obispo Co., 4-Apr-1994 | LACM | | |
| 754 | PAO-148 | *Paratrytone snowi* | | | USA: CO, Gilpin Co., Golden Gate Canyon State Park, 24-Jul-2016 | | | |
| 755 | NVG-8159 | *Oligoria maculata* | | F | USA: FL, Miami-Dade Co., 4.8 air mi E of Florida City, 18-Mar-2017 | | | |
| 756 | NVG-3761 | *Oligoria percosius* | | F | USA: TX, Hidalgo Co., Old Rio Rico Rd. 1.5 air mi SE of Relampago, 27-Jun-2015 | | | |
| 757 | NVG-17102D10 | *Atrytonopsis quinteri* | HT | M | USA: NC, Onslow Co., Hammocks Beach State Park, 28-Jul-1983 | USNM | | |
| 758 | NVG-14065B04 | *Atrytonopsis loammi* | | F | USA: FL, Duval Co., Jacksonville, 22-Aug-1993 | JAScott | | |
| 759 | NVG-8567 | *Atrytonopsis hianna turneri* | | F | USA: OK, Atoka Co., 2.4 air mi ESE Stringtown, 15-Apr-2017 | | | |
| 760 | NVG-14065B03 | *Atrytonopsis deva* | | F | USA: AZ, Pima Co., 17-Apr-1972 | JAScott | | |
| 761 | NVG-11172 | *Atrytonopsis vierecki* | | M | USA: NM, Otero Co., Lincoln National Forest, 19-May-2018 | | | |
| 762 | NVG-17111H08 | *Atrytonopsis lunus* | | M | USA: AZ, Cochise Co., 5-Aug-1979 | LACM | | |
| 763 | NVG-11324 | *Atrytonopsis margarita* | | F | USA: NM, Santa Fe Co., Santa Fe National Forest, 21-May-2018 | | | |
| 764 | NVG-12265 | *Atrytonopsis python* | | | USA: AZ, Cochise Co., Chiricahua Mtns. FS42, 27-May-2019 | | | |
| 765 | NVG-17111H09 | *Atrytonopsis cestus* | | M | USA: AZ, Graham Co., 21-Apr-1990 | LACM | | |
| 766 | NVG-5995 | *Atrytonopsis pittacus* | | F | USA: AZ, Santa Cruz Co., Pajarito Mtns., 30-Mar-2016 | | | |
| 767 | NVG-11277 | *Atrytonopsis edwardsi* | | F | USA: TX, Jeff Davis Co., 18-May-2018 | | | |
| 768 | NVG-18012A08 | *Thespieus macareus* | | | Mexico: Oaxaca, Jul-Sep-1992 | USNM | | |
| 769 | NVG-17116C08 | *Rhinthon osca* | | M | USA: TX, Hidalgo Co., 24-Oct-1974 | TAMU | | |
| 770 | NVG-18012C11 | *Niconiades nikko* | | | Mexico: San Luis Potosi, 15-Oct-1976 | USNM | | |
| 771 | NVG-10672 | *Amblyscirtes carolina* | | M | USA: VA, Chesapeake, 20-Jun-1970 | TAMU | NVG180106-89 | |
| 772 | NVG-10673 | *Amblyscirtes reversa* | | M | USA: VA, Virginia Beach, 8-Aug-1971 | TAMU | NVG180106-90 | |
| 773 | NVG-3524 | *Amblyscirtes aesculapius* | | F | USA: TX, San Jacinto Co., Sam Houston NF, Big Creek Scenic Area, 7-Jun-2015 | | | |
| 774 | NVG-10674 | *Amblyscirtes hegon* | | M | USA: TX, Smith Co., Tyler State Park, 15-Mar-1986 | TAMU | NVG180106-91 | |
| 775 | NVG-7646 | *Amblyscirtes nysa* | | M | USA: TX, Terrell Co., USH90, 10 mi W of Sanderson, 5-Sep-1976 | TAMU | NVG170107-77 | |
| 776 | NVG-7614 | *Amblyscirtes alternata* | | M | USA: TX, Leon Co., 3 mi W Buffalo, 29-May-1974 | TAMU | NVG170108-67 | |
| 777 | NVG-7612 | *Amblyscirtes vialis* | | M | USA: TX, Brown Co., Lake Brownwood State Park, 11-Apr-1978 | TAMU | NVG170108-65 | |

| | | | | | | | | |
|---|---|---|---|---|---|---|---|---|
| 778 | NVG-8572 | *Amblyscirtes linda* | | F | USA: OK, Atoka Co., McGee Creek Natural Scenic Recreation Area, 15-Apr-2017 | | | |
| 779 | NVG-810 | *Amblyscirtes aenus* | | | USA: TX, Blanco Co., Pedernales Falls State Park , 2-Apr-2011 | | | |
| 780 | NVG-10656 | *Amblyscirtes cassus* | | M | USA: TX, Jeff Davis Co., 11-Jul-1969 | TAMU | NVG180106-73 | |
| 781 | NVG-10659 | *Amblyscirtes texanae* | | M | USA: TX, Jeff Davis Co., 11-Jul-1969 | TAMU | NVG180106-76 | |
| 782 | NVG-3699 | *Amblyscirtes belli* | | M | USA: TX, Dallas Co., Dallas, 21-Jun-2015 | | | |
| 783 | NVG-1671 | *Amblyscirtes celia* | | | USA: TX, Travis Co., Platt Lane, 16-Apr-2011 | WDempwolf | | |
| 784 | NVG-17114A07 | *Amblyscirtes tolteca prenda* | | M | Mexico: Sonora, Yecora, 29-Jul-1997 | CSUC | | CSU_ENT1028834 |
| 785 | NVG-9693 | *Amblyscirtes nereus* | | M | USA: AZ, Santa Cruz Co., Coronado National Forest, Harshaw Creek Rd, 31-Jul-2017 | | | |
| 786 | NVG-7643 | *Amblyscirtes eos* | | F | USA: TX, Tarrant Co., Benbrook Reservoir, 5-Sep-1976 | TAMU | NVG170107-74 | |
| 787 | NVG-9685 | *Amblyscirtes elissa arizonae* | | F | USA: AZ, Santa Cruz Co., Coronado National Forest, Harshaw Creek Rd, 31-Jul-2017 | | | |
| 788 | NVG-7604 | *Amblyscirtes oslari* | | M | USA: TX, Lubbock Co., Buffalo Springs Lake, 22-May-1976 | TAMU | NVG170108-57 | |
| 789 | NVG-9727 | *Amblyscirtes exoteria* | | F | USA: AZ, Cochise Co., Chiricahua Mtns., Coronado National Forest, 3-Aug-2017 | | | |
| 790 | NVG-18037G10 | *Amblyscirtes phylace* | | | USA: NM, Valencia Co., Zuni Mts., Pole Camp, 29-Jun-1977 | CSUC | | CSU-ENT 1005459 |
| 791 | NVG-17114A09 | *Amblyscirtes fimbriata fimbriata* | | M | USA: AZ, Cochise Co., 28-Jul-2005 | CSUC | | CSU_ENT1038258 |
| 792 | NVG-3924 | *Nastra lherminier* | | F | USA: AR, Montgomery Co., Ouachita National Forest, 4-Jul-2015 | | | |
| 793 | NVG-4747 | *Nastra neamathla* | | F | USA: FL, Levy Co., NW of Williston Highlands, along CR337, 27-Sep-2015 | | | |
| 794 | NVG-3478 | *Nastra julia* | | M | USA: TX, Duval Co., Parilla Creek, North Fork at SH16, 31-May-2015 | | | |
| 795 | NVG-17111E05 | *Nastra perigenes* | | M | USA: TX, Cameron Co., Brownsville, 21-Oct-1963 | LACM | | |
| 796 | NVG-4062 | *Lerodea eufala* | | F | USA: TX, Dallas Co., Dallas, Norbuck Park, 14-Jul-2015 | | | |
| 797 | NVG-5380 | *Lerodea arabus* | | F | USA: TX, Starr Co., Roma, 24-Dec-2015 | | | |
| 798 | NVG-17111E07 | *Monca crispinus* | | M | USA: TX, Hidalgo Co., 20-May-1972 | LACM | | |
| 799 | NVG-10654 | *Troyus fantasos* | | M | USA: TX, Hidalgo Co., Penitas, 24-Oct-1975 | TAMU | NVG180106-71 | |
| 800 | NVG-4842 | *Cymaenes tripunctus tripunctus* | | F | USA: FL, Collier Co., Copeland, 2-Oct-2015 | | | |
| 801 | NVG-3401 | *Cymaenes trebius* | | F | USA: TX, Hidalgo Co., Old Rio Rico Rd. 1.5 air mi SE of Relampago, 30-May-2015 | | | |
| 802 | NVG-4792 | *Lerema accius* | | F | USA: FL, Levy Co., Gulf Hammock, 27-Sep-2015 | | | |
| 803 | NVG-3194 | *Lerema liris* | | M | Mexico: Tamaulipas, Ciudad Mante, 19-Dec-1973 | TAMU | NVG15011-10 | |
| 804 | NVG-7968 | *Mnasilus allubita* | | | Costa Rica: Guanacaste Prov., ACG, Sector Santa Elena, ecl. on 23-Sep-2002 | USNM | NVG170207-53 | 02-SRNP-13739 |
| 805 | NVG-10653 | *Corticea corticea* | | M | USA: TX, Hidalgo Co., SH1016 S of Mission nr. Madero, 4-Nov-1973 | TAMU | NVG180106-70 | |
| 806 | NVG-4881 | *Asbolis capucinus* | | | USA: FL, Monroe Co., Key West, 3-Oct-2015 | | | |
| 807 | NVG-17111D11 | *Synapte pecta* | | M | USA: TX, Hidalgo Co., Pharr, 22-Sep-1945 | LACM | | |
| 808 | NVG-10652 | *Synapte salenus salenus* | | M | Mexico: San Luis Potosi, 21-Feb-1976 | TAMU | NVG180106-69 | |
| 809 | NVG-10671 | *Synapte shiva* | | M | Mexico: Chiapas, 13-Aug-1967 | TAMU | NVG180106-88 | |
| 810 | NVG-7988 | *Adopaeoides prittwitzi* | | M | USA: AZ, Santa Cruz Co., San Rafael Valley, 14-Aug-1999 | USNM | NVG170207-73 | |
| 811 | NVG-4461 | *Ancyloxypha numitor* | | F | USA: TX, Dallas Co., Dallas, 10-Aug-2015 | | | |
| 812 | NVG-17068D03 | *Ancyloxypha arene* | | | USA: AZ, Maricopa Co., S. Shore Apache Lake near Boat Launch, 20-Apr-2013 | CSUC | | CSU_ENT1025075 |
| 813 | NVG-9438 | *Thymelicus lineola lineola* | | M | USA: WY, Park Co., Yellowstone National Park, 24-Jul-2017 | | | |
| 814 | NVG-17068C03 | *Oarisma poweshiek* | | | USA: MN, Pipestone Co., NE of Holland, 11-Jul-1986 | CSUC | | CSU_ENT1025108 |
| 815 | NVG-6395 | *Oarisma garita garita* | | F | USA: CO, Grand Co., 6 air mi E Kremmling, 5-Jul-2016 | | | |
| 816 | NVG-17068B09 | *Oarisma edwardsii* | | | USA: NM, Catron Co., 7.2 miles north of NM 12 on USFS 11 at Toro Canyon, 31-Jul-1995 | CSUC | | CSU_ENT1026918 |
| 817 | NVG-8381 | *Oarisma aurantiaca* | | M | USA: TX, Blanco Co., Middle Creek Rd. nr. USH290, 31-Mar-2017 | | | |
| 818 | NVG-9127 | *Oarisma minima* | | F | USA: FL, Hernando Co., Hill 'N Dale, 8-Jun-2017 | | | |
| 819 | NVG-17068F04 | *Panoquina errans* | | F | Mexico: Sonora, Guaymas, 2 miles West San Carlos, 23-Mar-2004 | CSUC | | CSU_ENT1027432 |
| 820 | NVG-4687 | *Panoquina panoquinoides* | | | USA: FL, Levy Co., Yankeetown, 26-Sep-2015 | | | |
| 821 | NVG-4155 | *Panoquina panoquin* | | F | USA: TX, Jefferson Co., S of Sabine Pass, 18-Jul-2015 | | | |
| 822 | NVG-14111H05 | *Panoquina evansi* | | M | USA: TX, Cameron Co., La Feria, 5-Nov-2014 | WDempwolf | | |
| 823 | NVG-16107D05 | *Panoquina hecebolus* | | M | USA: TX, Hidalgo Co., Rio Rico Rd., 21-Oct-2013 | WDempwolf | | |
| 824 | NVG-5158 | *Panoquina ocola ocola* | | F | USA: TX, Hidalgo Co., LaGrulla, 15-Nov-2015 | | | |
| 825 | NVG-18037G09 | *Panoquina lucas* | | F | USA: TX, Hidalgo Co., 24-Oct-2004 | | | |
| 826 | NVG-4591 | *Calpodes ethlius* | | F | USA: TX, Cameron Co., La Feria, 9-Sep-2015 | | | |
| 827 | NVG-18112G04 | *Calpodes esperi esperi* | | | Guyana, 30-Nov-5-Dec-2000 | USNM | | |
| 828 | NVG-17109H02 | *Perichares adela* | | M | Mexico: Hidalgo, 30-Mar-1981 | LACM | | |
| 829 | NVG-2118 | *Agathymus gentryi* | | | USA: AZ, Pima Co. | | | |
| 830 | NVG-312 | *Agathymus baueri baueri* | | | USA: AZ, Yavapai Co., Oct-2004 | | | |
| 831 | NVG-33 | *Agathymus aryxna* | | | USA: AZ, Cochise Co., Sep-2003 | | | |
| 832 | NVG-25 | *Agathymus evansi* | | | USA: AZ, Cochise Co., Sep-2003 | | | |
| 833 | NVG-19 | *Agathymus polingi* | | | USA: AZ, Cochise Co., S of Benson, 20-Aug-2003 | | | |

| | | | | | | | | |
|---|---|---|---|---|---|---|---|---|
| 834 | NVG-214 | *Agathymus neumoegeni neumoegeni* | | | USA: AZ, Coconino Co., SR260, 0.8mi E county line, 15-Sep-2004 | | | |
| 835 | NVG-199 | *Agathymus chisosensis* | | F | USA: TX, Brewster Co., Big Bend National Park, 11-Sep-2004 | | | |
| 836 | NVG-29 | *Agathymus alliae alliae* | | M | USA: AZ, Navajo Co., 17-Sep-2003 | | | |
| 837 | NVG-202 | *Agathymus stephensi* | | | USA: CA, San Diego Co., Sep-2004 | | | |
| 838 | NVG-256 | *Agathymus mariae mariae* | | | USA: TX, Hudspeth Co., Sep-2004 | | | |
| 839 | NVG-18022C01 | *Agathymus gilberti* | HT | F | USA: TX, Kinney Co., 14 mi N Brackettville, 22-Oct-1961 | AMNH | F.H.R.No.19,774 | M1963 |
| 840 | NVG-94 | *Agathymus estelleae valverdiensis* | | | USA: TX, Uvalde Co., Mar-2004 | | | |
| 841 | NVG-18024B10 | *Stallingsia maculosus* | PT | M | USA: TX, Hidalgo Co., Sullivan City, 4-Apr-1953 | AMNH | | |
| 842 | NVG-1461 | *Megathymus streckeri streckeri* | | M | USA: AZ, Apache Co., SE of Holbrook, 19-May-2013 | | | |
| 843 | NVG-1536 | *Megathymus cofaqui cofaqui* | | F | USA: GA, Burke Co., NW of Girard, 2-Aug-2013 | | | |
| 844 | NVG-1185 | *Megathymus yuccae yuccae* | | M | USA: SC, Aiken Co., W of Jackson, 25-Feb-2013 | | | |
| 845 | NVG-1528 | *Megathymus ursus ursus* | | M | USA: AZ, Pima Co., Rincon Mtns., ecl. on 6-Aug-2013 | | | |
| out | NVG-18032H12 | *Pseudothyris sepulchralis* | | | USA: TX, Travis Co., Austin, Zilker park, 12-Mar-2000 | | | |

| Collection abbreviations | |
|---|---|
| AMNH | American Museum of Natural History, New York, NY, USA |
| ANSP | Academy of Natural Sciences of Drexel University, Philadelphia, PA, USA |
| BMUW | Burke Museum of Natural History and Culture, Seattle, WA, USA |
| CAS | California Academy of Sciences, San Francisco, CA, USA |
| CSUC | Colorado State University Collection, Fort Collins, CO, USA |
| FMNH | Field Museum of Natural History, Chicago, FL, USA |
| JAScott | research collection of James A. Scott |
| JAShuey | research collection of John A. Shuey |
| JPBrock | research collection of Jim P. Brock |
| JMcDermott | research collection of James McDermott |
| JSCarter | research collection of Jack S. Carter |
| LACM | Los Angeles County Museum of Natural History, Los Angeles, CA, USA |
| MWalker | research collection of Mark Walker |
| MGCL | McGuire Center for Lepidoptera and Biodiversity, Gainesville, FL, USA |
| HPavulaan | research collection of Harry Pavulaan |
| TAMU | Texas A&M University Insect Collection, College Station, TX, USA |
| TLS | Research collection of Texas Lepidoptera Survey, Houston, TX, USA |
| TMMC | University of Texas Biodiversity Center, Austin, TX, USA |
| UCDC | Bohart Museum of Entomology, University of California, Davis, CA, USA |
| USNM | National Museum of Natural History, Smithsonian Institution, Washington, DC, USA |
| WDempwolf | research collection of William R. Dempwolf |

**HT**, holotype; **NT**, neotype; **PLT**, paralectotype; **PT**, paratype; **ST**, syntype

### References:

**Cong, Q., J. Zhang, J. Shen, and N. V. Grishin**. **2019**. Fifty new genera of Hesperiidae (Lepidoptera). Insecta Mundi 0731: 1–56.

**Pelham, J. P. 2019**. Catalogue of the Butterflies of the United States and Canada. Revised 7 Oct 2019. Accessed 8 Oct 2019. http://www.butterfliesofamerica.com/US-Can-Cat.htm.

**Zhang, J., Q. Cong, J. Shen, P. A. Opler, and N. V. Grishin**. **2019**. Changes to North American butterfly names. The Taxonomic Report of the International Lepidoptera Survey 8(2): 1–11.
